# Identification of novel prognostic targets in acute kidney injury using bioinformatics and next generation sequencing data analysis

**DOI:** 10.1101/2024.03.17.585400

**Authors:** Basavaraj Vastrad, Chanabasayya Vastrad

## Abstract

Acute kidney injury (AKI) is a type of renal disease occurs frequently in hospitalized patients, which may cause abnormal renal function and structure with increase in serum creatinine level with or without reduced urine output. With the incidence of AKI is increasing. However, the molecular mechanisms of AKI have not been elucidated. It is significant to further explore the molecular mechanisms of AKI. We downloaded the GSE139061 next generation sequancing (NGS) dataset from the Gene Expression Omnibus (GEO) database. Limma R bioconductor package was used to screen the differentially expressed genes (DEGs). Then, the enrichment analysis of DEGs in Gene Ontology (GO) function and REACTOME pathways was analyzed by g:Profiler. Next, the protein-protein interaction (PPI) network and modules was constructed and analysed, and the hub genes were identified. Next, the miRNA-hub gene regulatory network and TF-hub gene regulatory network were built. We also validated the identified hub genes via receiver operating characteristic (ROC) curve analysis. Overall, 956 DEGs were identified, including 478 up regulated and 478 down regulated genes. The enrichment functions and pathways of DEGs involve primary metabolic process, small molecule metabolic process, striated muscle contraction and metabolism. Topological analysis of the PPI network and module revealed that hub genes, including PPP1CC, RPS2, MDFI, BMI1, RPL23A, VCAM1, ALB, SNCA, DPP4 and RPL26L1, might be involved in the development of AKI. miRNA-hub gene and TF-hub gene regulatory networks revealed that miRNAs and TFs including hsa-mir-510-3p, hsa-mir-6086 and mir-146a-5p, MAX and PAX2, might be involved in the development of AKI. Various known and newtherapeutic targets were obtained via network analysis. The results of the current investigation might be beneficial for the diagnosis and treatment of AKI.

## Introduction

Acute kidney injury (AKI) is a sudden episode of kidney failure or kidney damage characterized by rapid decline in glomerular filtration rate and increase in serum creatinine level with or without reduced urine output [1]. Plentiful widespread kidney diseases complicated with AKI, which badly increases morbidity and mortality [2]. PAH is a reported prevalence of 5.0–7.5% of hospitalized patients and in 50–60% of critically ill patients [3]. There are substantial causative factors of AKI, like cardiovascular events [4], older adults [5], mechanical ventilator [6], nephrotoxic agents [7], vasopressors [8], hypertension [9], hypercholesterolemia [10], liver disease [11], hypoalbuminemia [12], sepsis [13], renal ischemia [14], diabetes mellitus [15], obesity [16], anemia [17] and chronic kidney disease [18]. Studies have revealed that the progression of AKI is related to genetic factors [19]. Rhabdomyolysis and renal replacement therapy is the major therapeutic option for AKI treatment [20]. Noradrenaline and furosemide are alternative treatment options [21–22]. However, the efficacy of these treatment strategies is still limited. Hence, developing new diagnosis biomarkers reflecting the damage of kidney is appealing for the timely diagnoses and therapies of AKI.

The recent next generation sequencing (NGS) analysis of specimens from sufferers and normal individuals enables us to investigate various diseases at the transcriptomic level [23–24]. Recently, many specific genes and signaling pathways have been discovered to participate in the progression of AKI. For instance, it was reported that the expression level of MCP-1 [25], BCL2, SERPINA4, SERPINA5, and SIK3 [26], CYBA [27], ATF3 [28] and TAUT [29] were considerably altered in AKI. PI3K/Akt/Nrf2 signaling pathway [30], NF-κB and JAK2/STAT3 signaling pathway [31], IL-18 receptor signaling pathway [32], TLR2, TLR4 and the MYD88 signaling pathway [33] and CXCL16/ROCK1 signaling pathway [34] can offer novel enlightenment for the treatment of AKI. These finding suggested the key roles of some function genes in AKI progression. However, the diagnostic value of various genes has not been investigated in AKI.

In this investigation, we aimed to identify novel diagnostic genes and therapeutic targets for AKI based on bioinformatics and NGS data analysis. We analyzed NGS NCBI Gene Expression Omnibus (GEO) [https://www.ncbi.nlm.nih.gov/geo/] [35] dataset (GSE139061) to determine DEGs between AKI specimens and normal control specimens. By analyzing the gene ontology (GO) and REACTOME pathway enrichment analysis, along with the construction of protein-protein interaction (PPI) network and module analysis, we selected hub genes. After evaluating the clinical prognosis of these hub genes and their miRNA-hub gene regulatory network and TF-hub gene regulatory network are drawn, we further validated these hub genes by receiver operating characteristic (ROC) curve analysis. Our findings provided novel key genes involved in the advancement of AKI.

## Materials and Methods

### Next generation sequencing data source

NGS dataset GSE139061 (Illumina HiSeq 4000 (Homo sapiens) was downloaded from the GEO database. The GSE139061 dataset consisted of 39 glomerular tissue samples from AKI patients and 9 glomerular tissue samples from normal control human kidneys.

### Identification of DEGs

“Limma” R bioconductor package [36] was utilized to identify the DEGs between the glomerular tissues of AKI patients and normal control human kidneys. True P-values were corrected using the Benjamini-Hochberg method [37]. A p-value <0.05, |log FC (fold change)| > 2.031 for up regulated genes and |log FC (fold change)| < - 1.855 for down regulated genes were considered statistically significant. The “ggplot2” package of R software was used to construct a volcano plot of the DEGs, and “gplot” package of R software was used to establish a heatmap of the DEGs.

### GO and pathway enrichment analyses of DEGs

Gene Ontology (GO) (http://www.geneontology.org) [38] and REACTOME (https://reactome.org/) [39] pathway enrichment analyses of AKI DEGs were performed using g:Profiler (http://biit.cs.ut.ee/gprofiler/) [40]. GO terms of biological processes (BP), cellular components (CC), and molecular functions (MF) linked with a p-value <0.05 were considered to be significantly enriched.

### Construction of the PPI network and module analysis

PPI network among all DEGs was established based on an online tool Human Integrated Protein-Protein Interaction rEference (HIPIE) (https://cn.string-db.org/) [41] and then software Cytoscape (version 3.10.1) (http://www.cytoscape.org/) [42] was employed to adjust and visualize PPI networks. To screen the hub genes that might be associated in AKI, we applied the Network Analyzer plugin, using various topological parameters such as degree [43], betweenness [44], stress [45] and closeness [46]. The PEWCC [47] Cytoscape software plug in was used to create modules in the PPI network of AKI.

### Construction of the miRNA-hub gene regulatory network

Hub gene information of miRNAs was collected from of miRNet database (https://www.mirnet.ca/) [48], which is an experimentally validated miRNA-hub gene interactions database. The interaction of miRNAs and hub genes in AKI was used to construct the miRNAs - hub genes regulated network. Cytoscape software [42] was used to visualize the network.

### Construction of the TF-hub gene regulatory network

Hub gene information of TFs was collected from of NetworkAnalyst database (https://www.networkanalyst.ca/) [49], which is an experimentally validated TF-hub gene interactions database. The interaction of TFs and hub genes in AKI was used to construct the TFs - hub genes regulated network. Cytoscape software [42] was used to visualize the network.

### Receiver operating characteristic curve (ROC) analysis

ROC was plotted using the pROC package of R software [50] to assess the performance of hub genes distinguishing between AKI and normal individuals. The diagnostic value of the hub genes and the integrated diagnostic model was validated with an external expression dataset. The area under the curve (AUC), sensitivity, and specificity were determined. Those hub genes with AUC values > 0.60 in AKI were considered with diagnostic values, and those with AUC values > 0.80 were considered with high diagnostic values.

## Results

### Identification of DEGs

In this investigation, we used NGS dataset with a total sample size 48, consisting of 39 AKI samples and 9 normal control samples. All samples were combined for batch correction. During the screening of DEGs, the cutoff value was set as |log FC (fold change)| > 2.031 for up regulated genes and |log FC (fold change)| < -1.855 for down regulated genes-, and the p value was < 0.05. Finally, 956 DEGs were obtained, including 478 up regulated genes and 478 down regulated genes. According to the DEGs, we constructed the volcano plot (Fig. 1) and heat map (Fig. 2). Volcano plot can show the distribution of DEGs and heat map can determine the similarity of expression of genes. The up regulated genes and down-regulated genes are listed in Table 1.

**Fig. 1.**
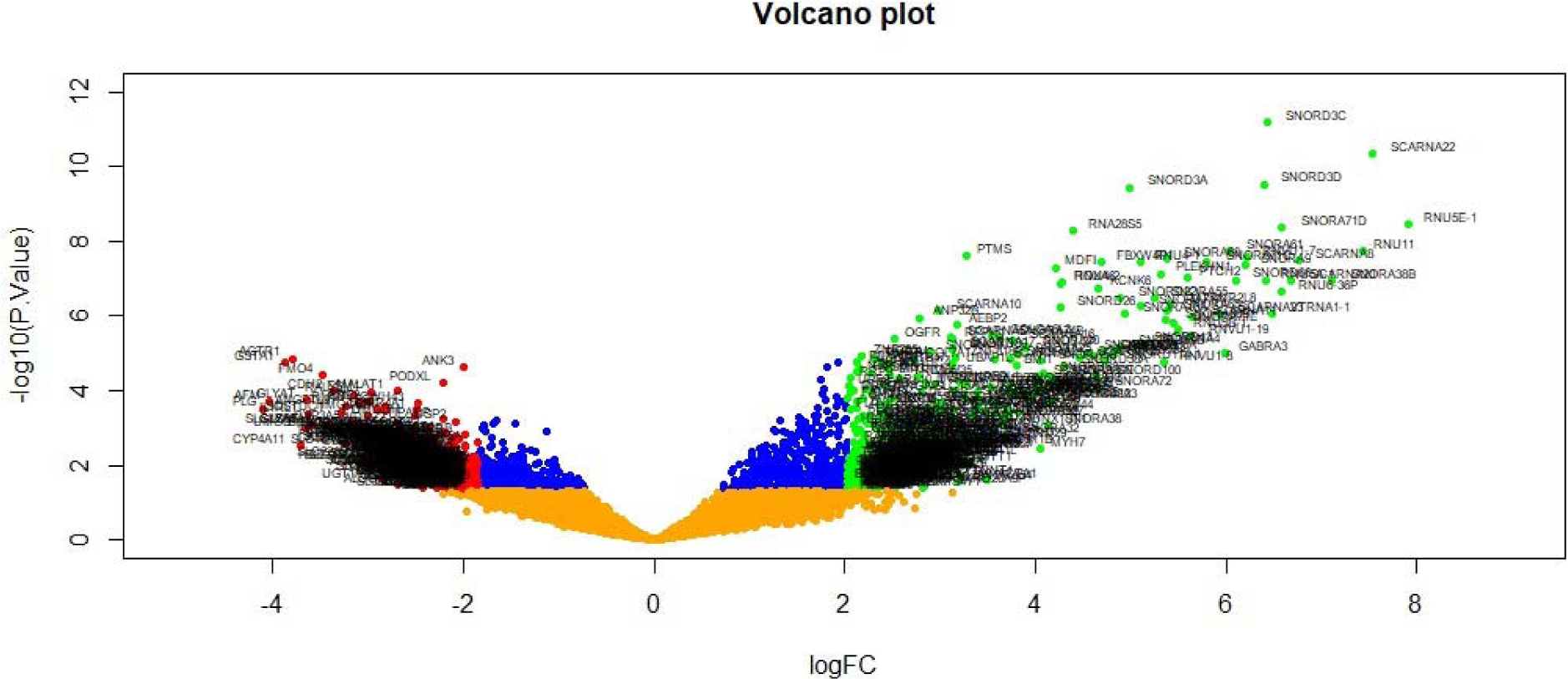
Volcano plot of differentially expressed genes. Genes with a significant change of more than two-fold were selected. Green dot represented up regulated significant genes and red dot represented down regulated significant genes.

**Fig. 2.**
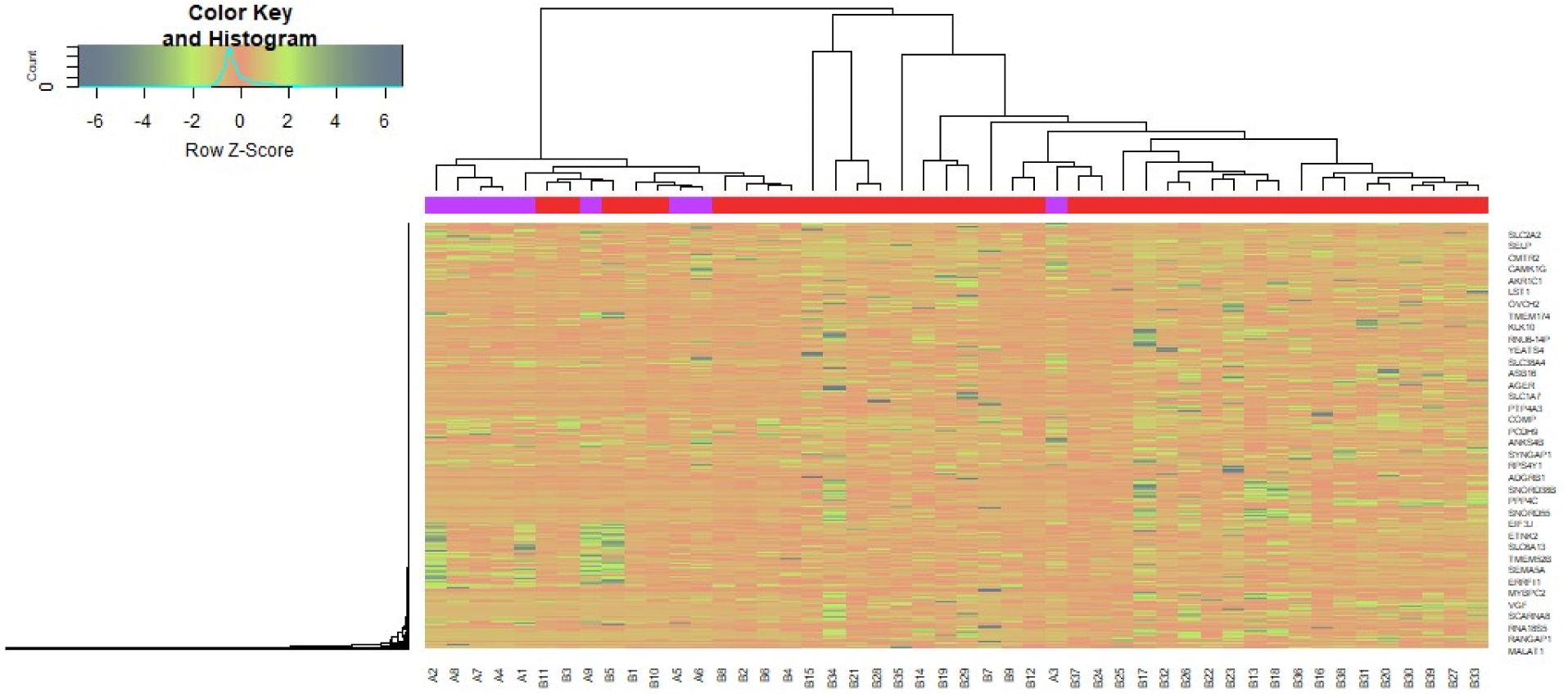
Heat map of differentially expressed genes. Legend on the top left indicate log fold change of genes. (A1 – A9 = Normal control samples; B1 – B 39 = AKI samples)

**Table 1.**
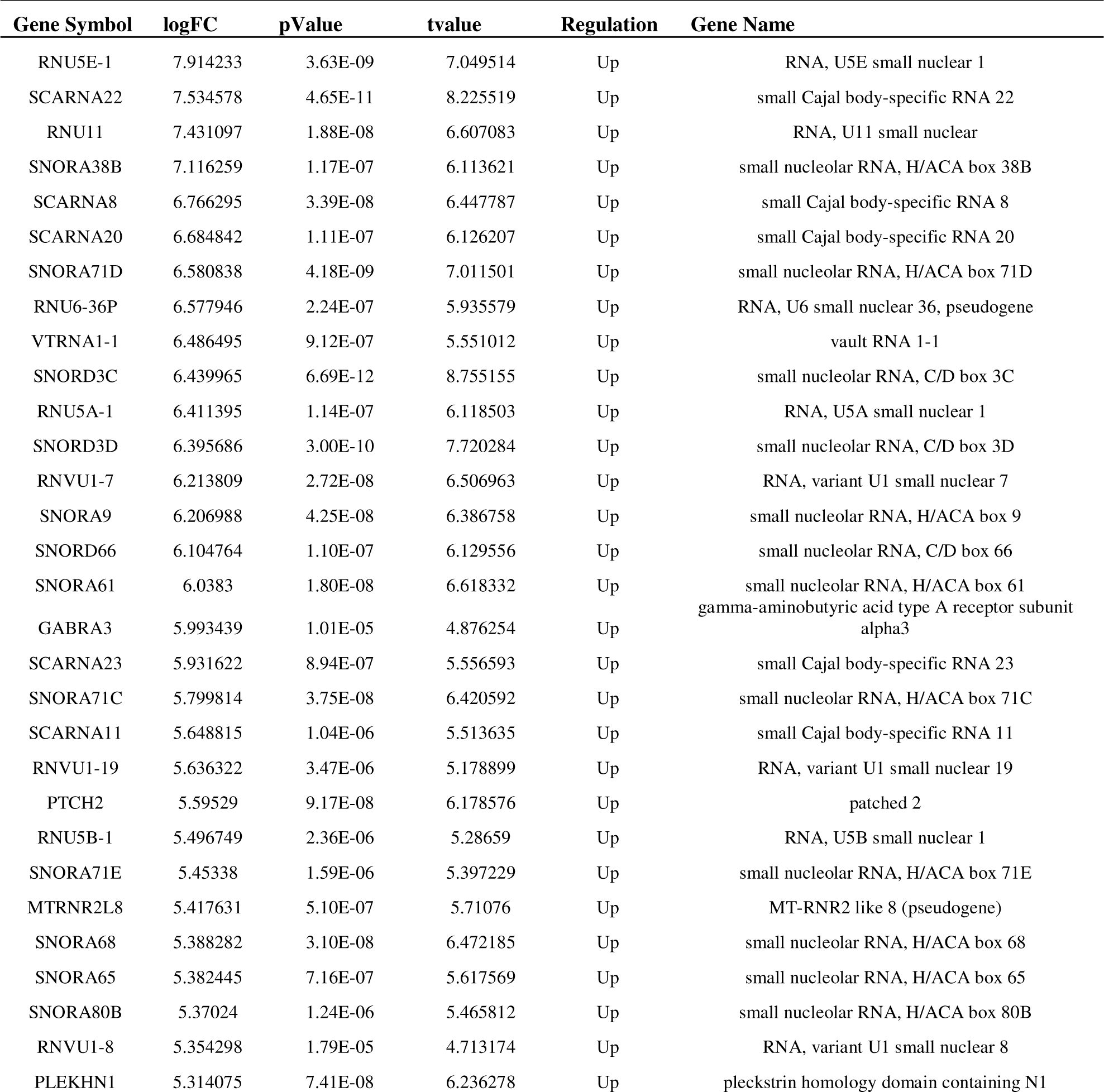
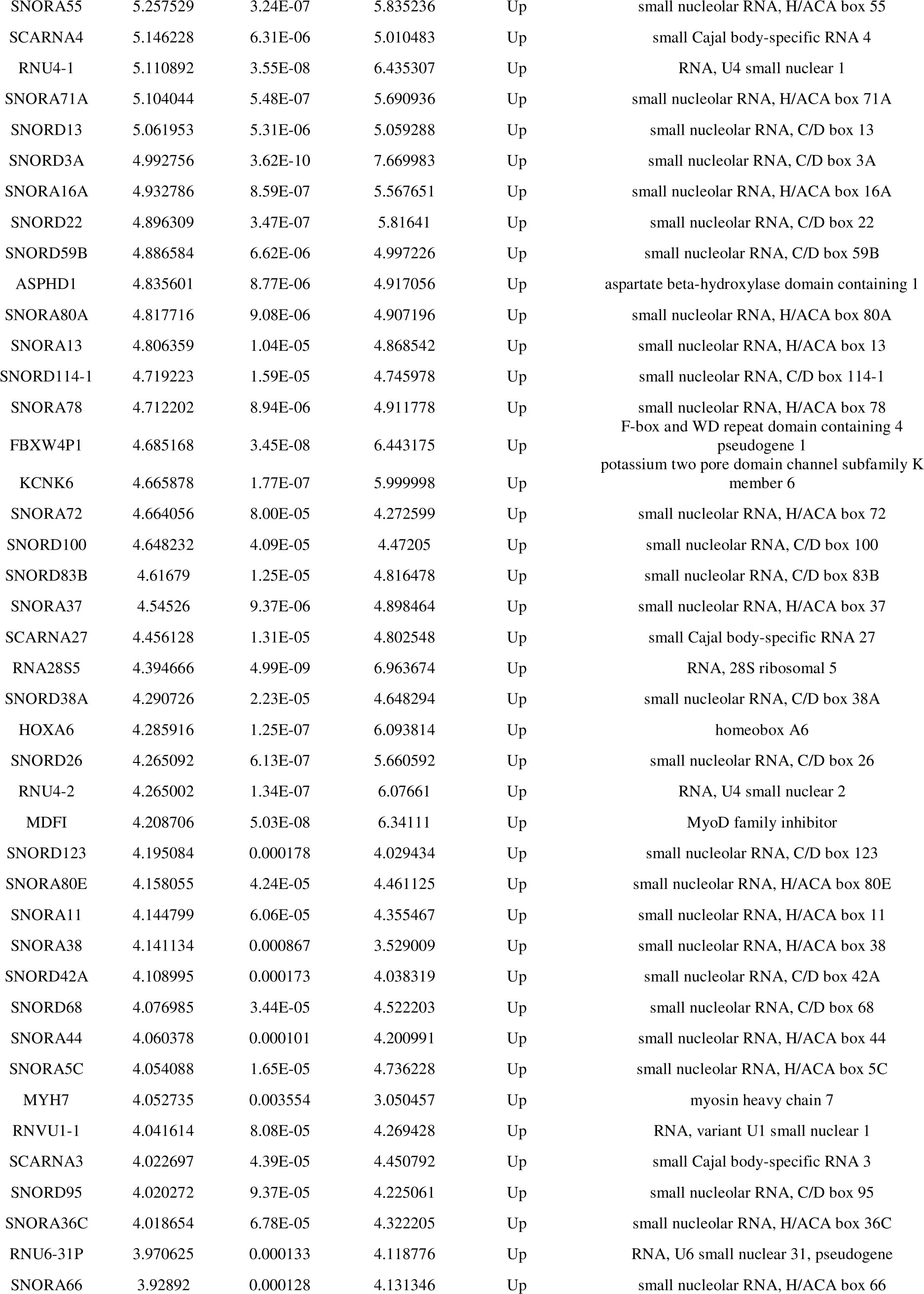
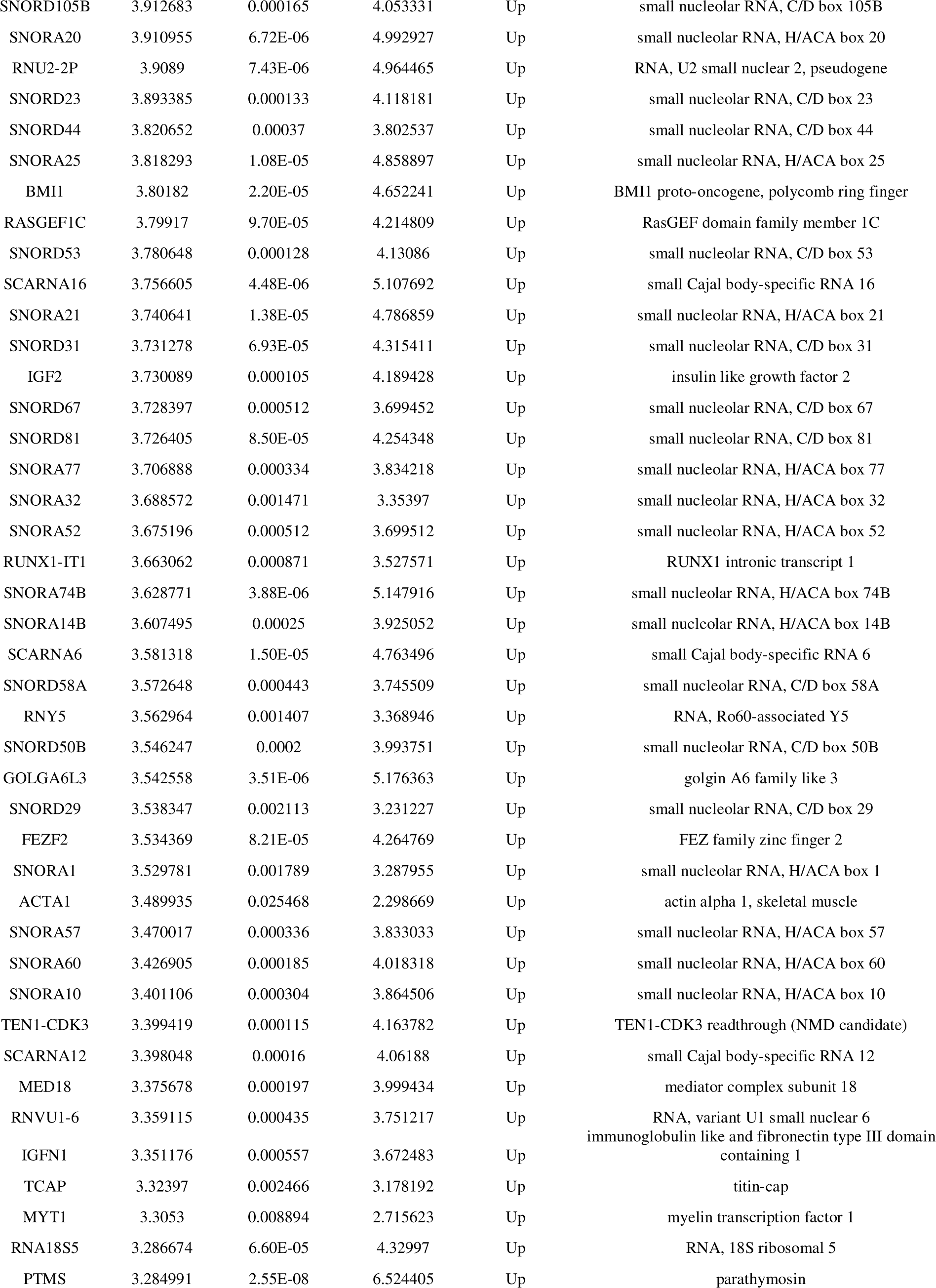
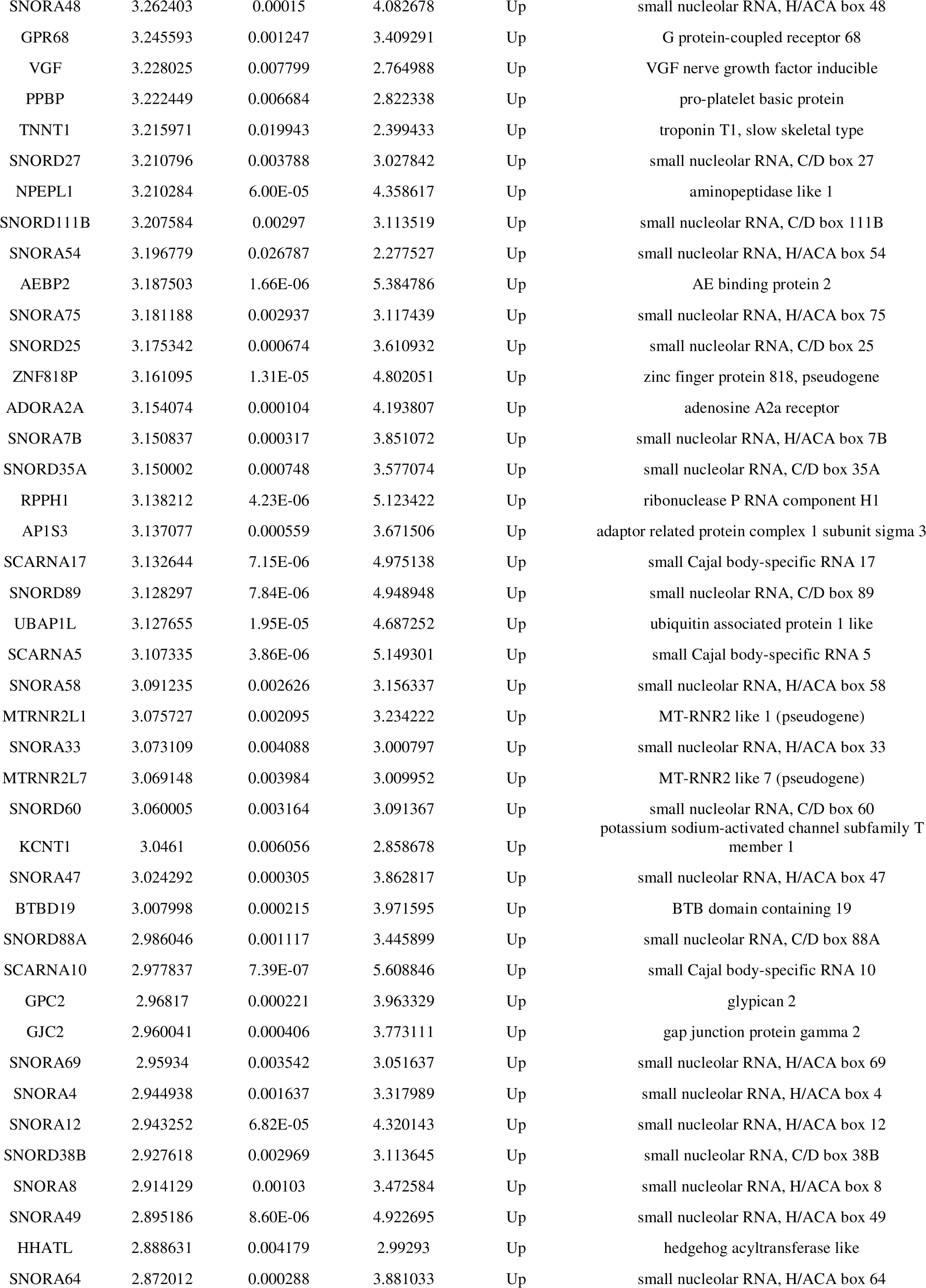
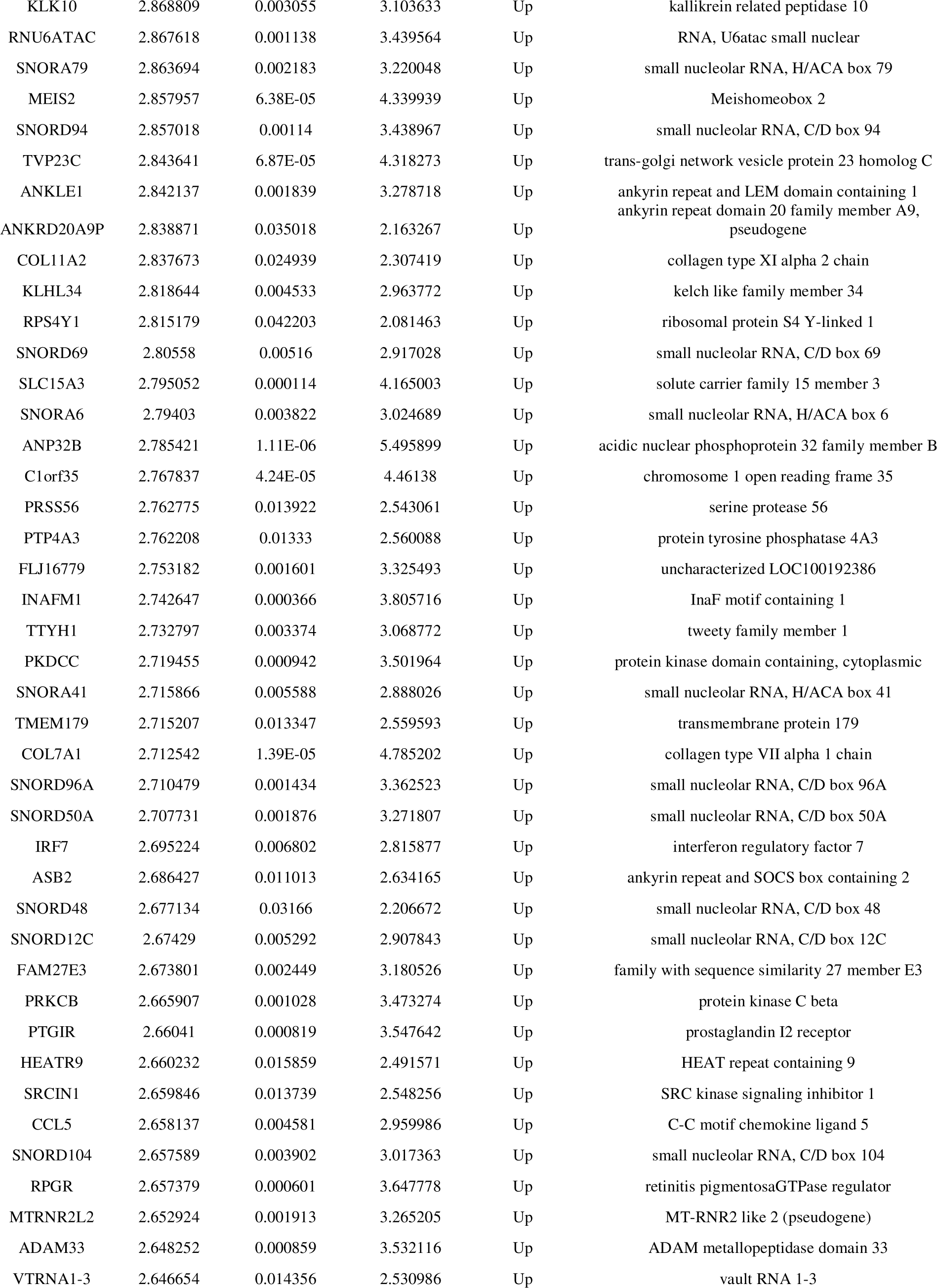
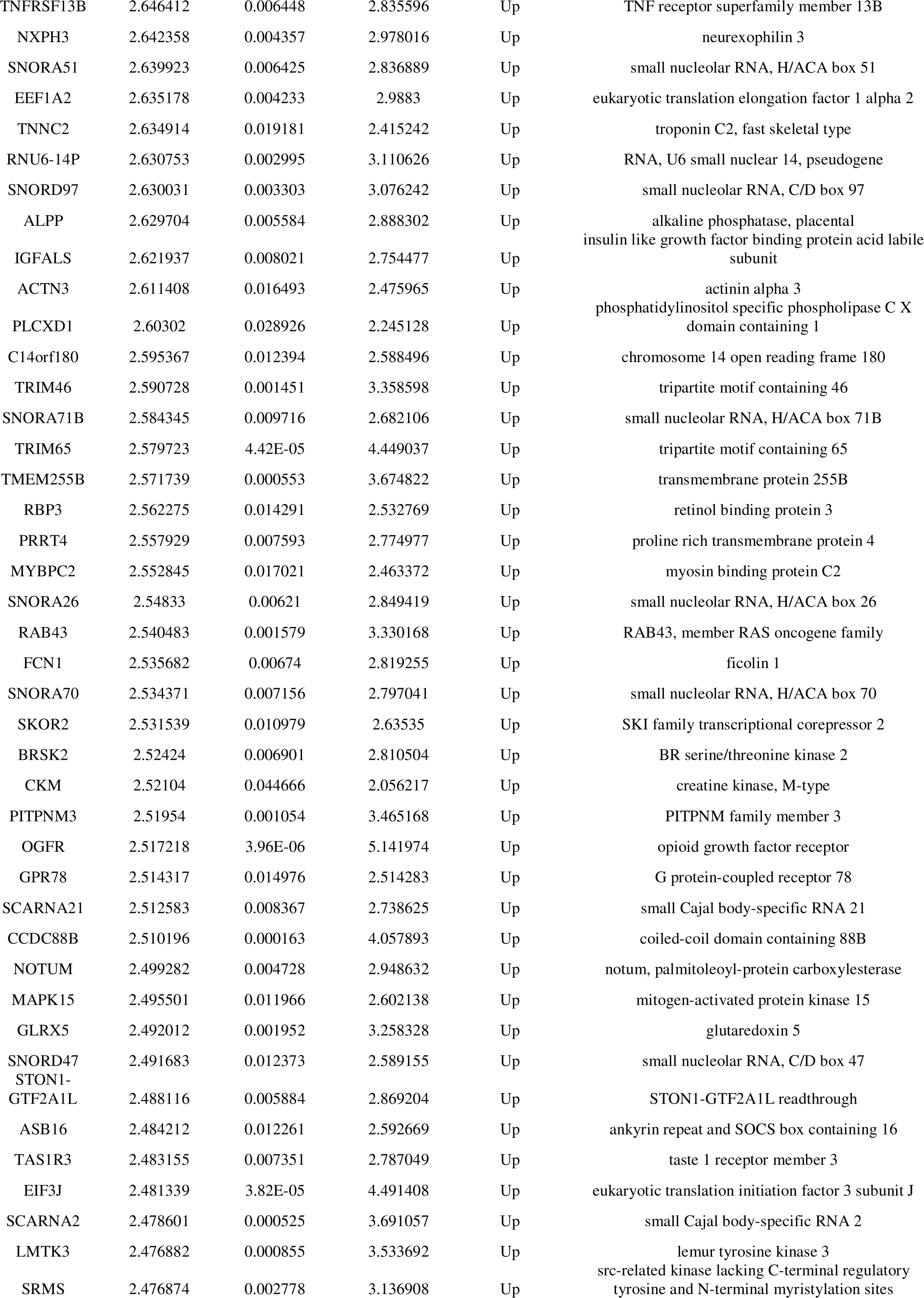
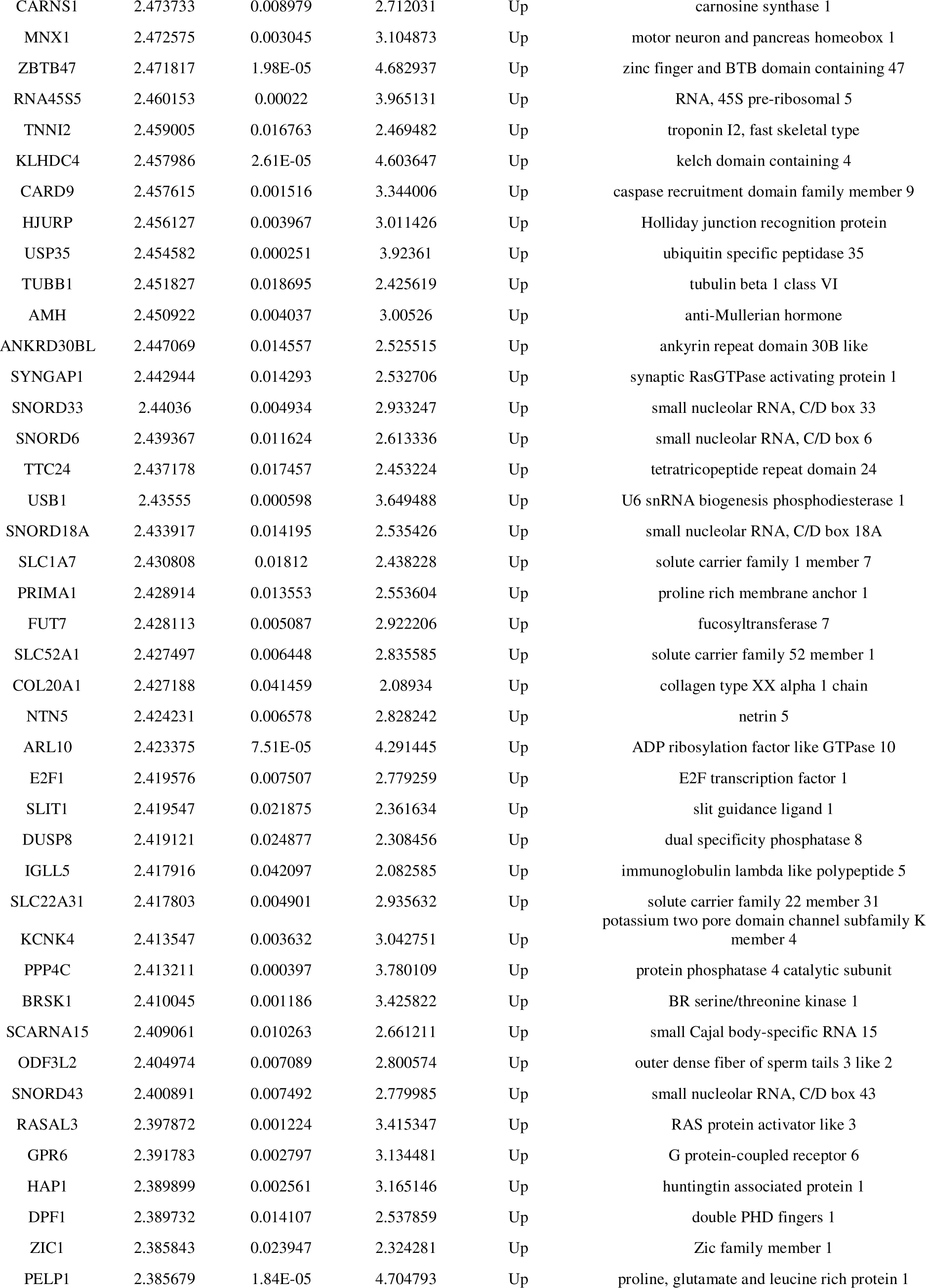
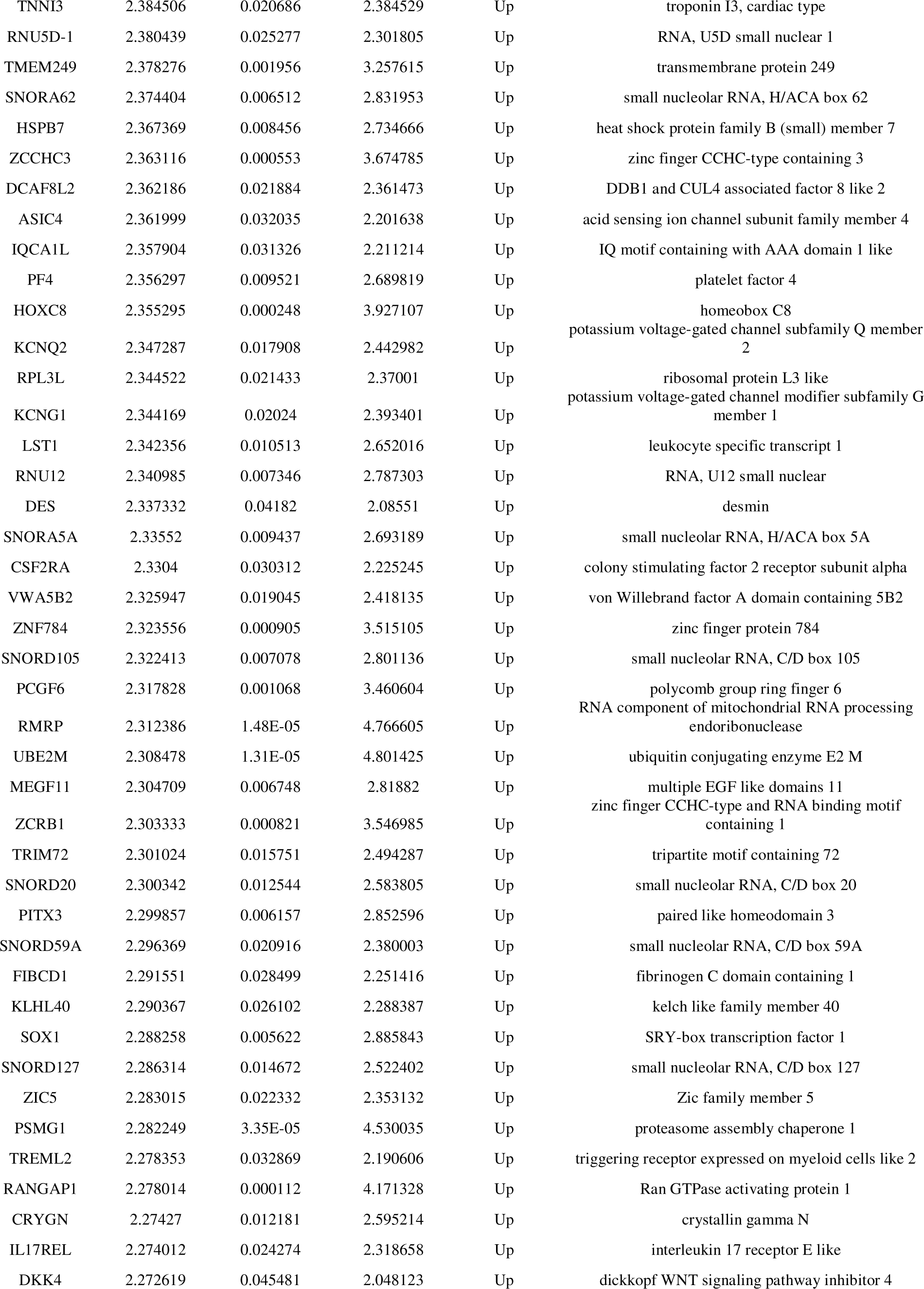
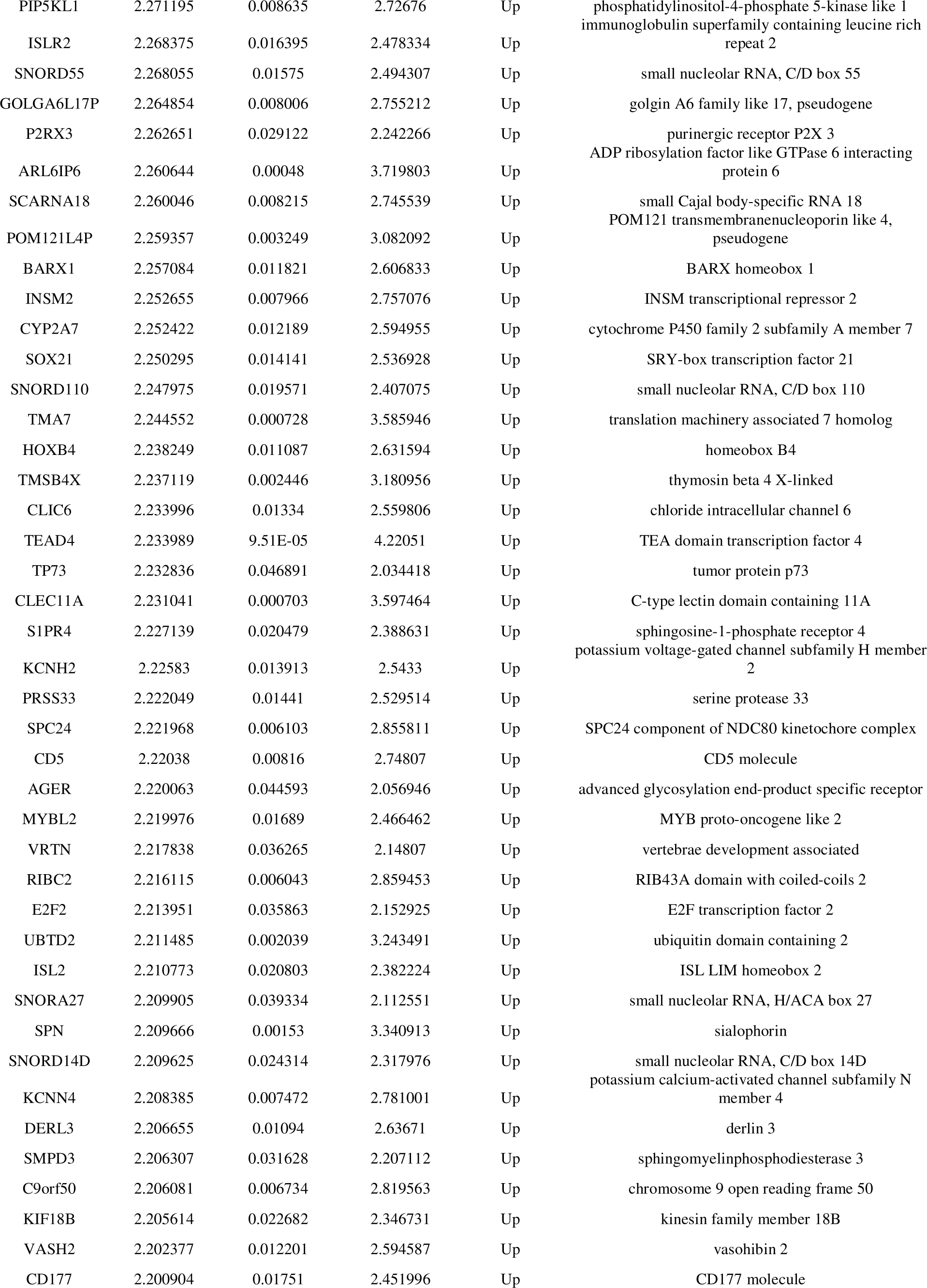
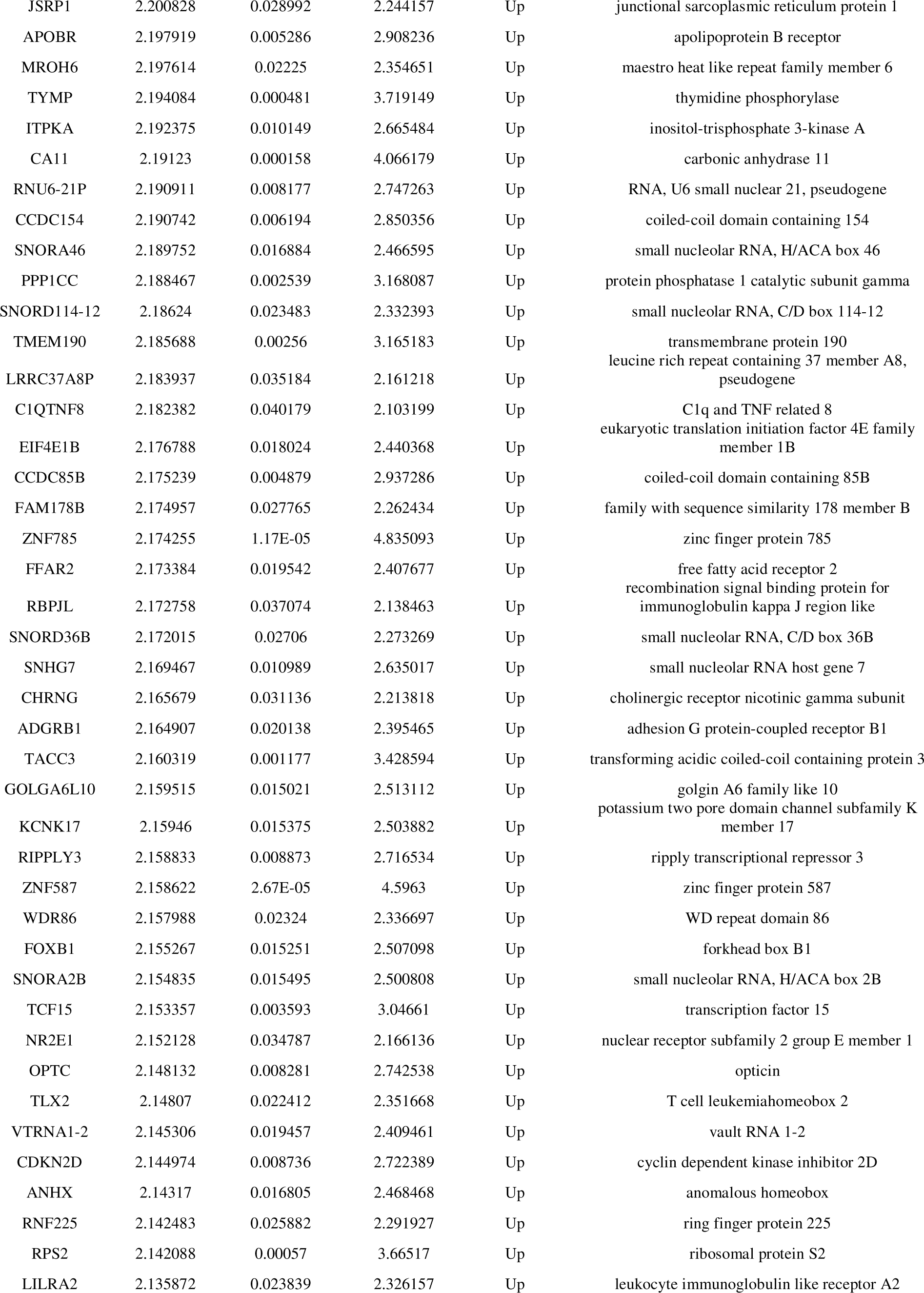
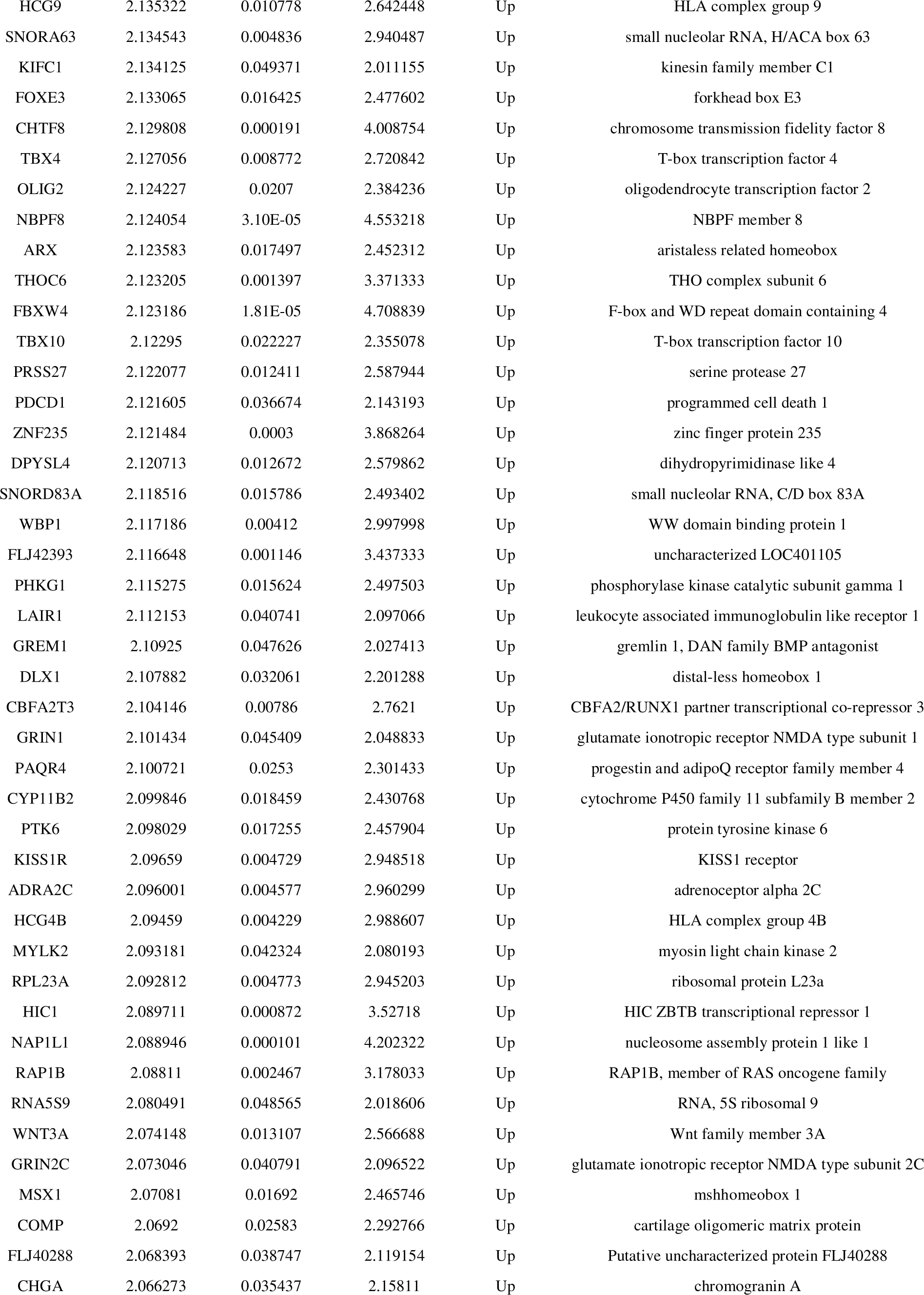
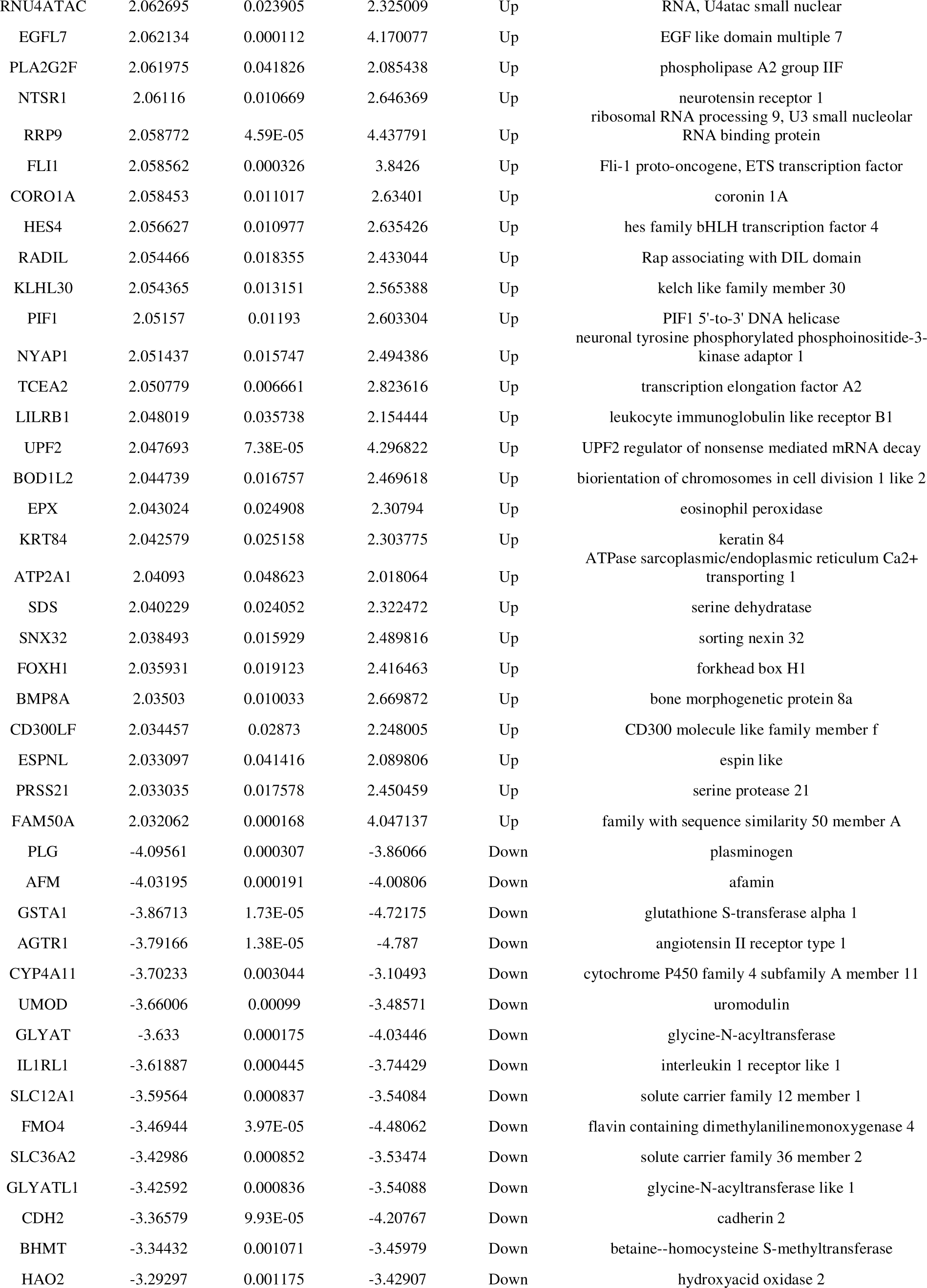
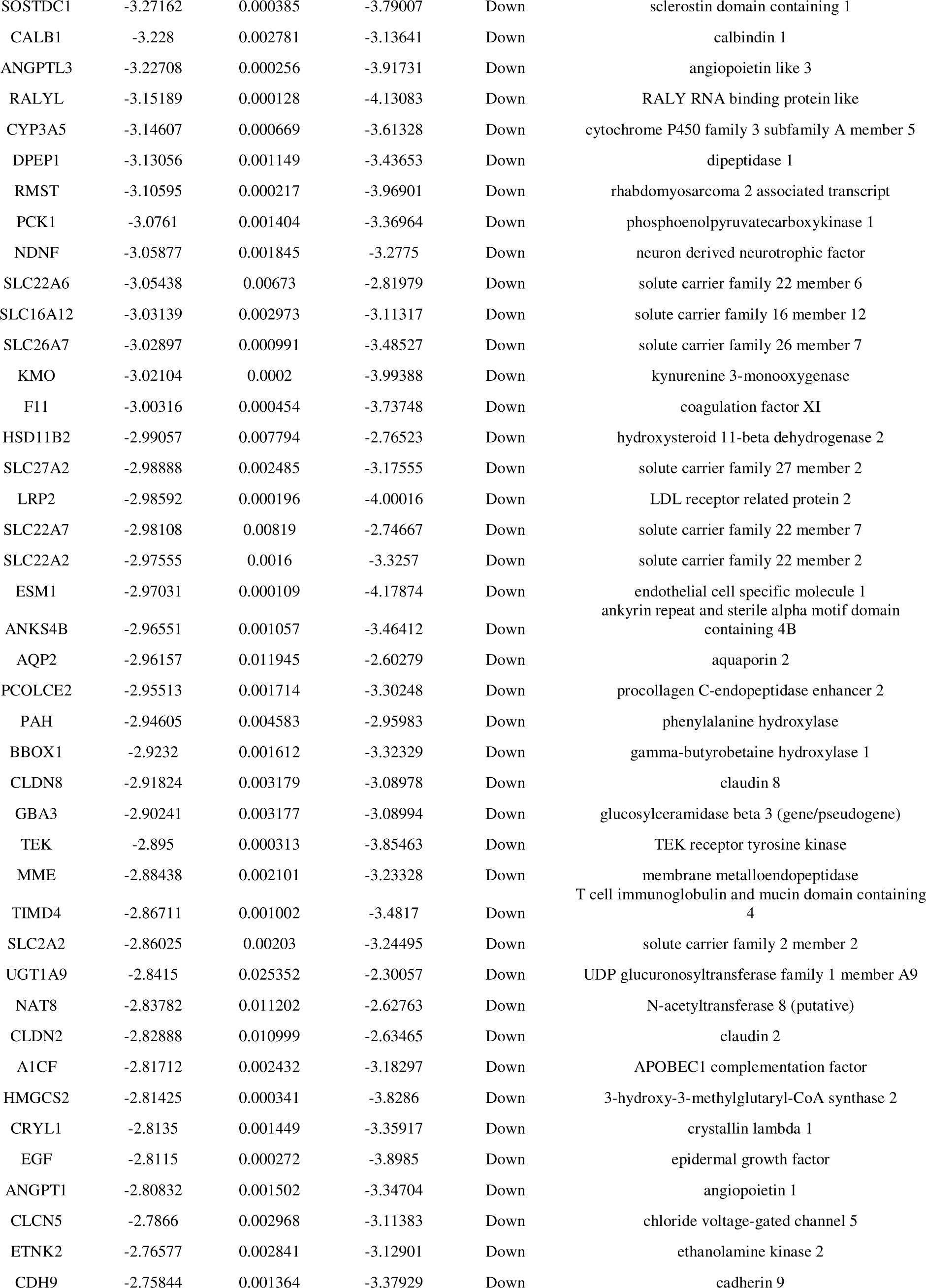
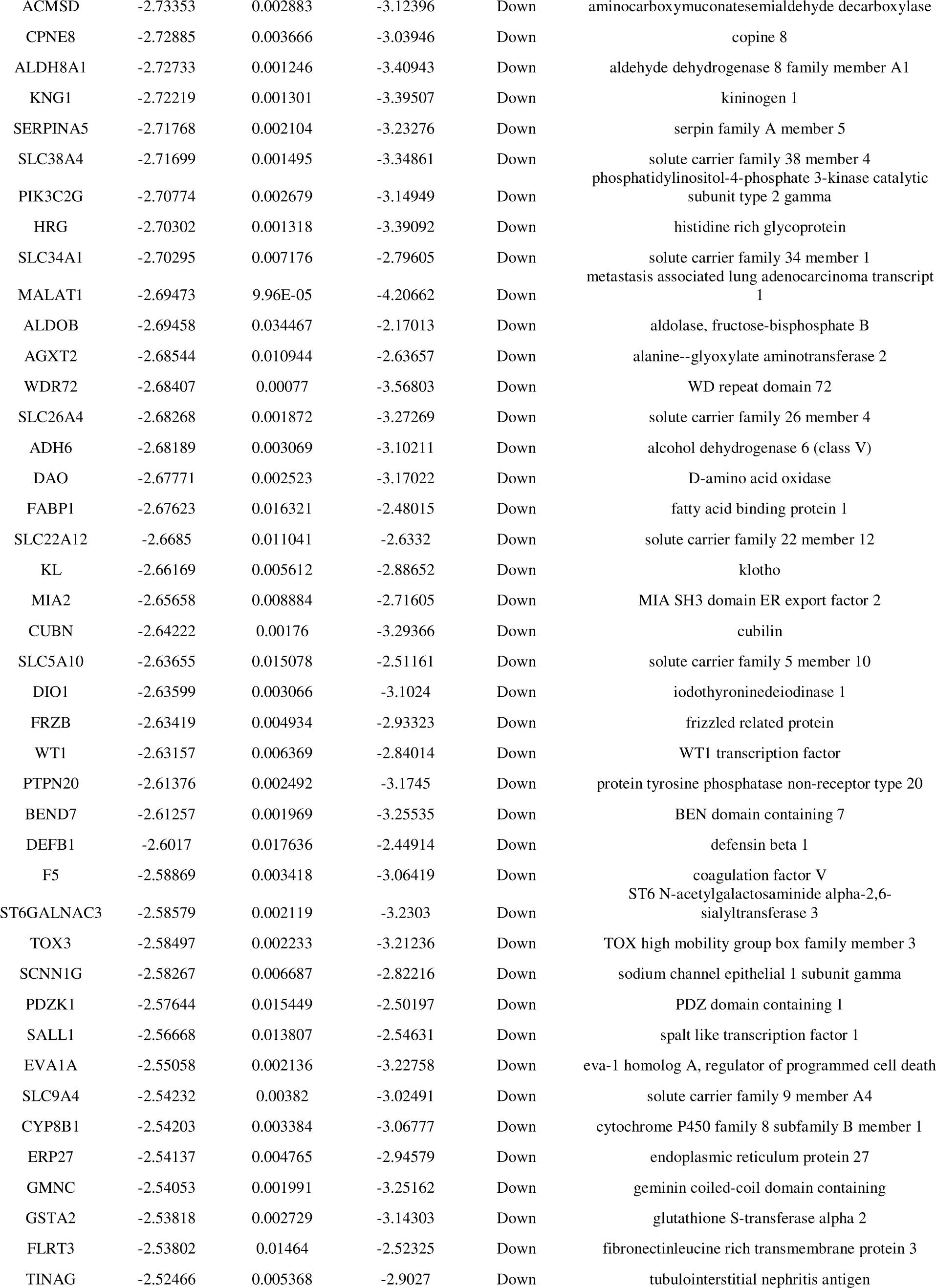
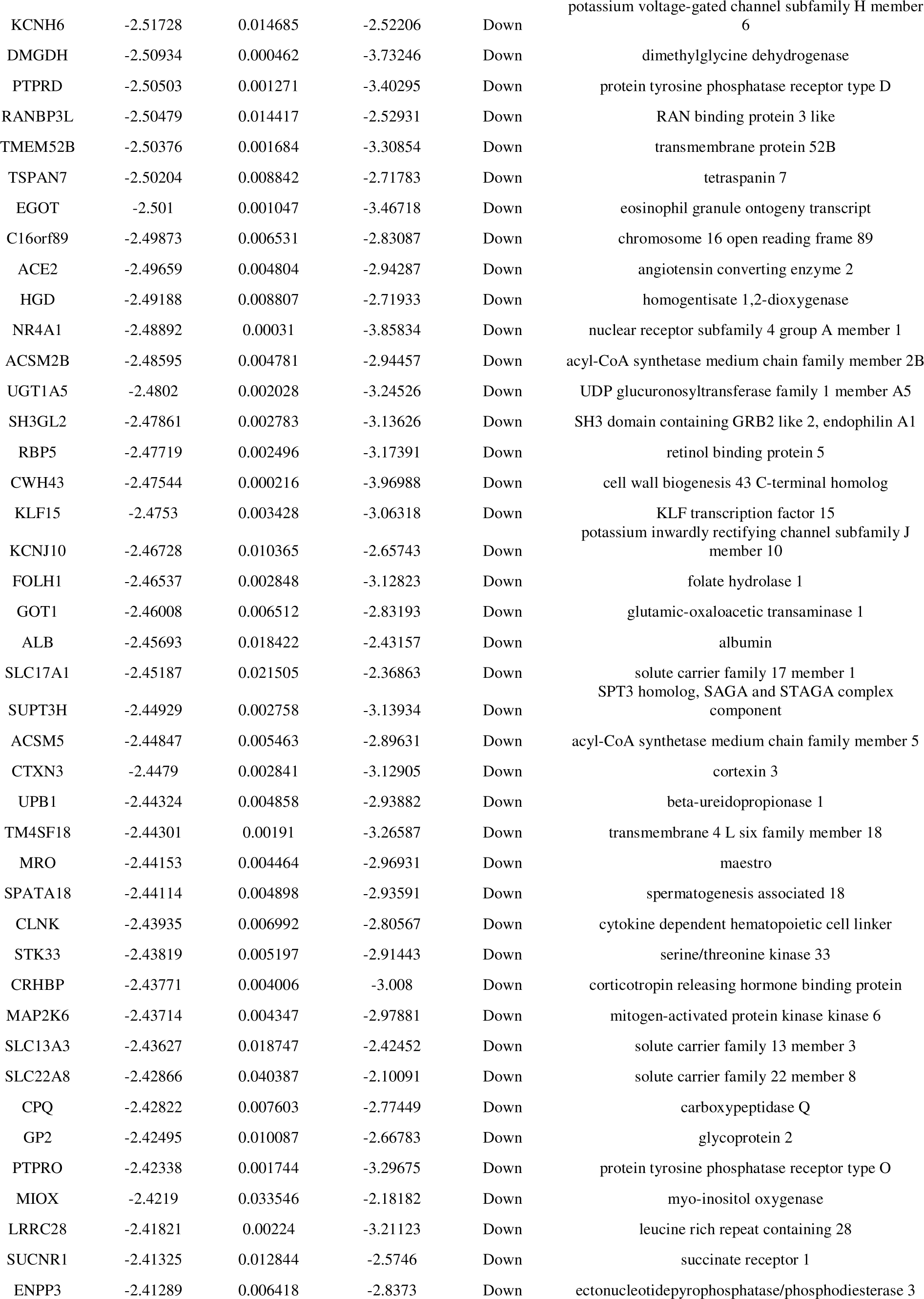
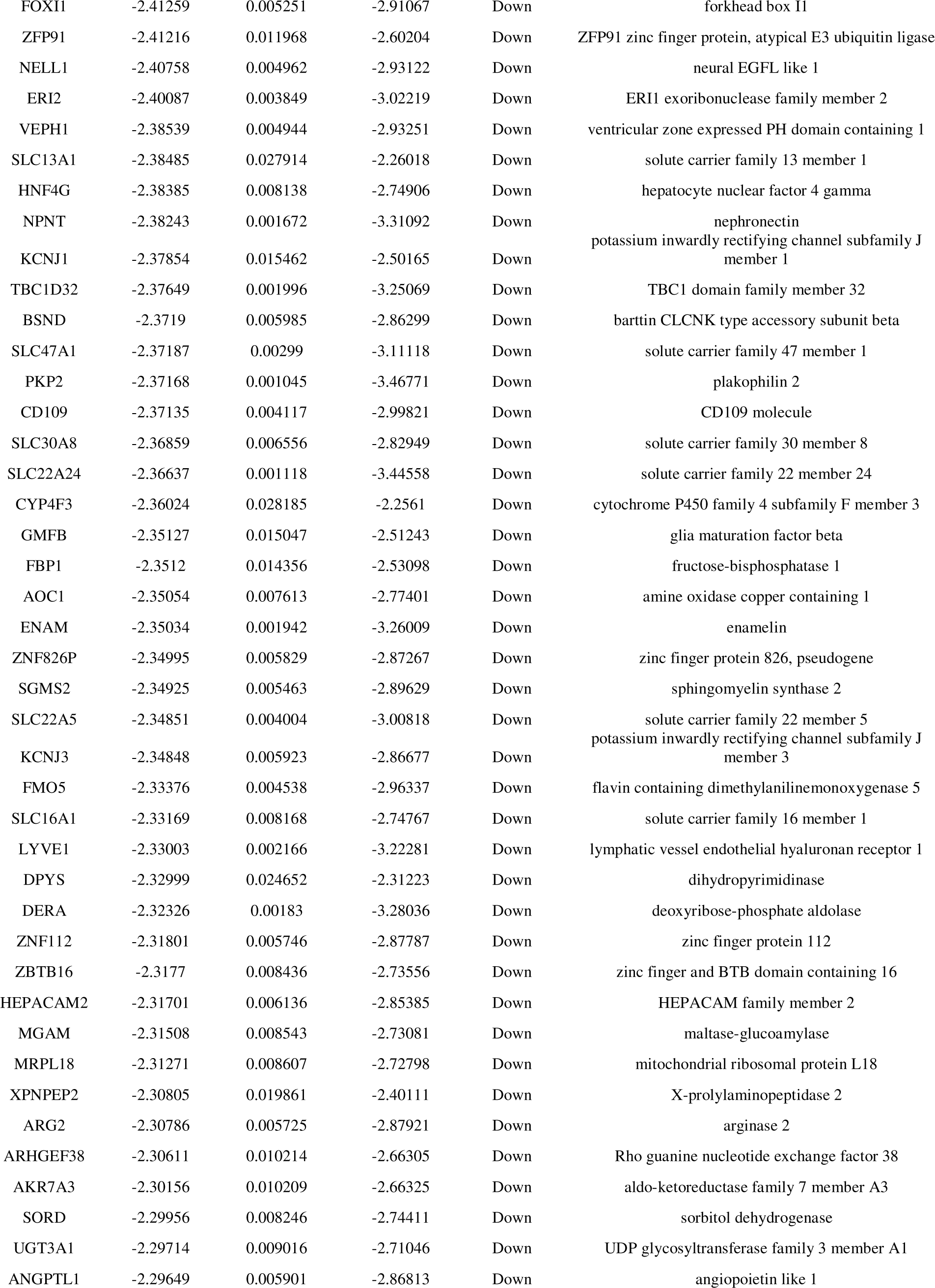
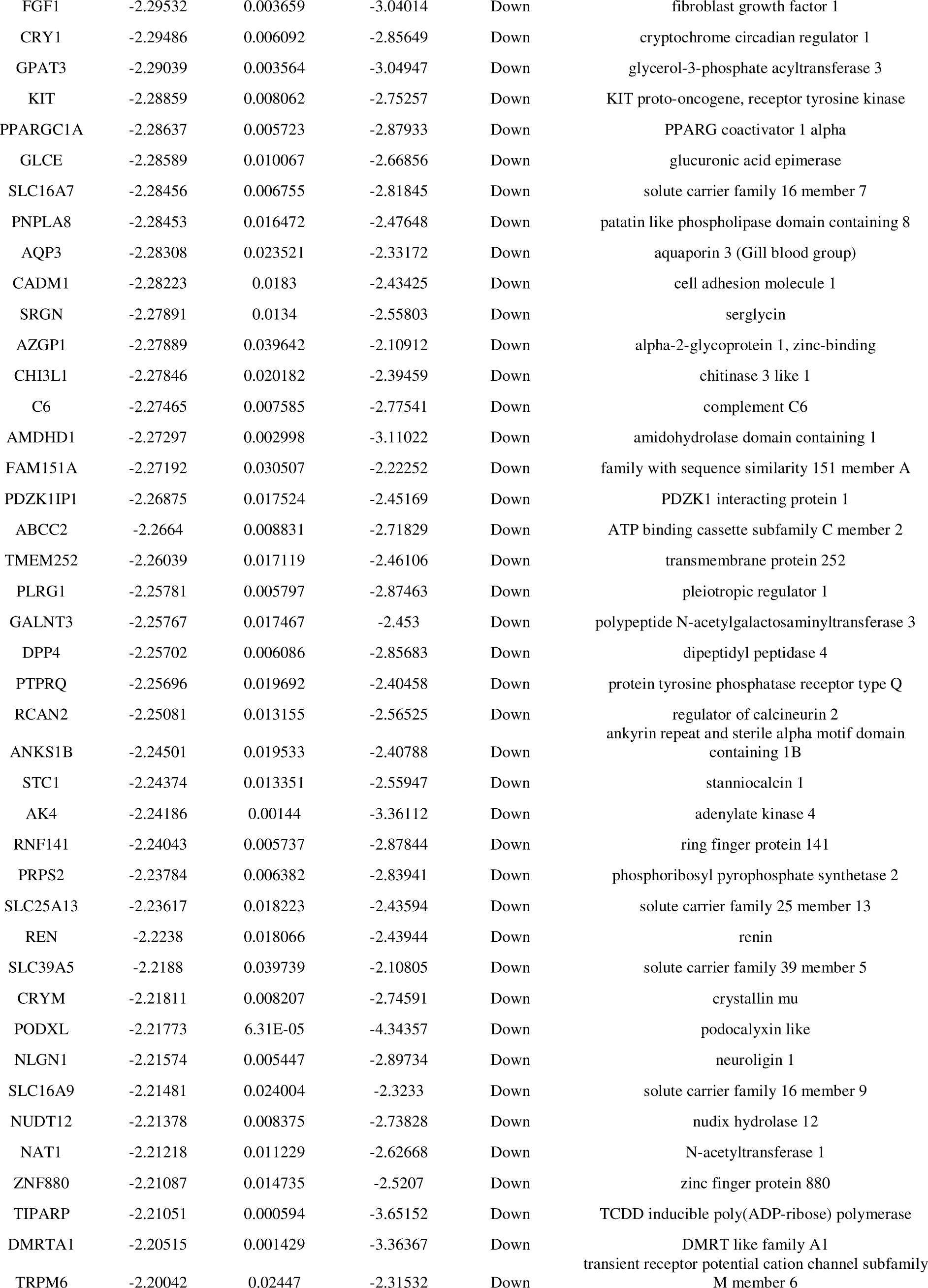
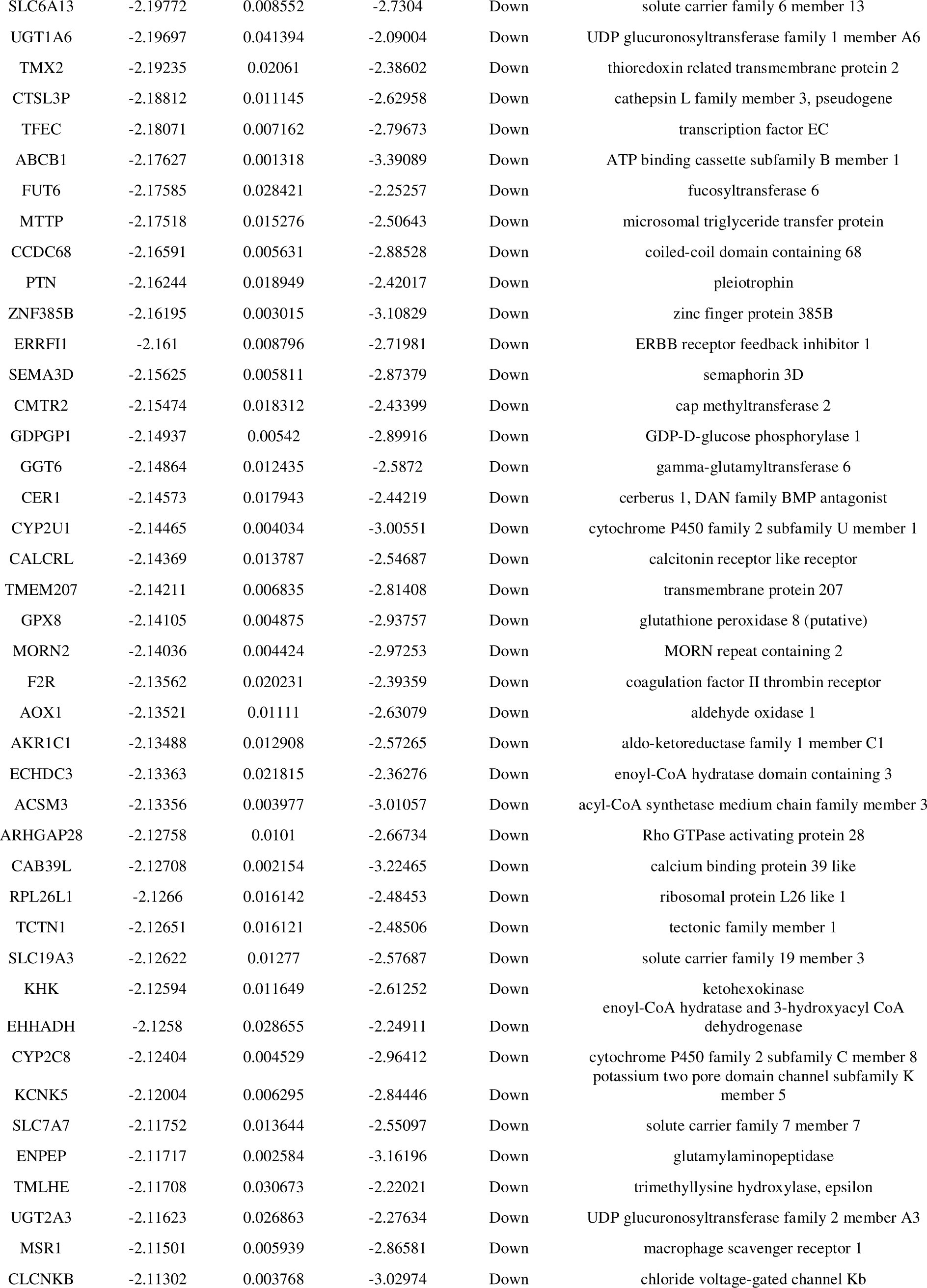
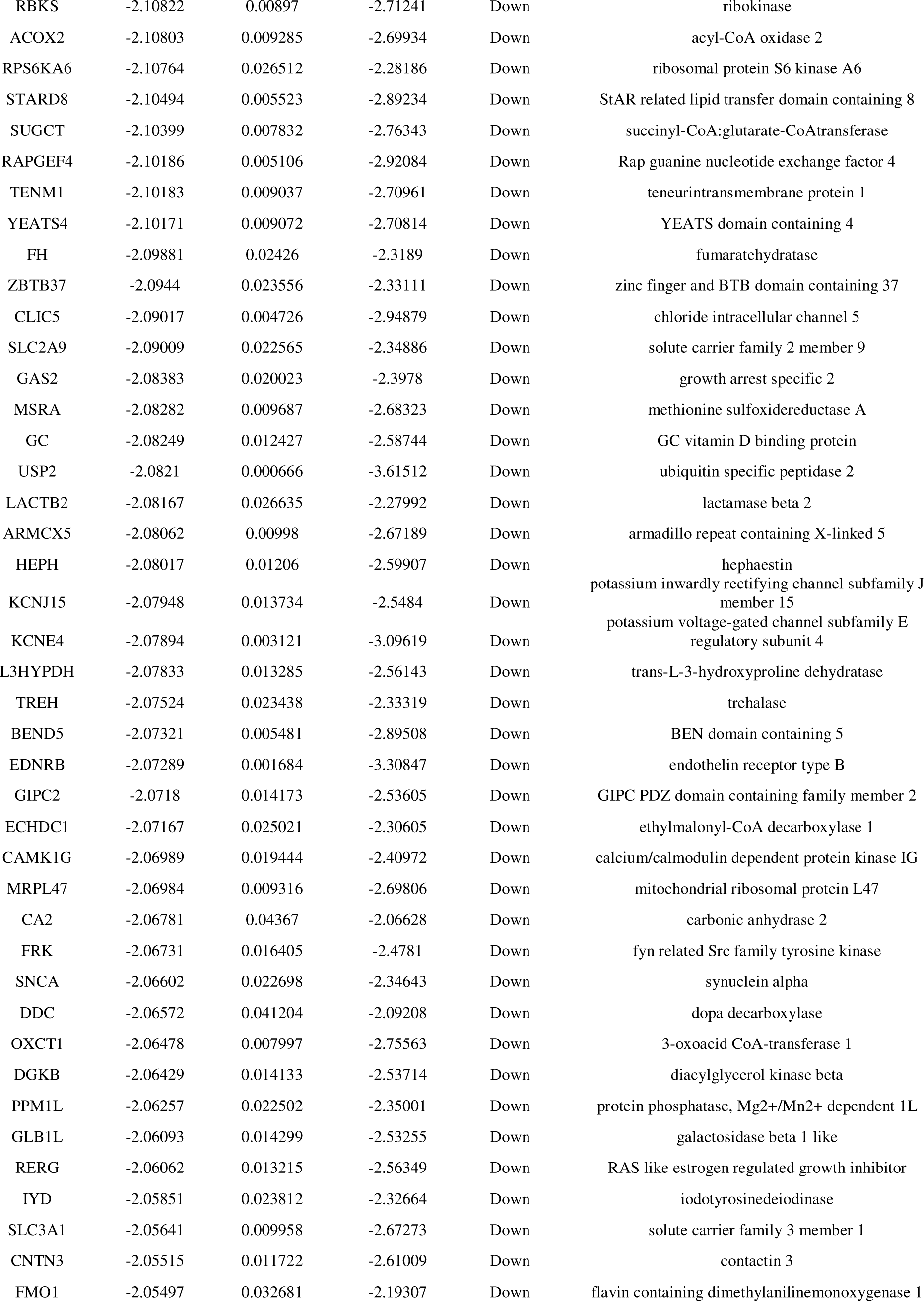
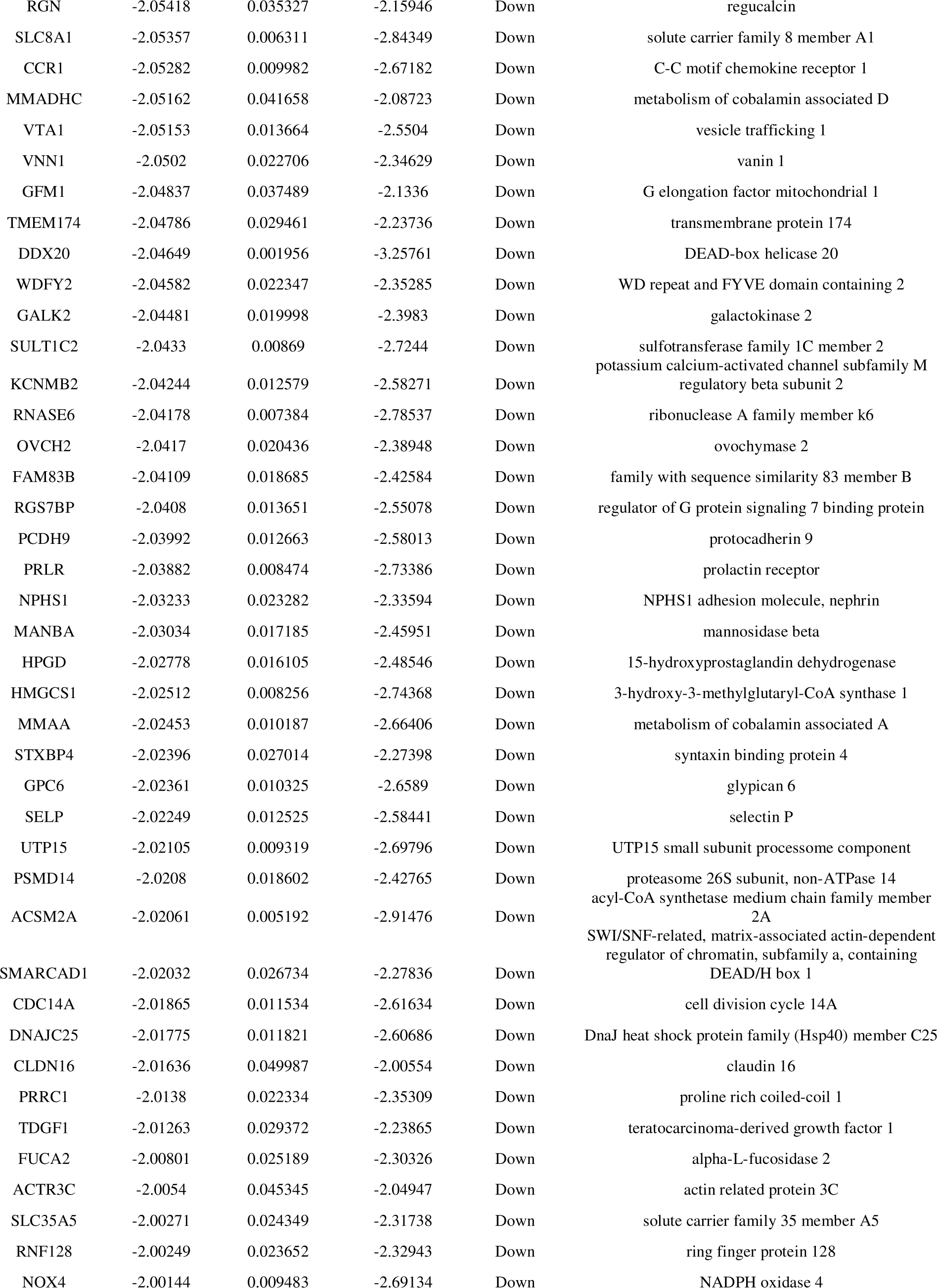
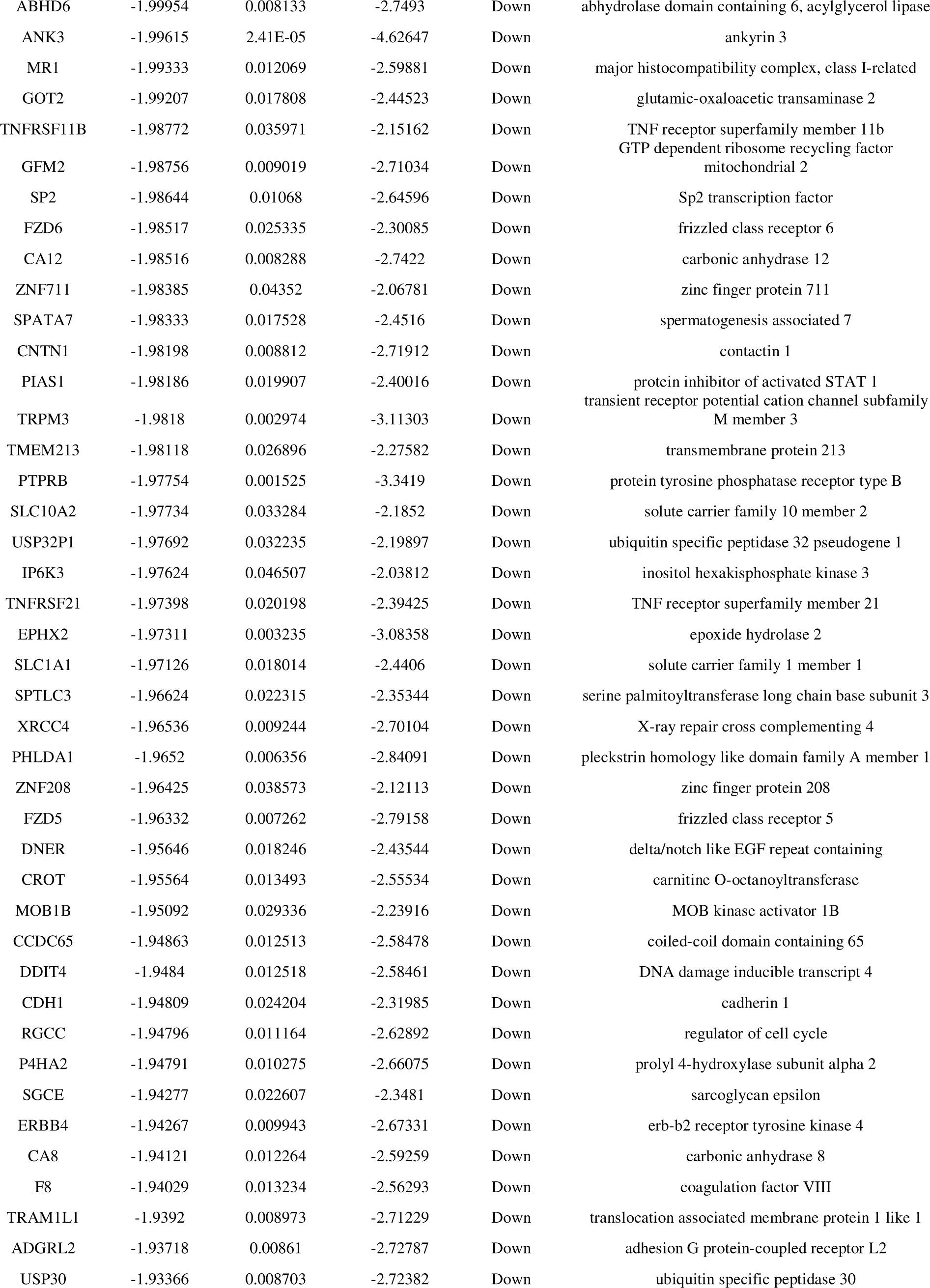
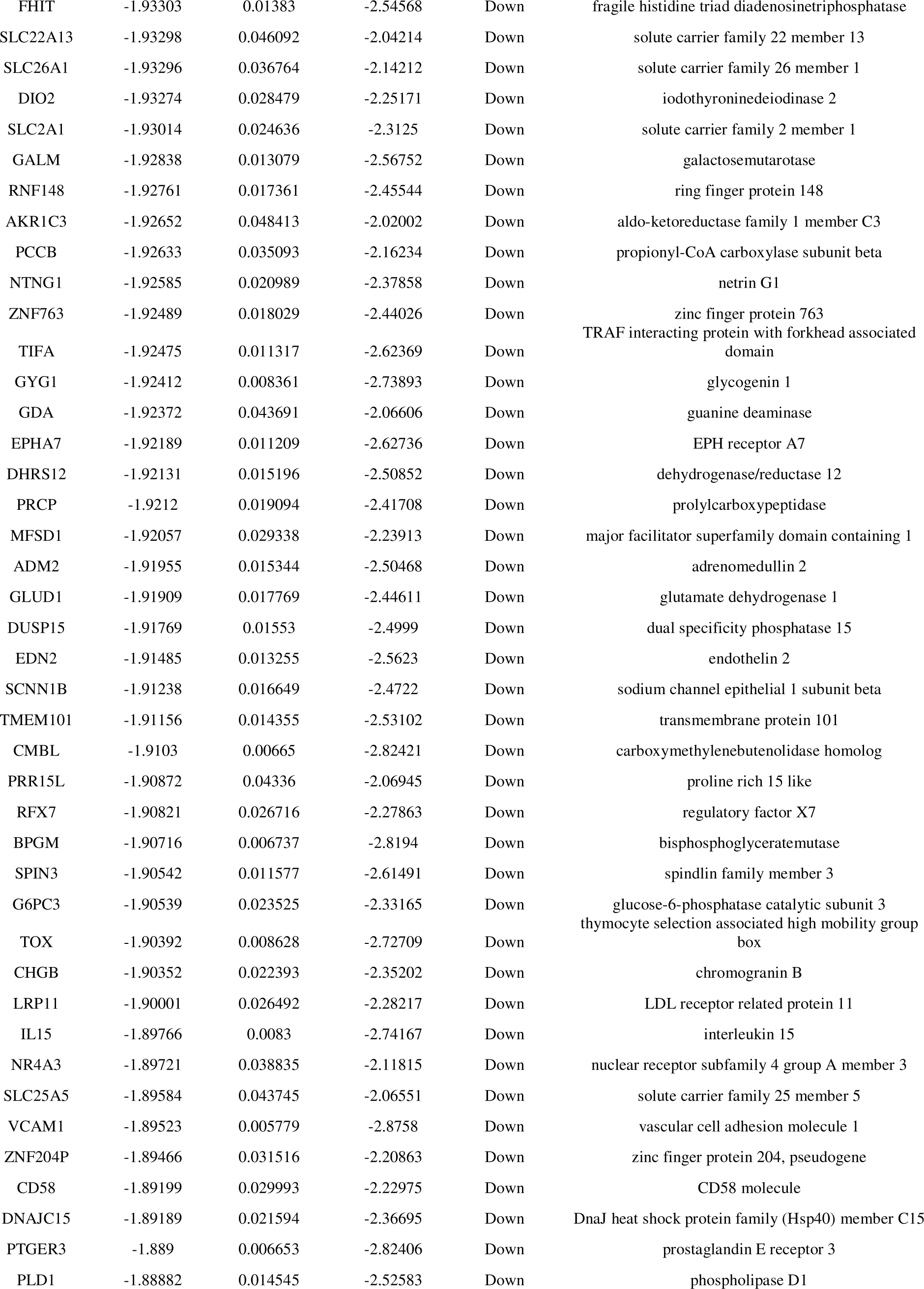
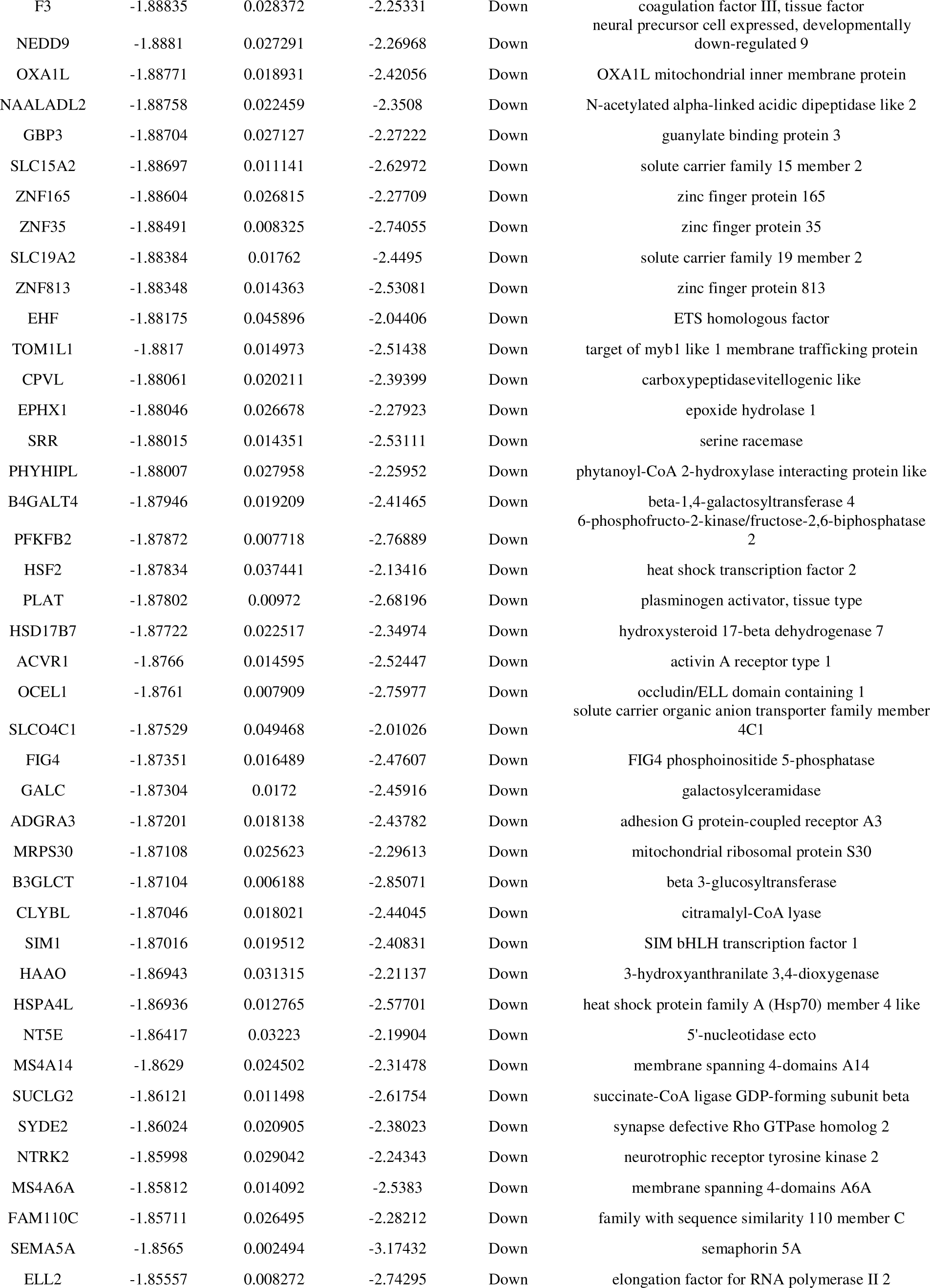
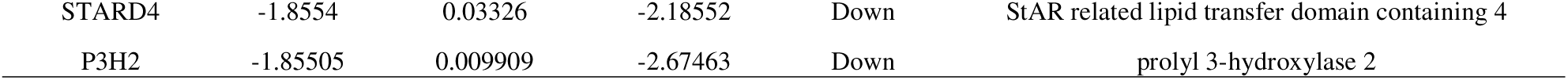
The statistical metrics for key differentially expressed genes (DEGs)

### GO and pathway enrichment analyses of DEGs

The GO and REACTOME pathway enrichment analyses were used to investigate the biological activities and pathways involved in the 956 DEGs. GO enrichment analysis was used to further explore the role of DEGs in AKI. BP is mainly concentrated in metabolic process, biosynthetic process, small molecule metabolic process and response to chemical (Table 2). In the CC, it is mainly enriched in the intracellular non-membrane-bounded organelle, nucleus, membrane and cytoplasm (Table 2). MF is mainly concentrated in the activities of such as sequence-specific DNA binding, DNA-binding transcription activator activity, catalytic activity and transporter activity (Table 2). As a result of REACTOME pathway enrichment analysis, the DEGs mainly involved in striated muscle contraction, muscle contraction, metabolism and drug ADME (Table 3).

**Table 2.**
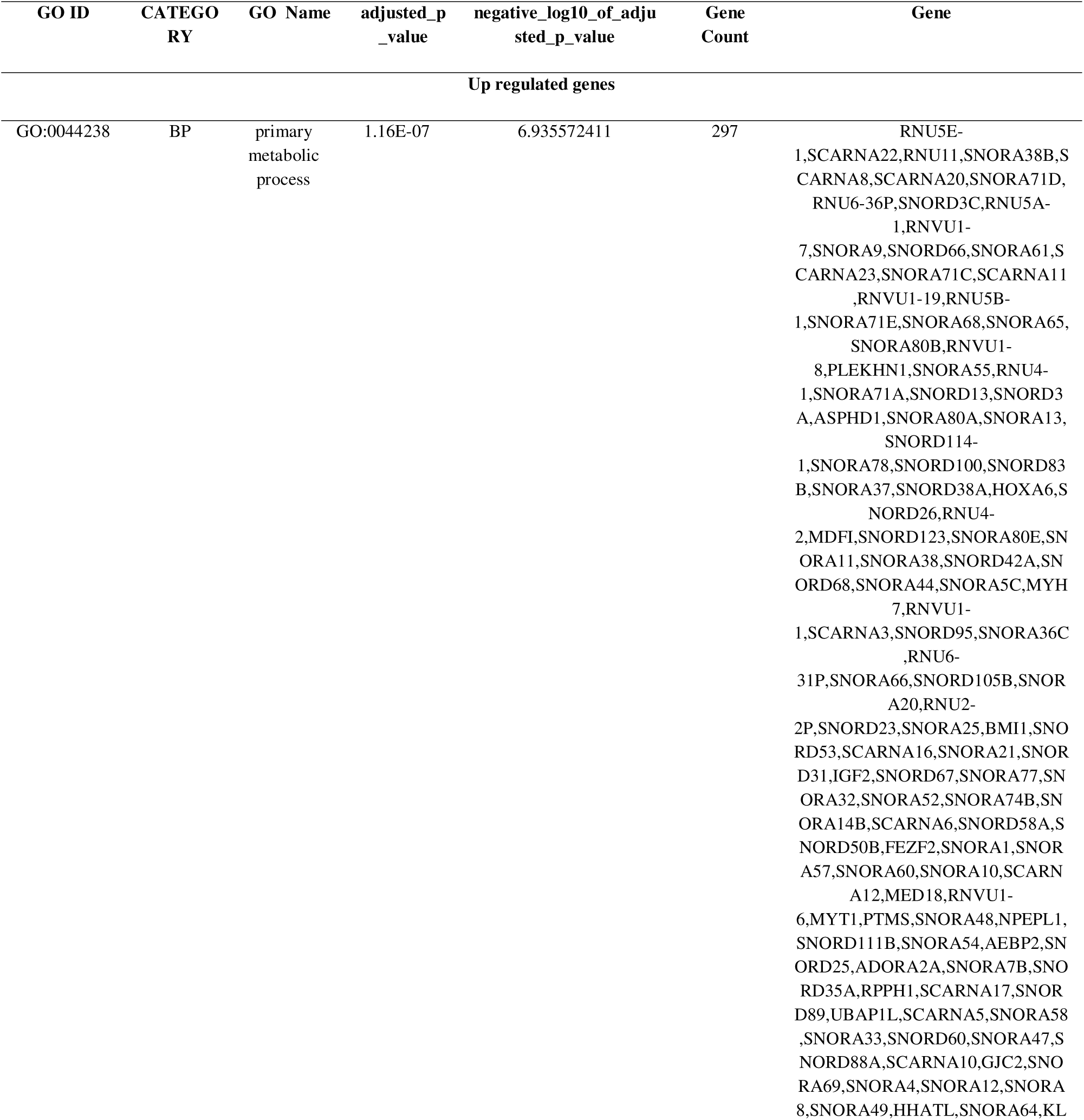
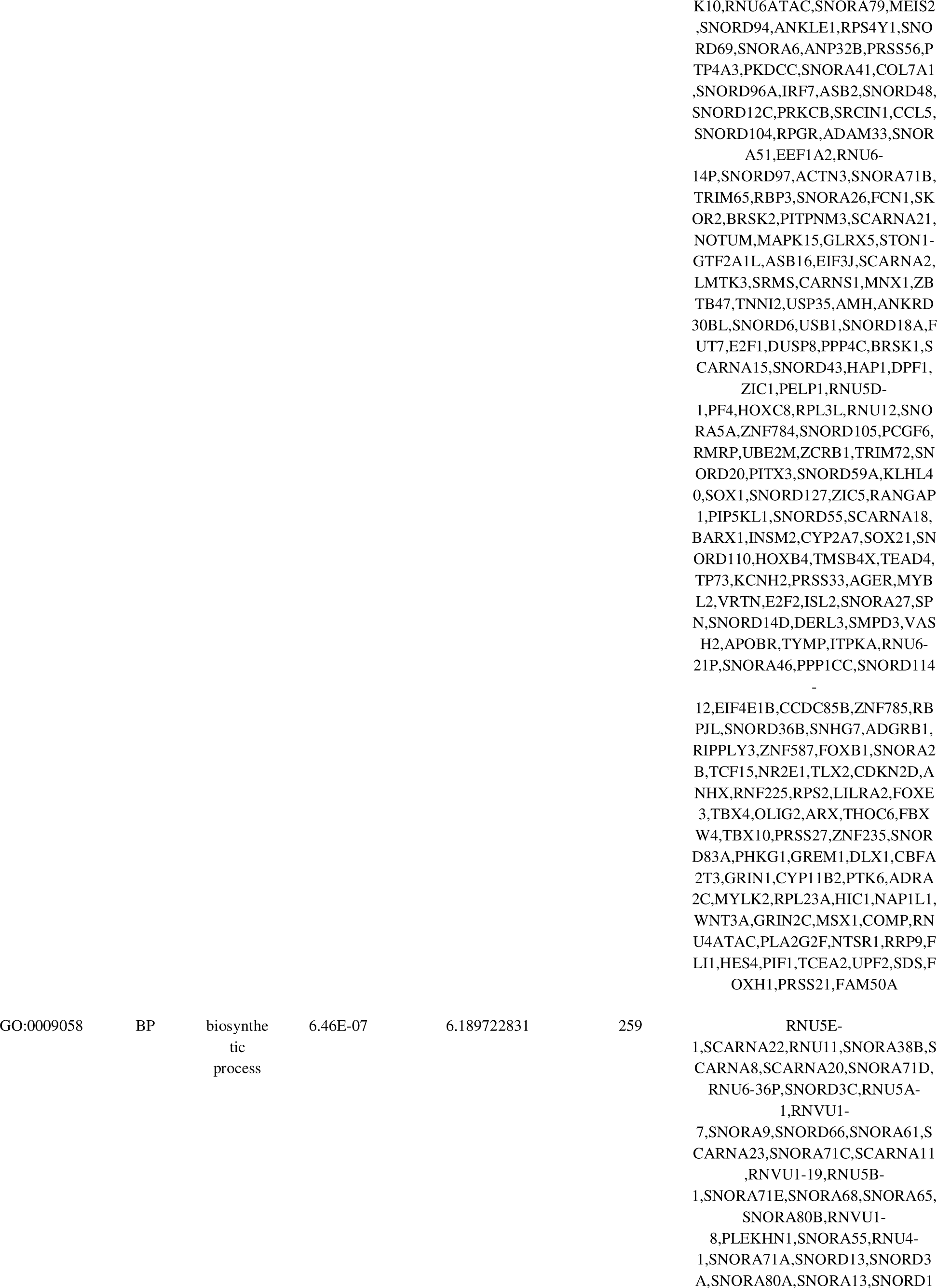
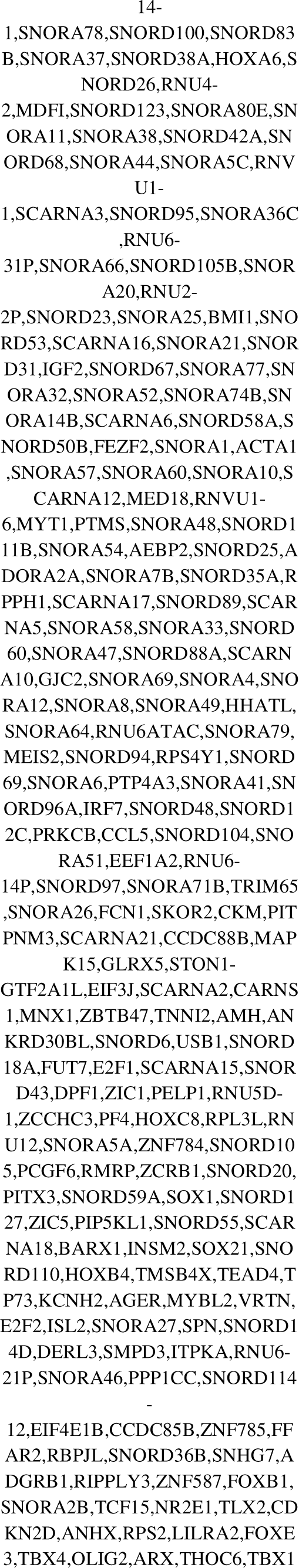
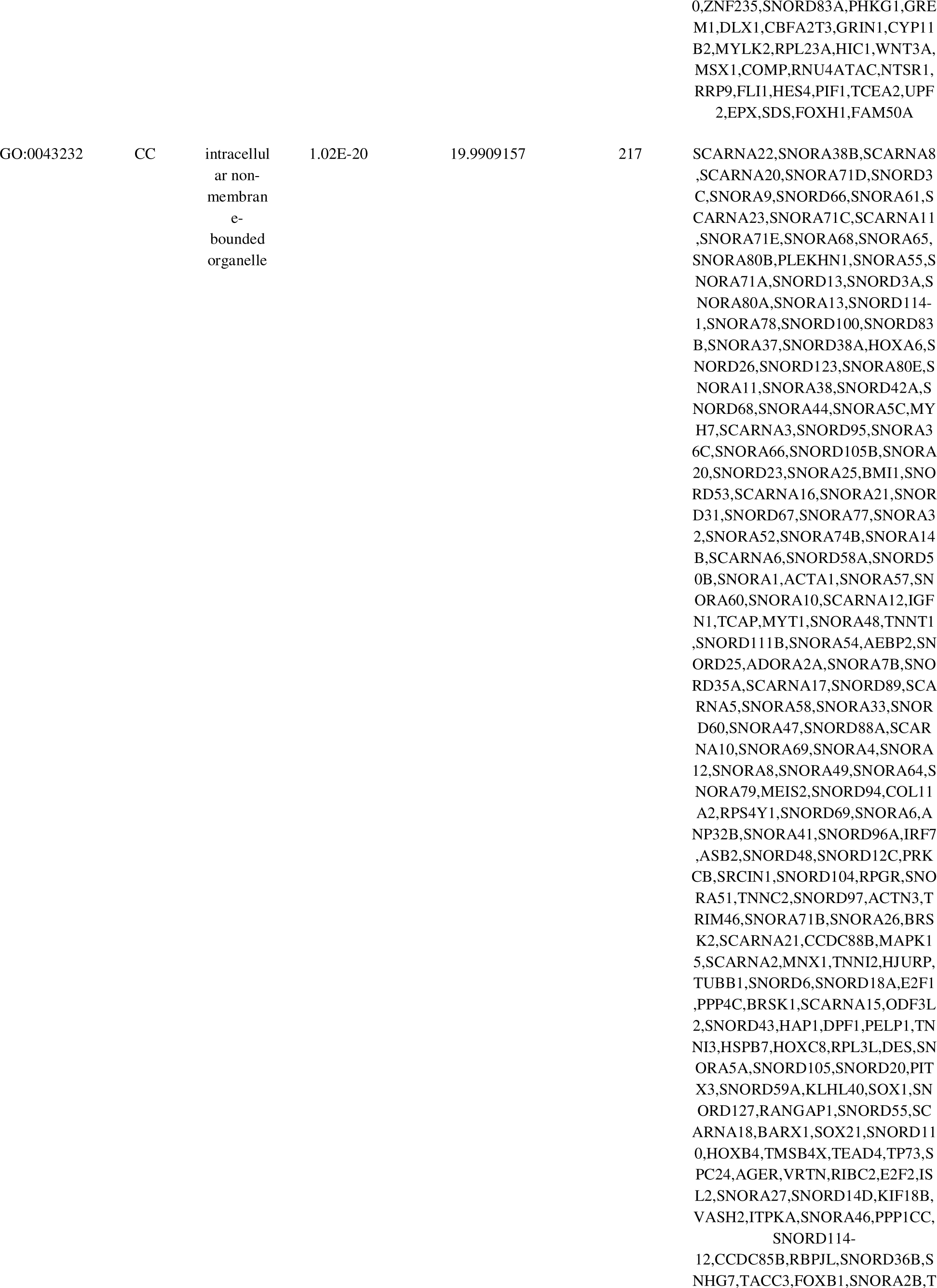
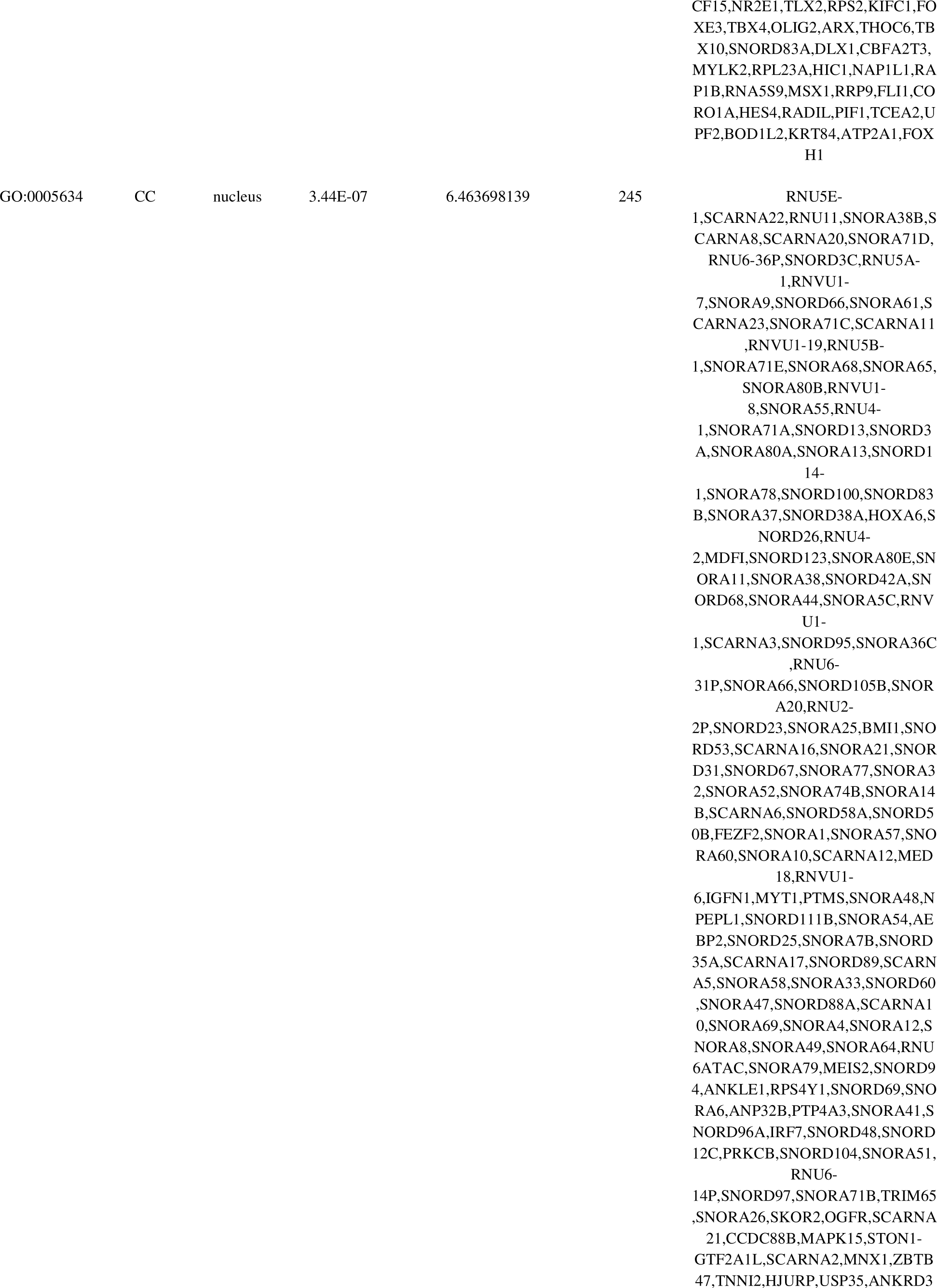
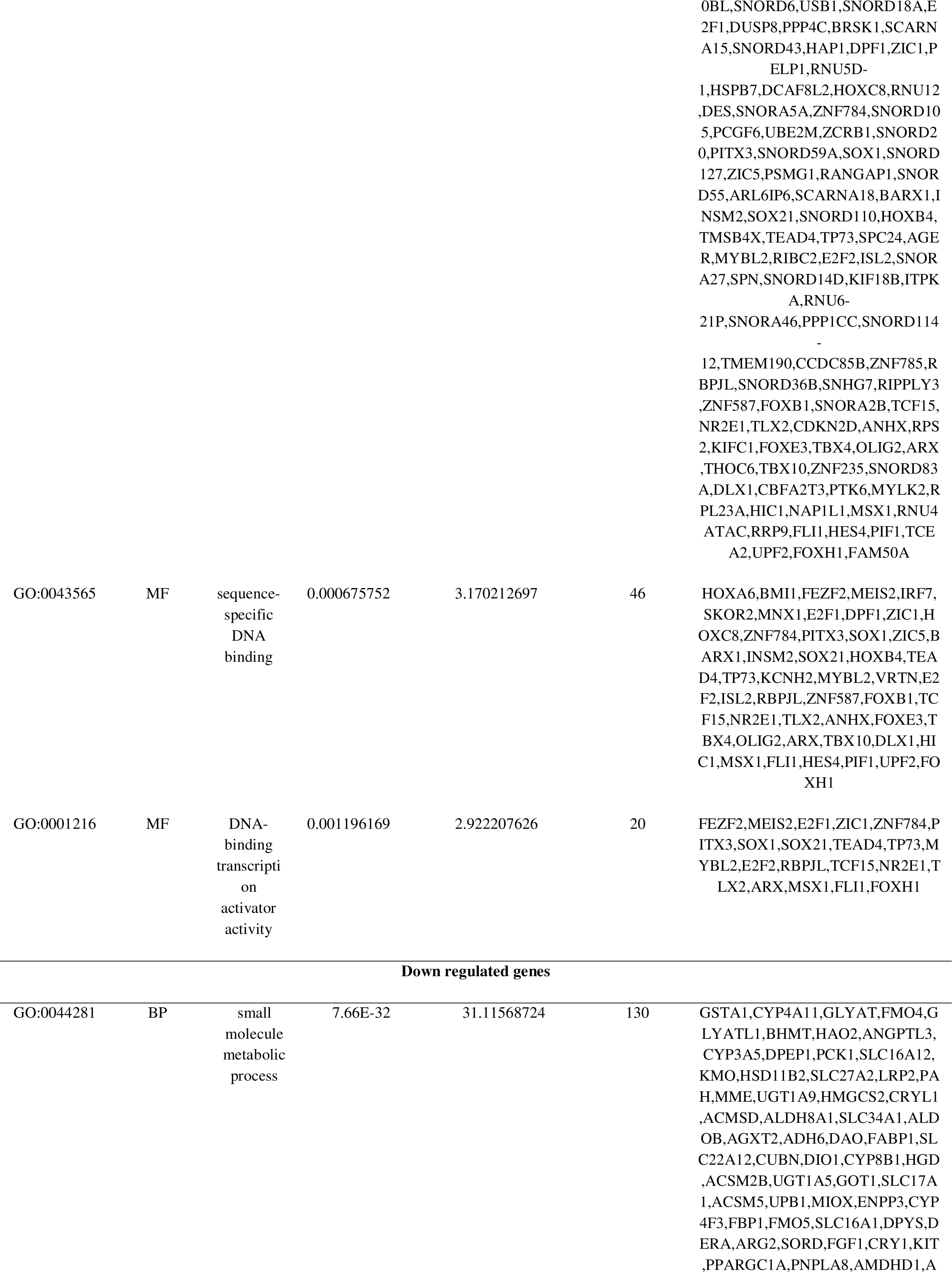
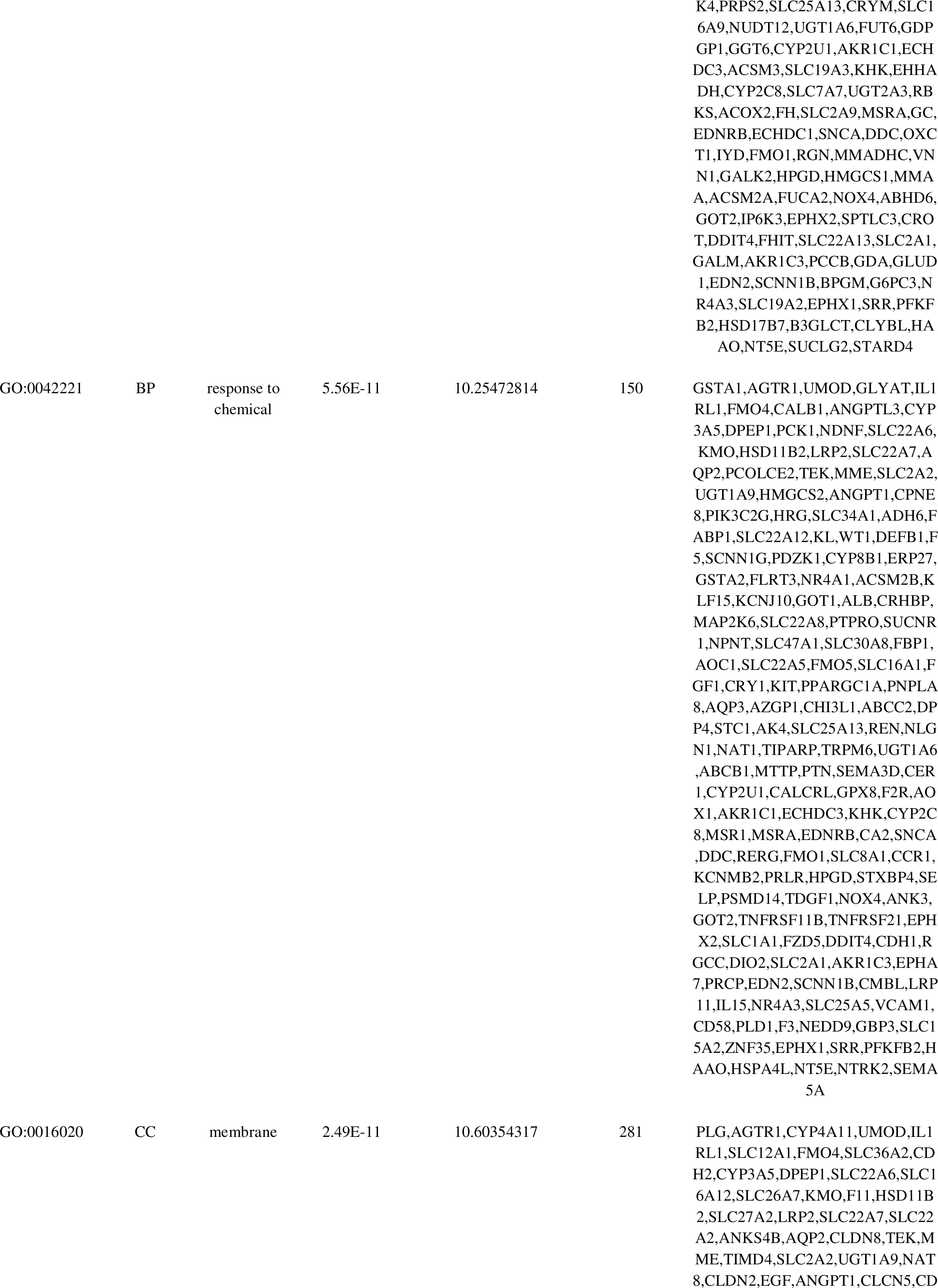
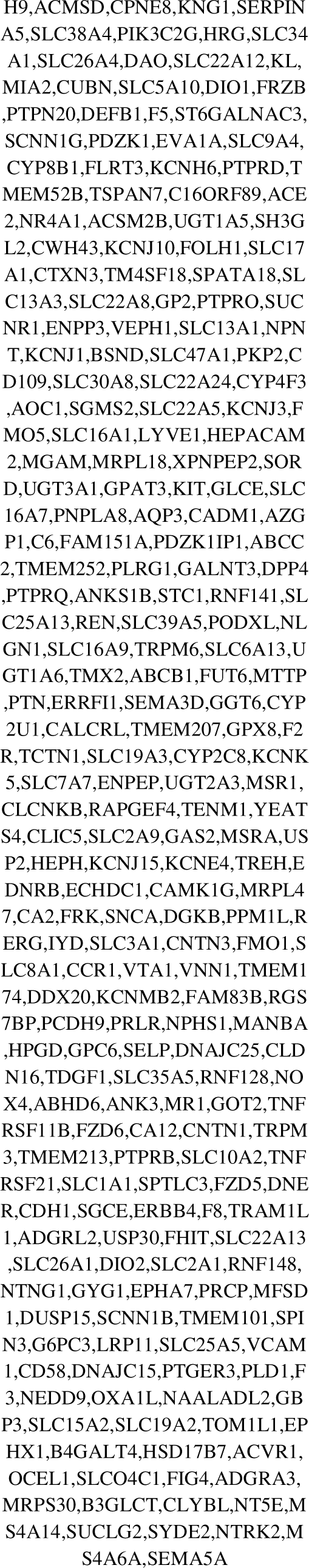
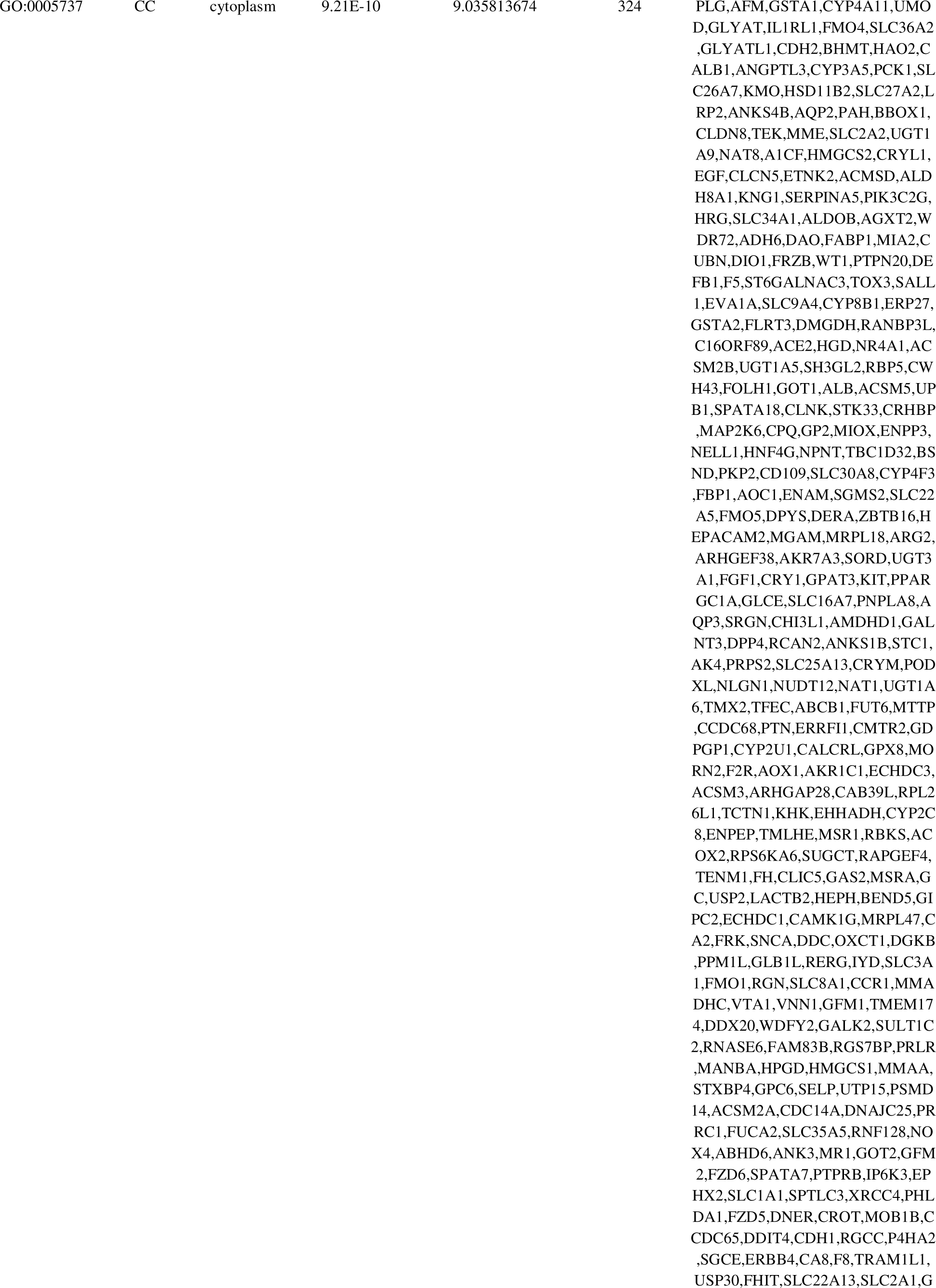
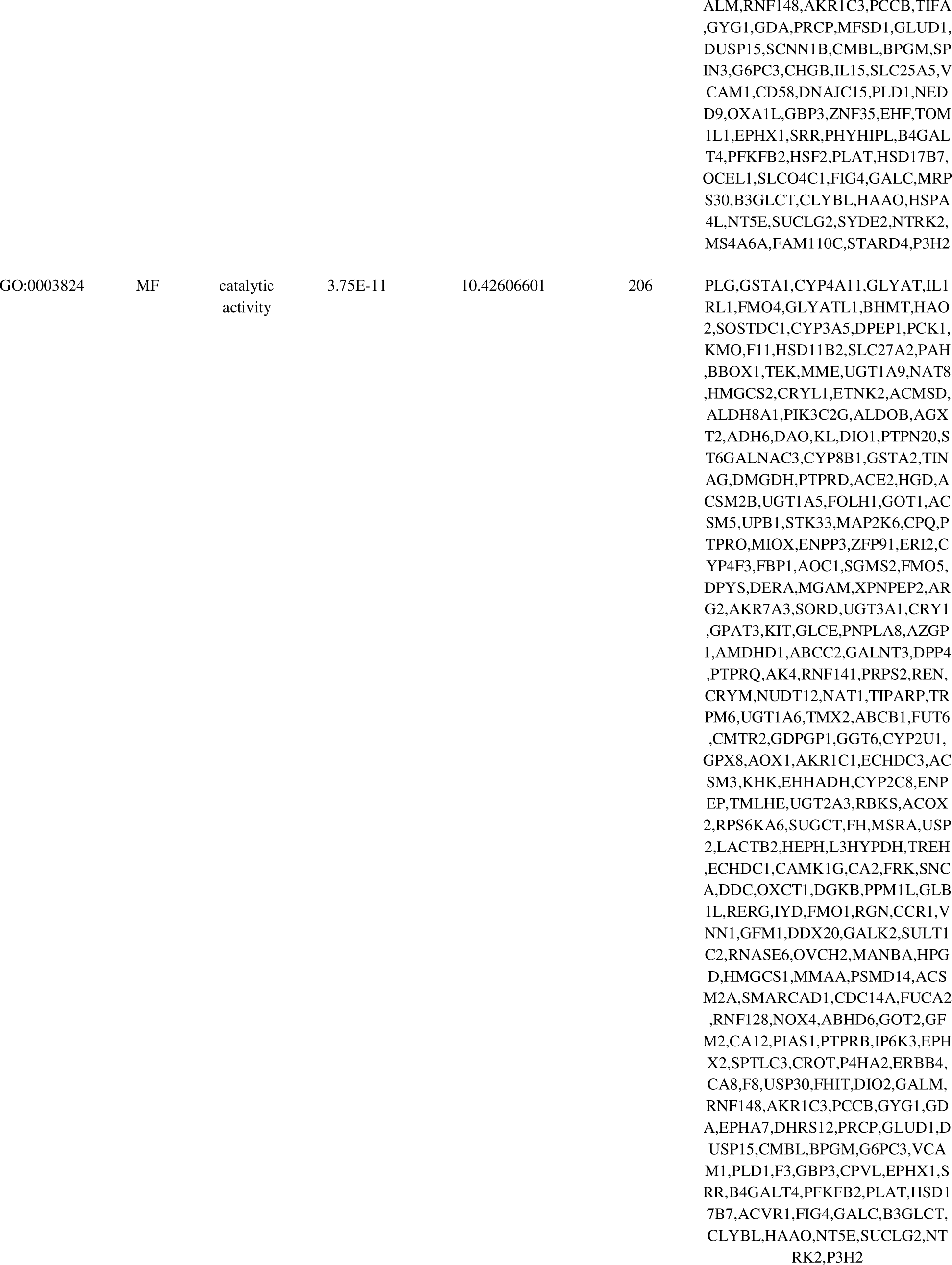
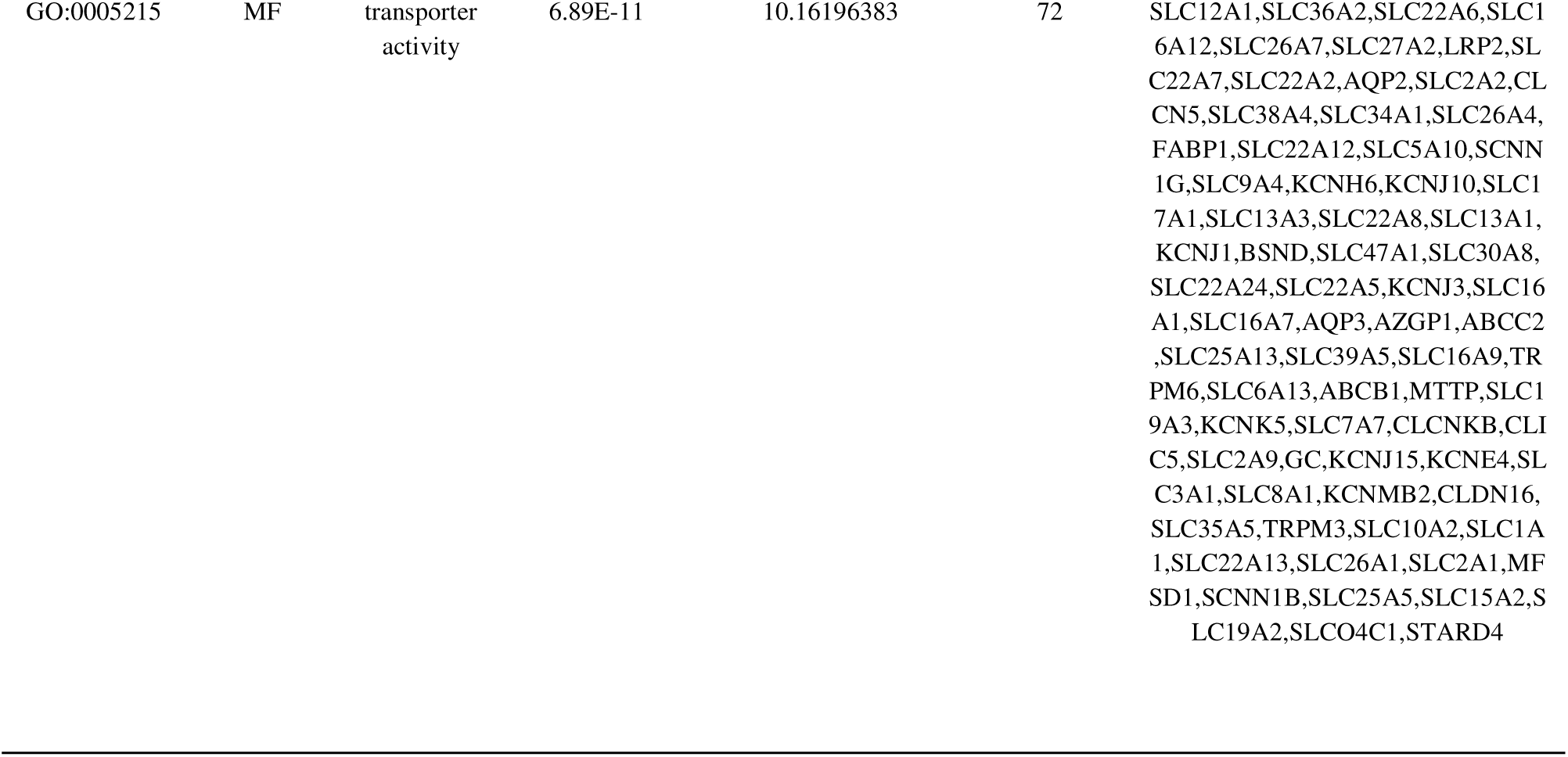
The enriched GO terms of the up and down regulated differentially expressed genes.

**Table 3.**
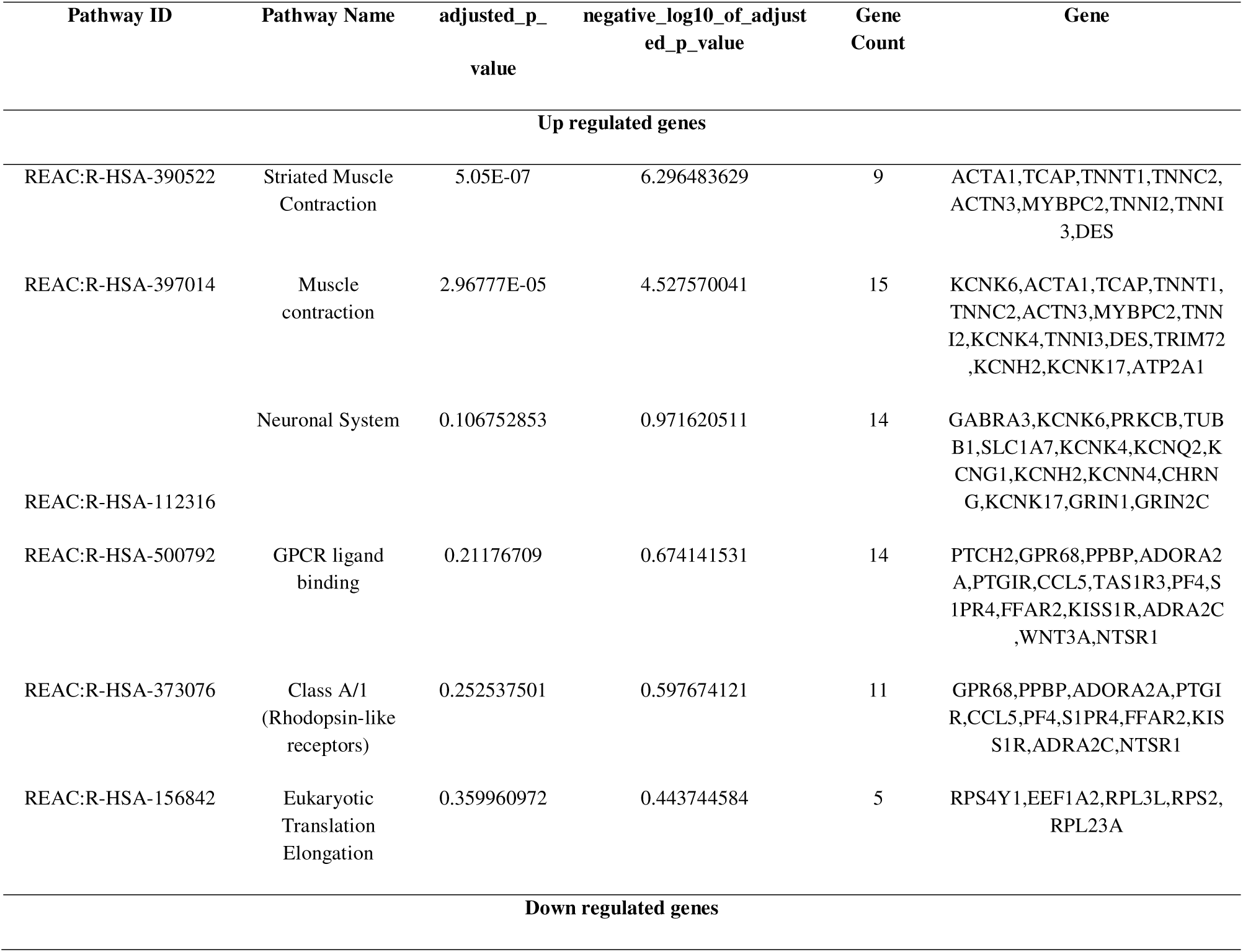
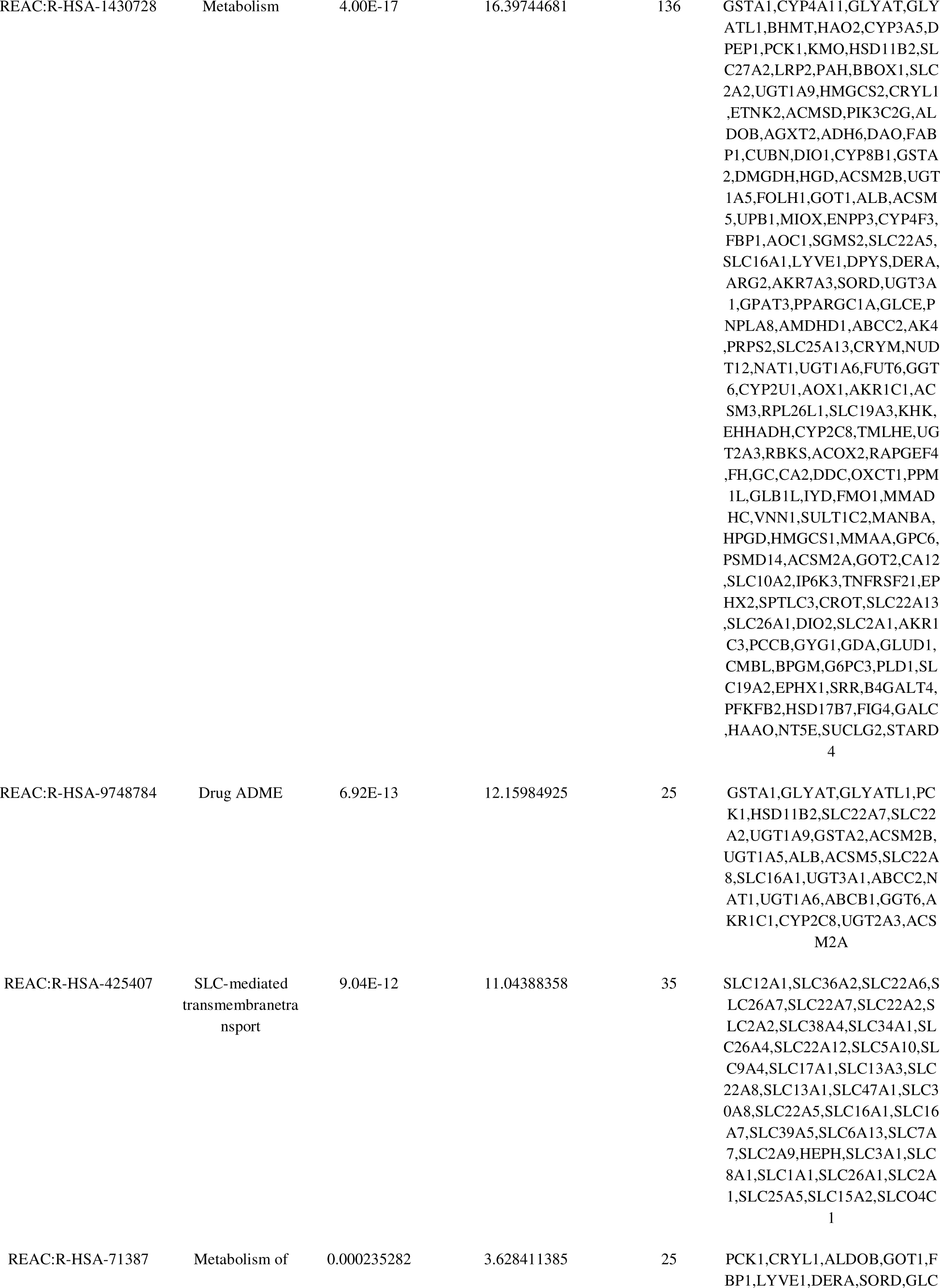
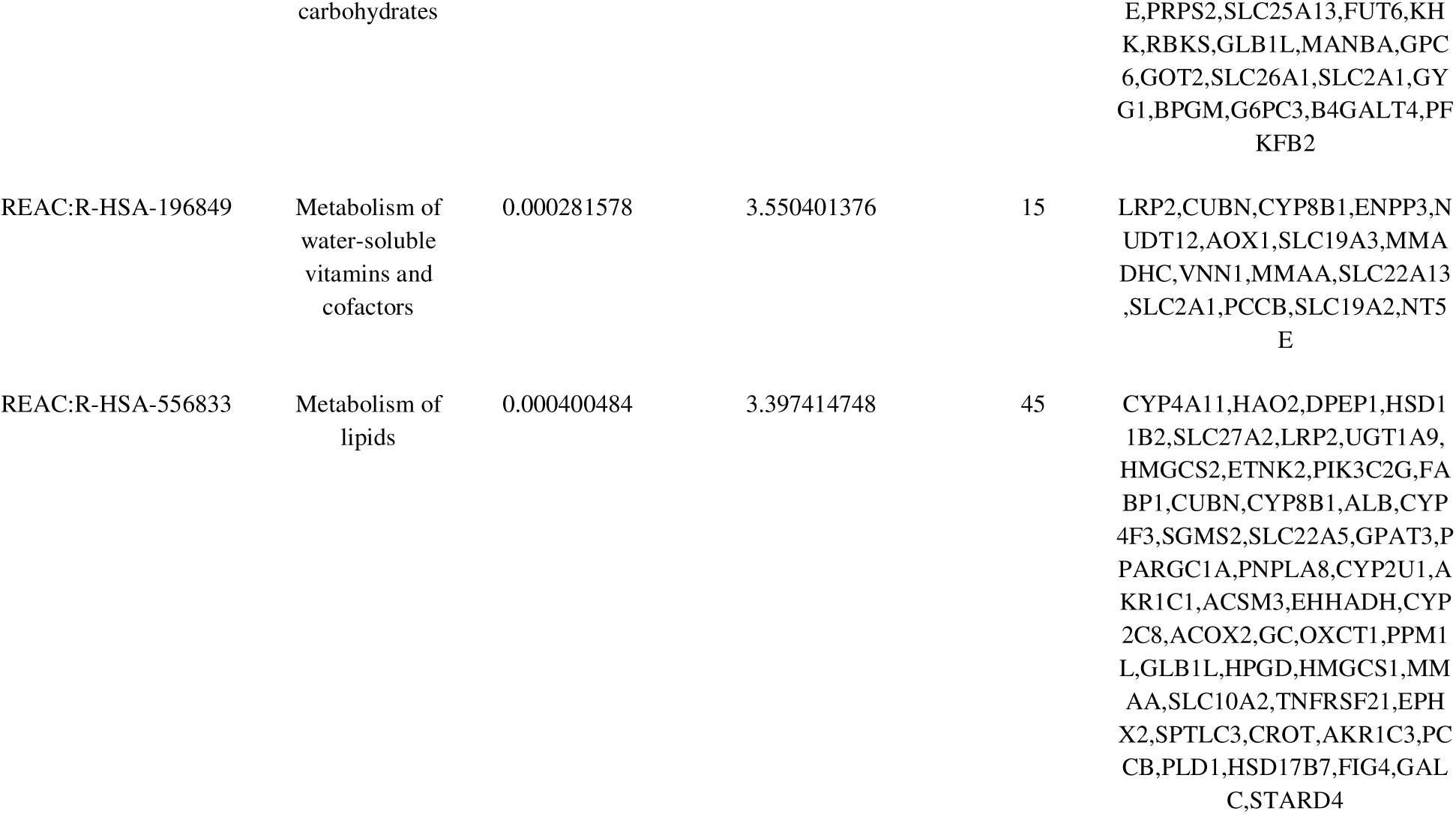
The enriched pathway terms of the up and down regulated differentially expressed genes.

### Construction of the PPI network and module analysis

Based on the HIPIE online database, a total of 956 DEGs were imported into the DEG PPI network complex which included 5038 nodes and 8544 edges (Fig.3). The Network Analyzer plugin of Cytoscape was used to score each node gene by 4 selected algorithms, including degree, betweenness, stress and closeness. Finally, we identified ten hub genes include PPP1CC, RPS2, MDFI, BMI1, RPL23A, VCAM1, ALB, SNCA, DPP4 and RPL26L1 (Table 4). In order to figure out the core modules of complex network, we performed PEWCC plug-in analysis and identified 2 modules. The most significant module 1 was shown in Fig. 4A, containing 19 nodes and 37 edges, and most significant module 2 was shown in Fig. 4B, containing 11 nodes and 24 edges. GO and pathway enrichment analysis showed that these genes are related to the primary metabolic process, biosynthetic process, metabolism and membrane.

**Fig. 3.**
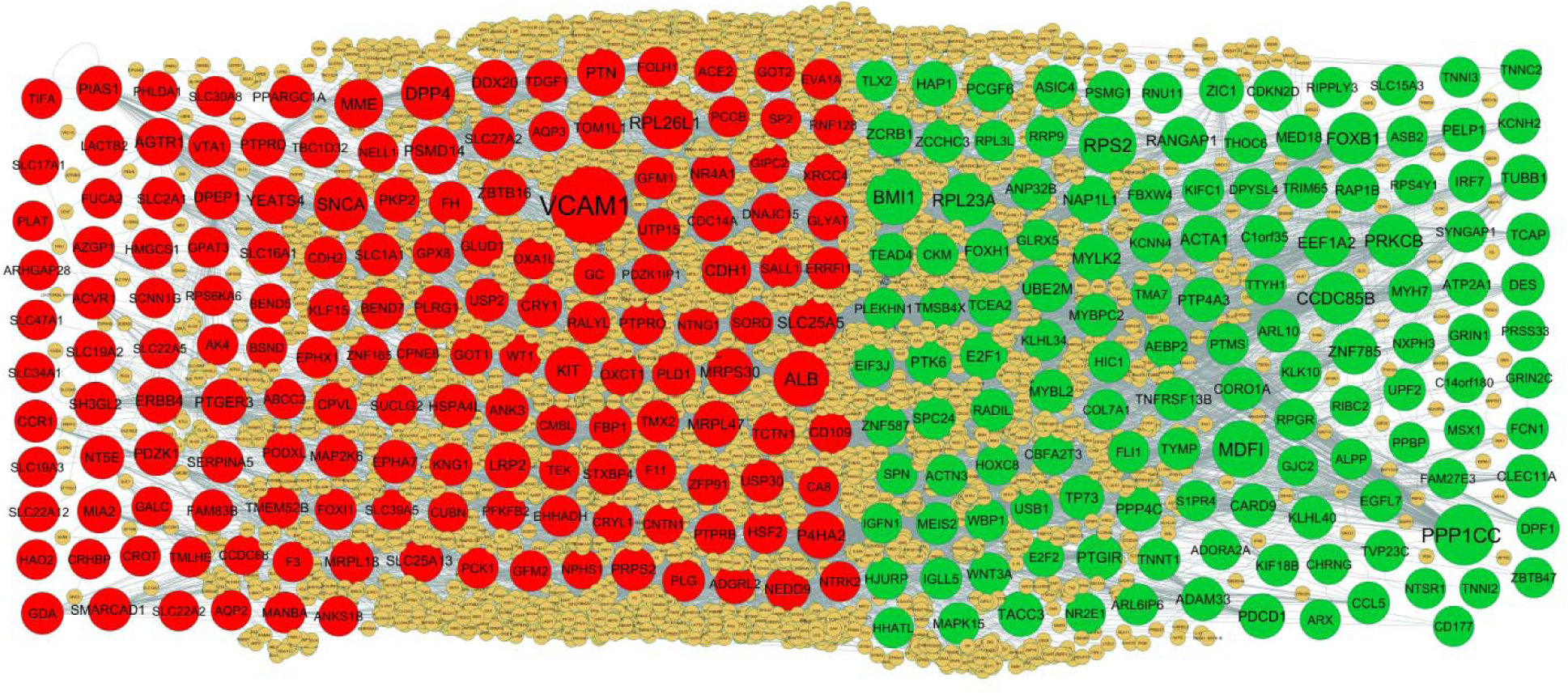
PPI network of DEGs. Up regulated genes are marked in parrot green; down regulated genes are marked in red.

**Fig. 4.**
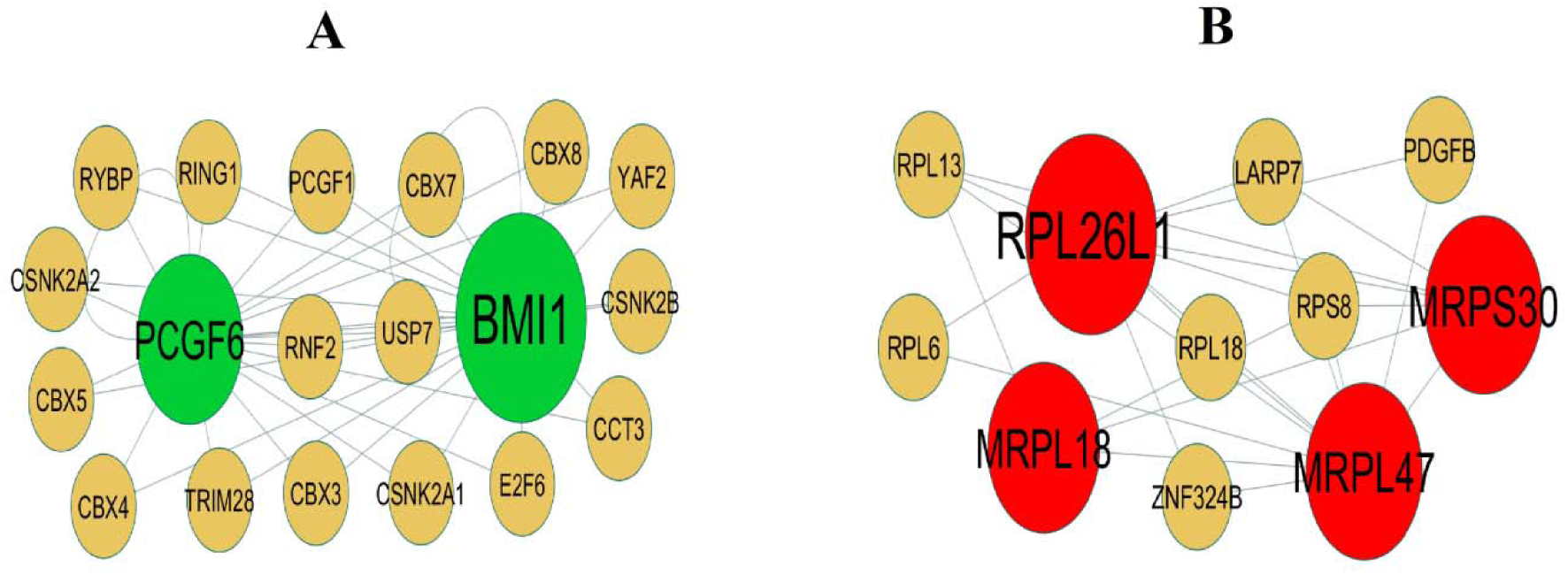
Modules selected from the PPI network. (A) The most significant module was obtained from PPI network with 19 nodes and 37 edges for up regulated genes (B) The most significant module was obtained from PPI network with 11 nodes and 24 edges for down regulated genes. Up regulated genes are marked in green; down regulated genes are marked in red.

**Table 4.**
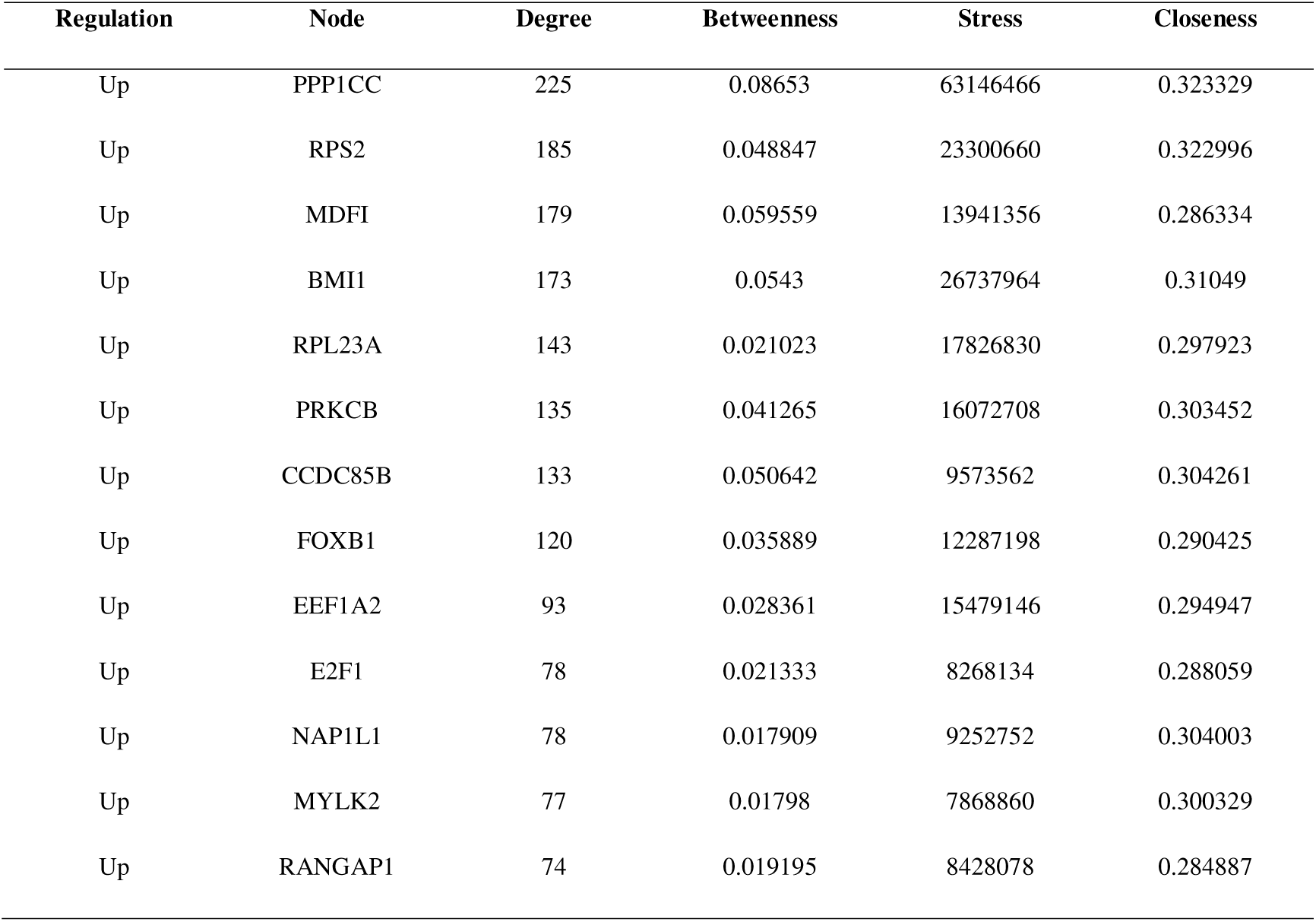
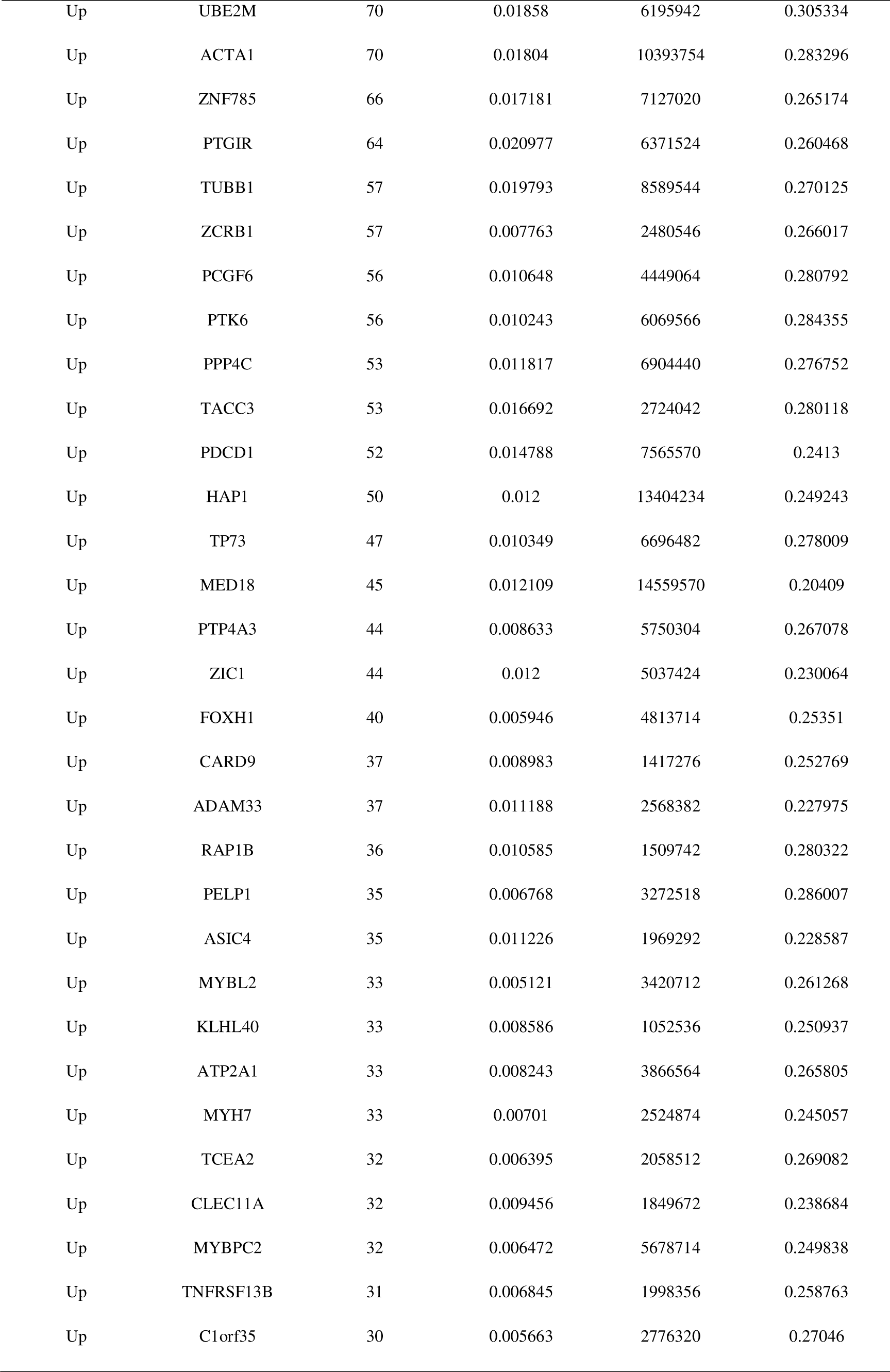
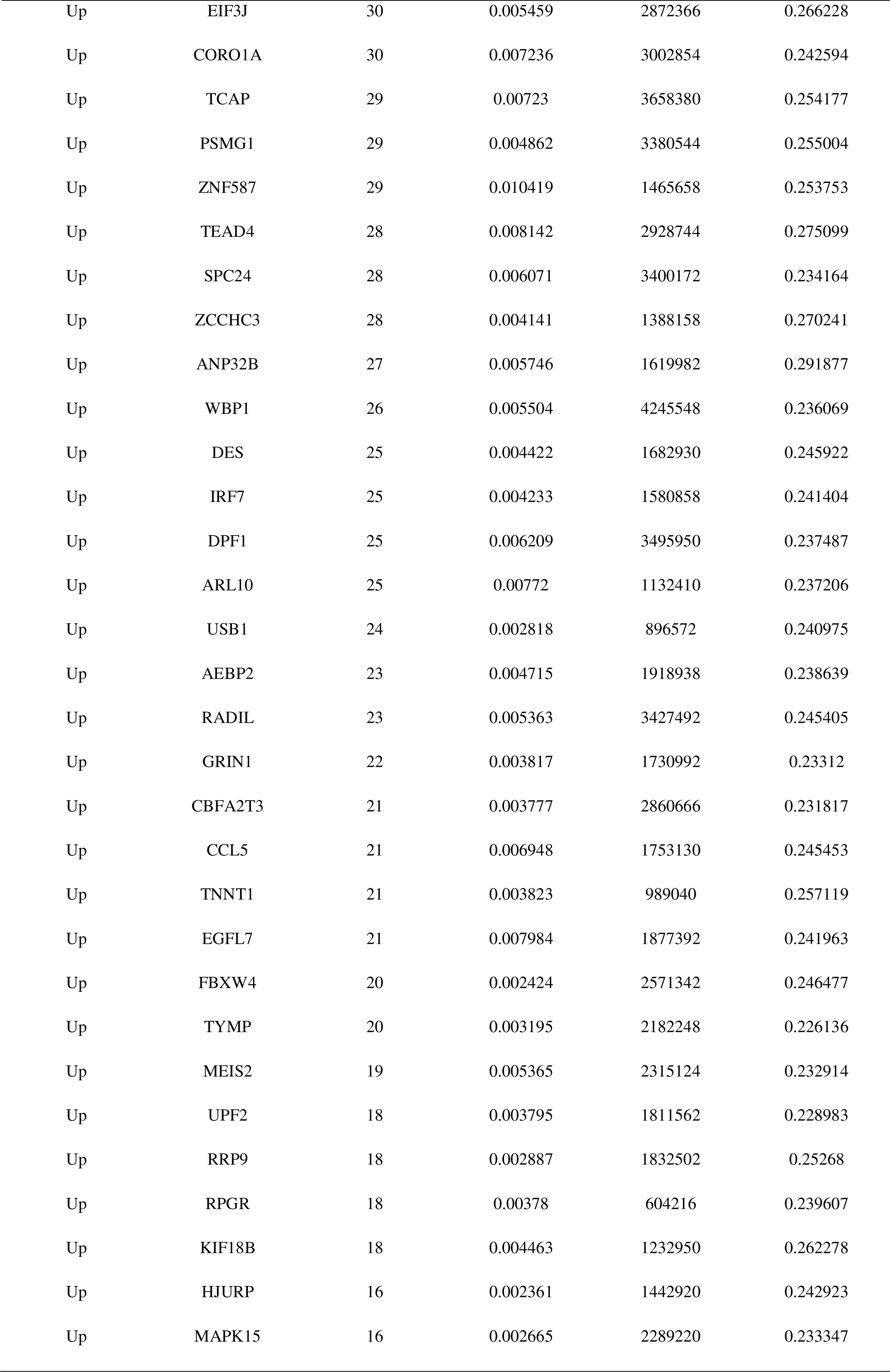
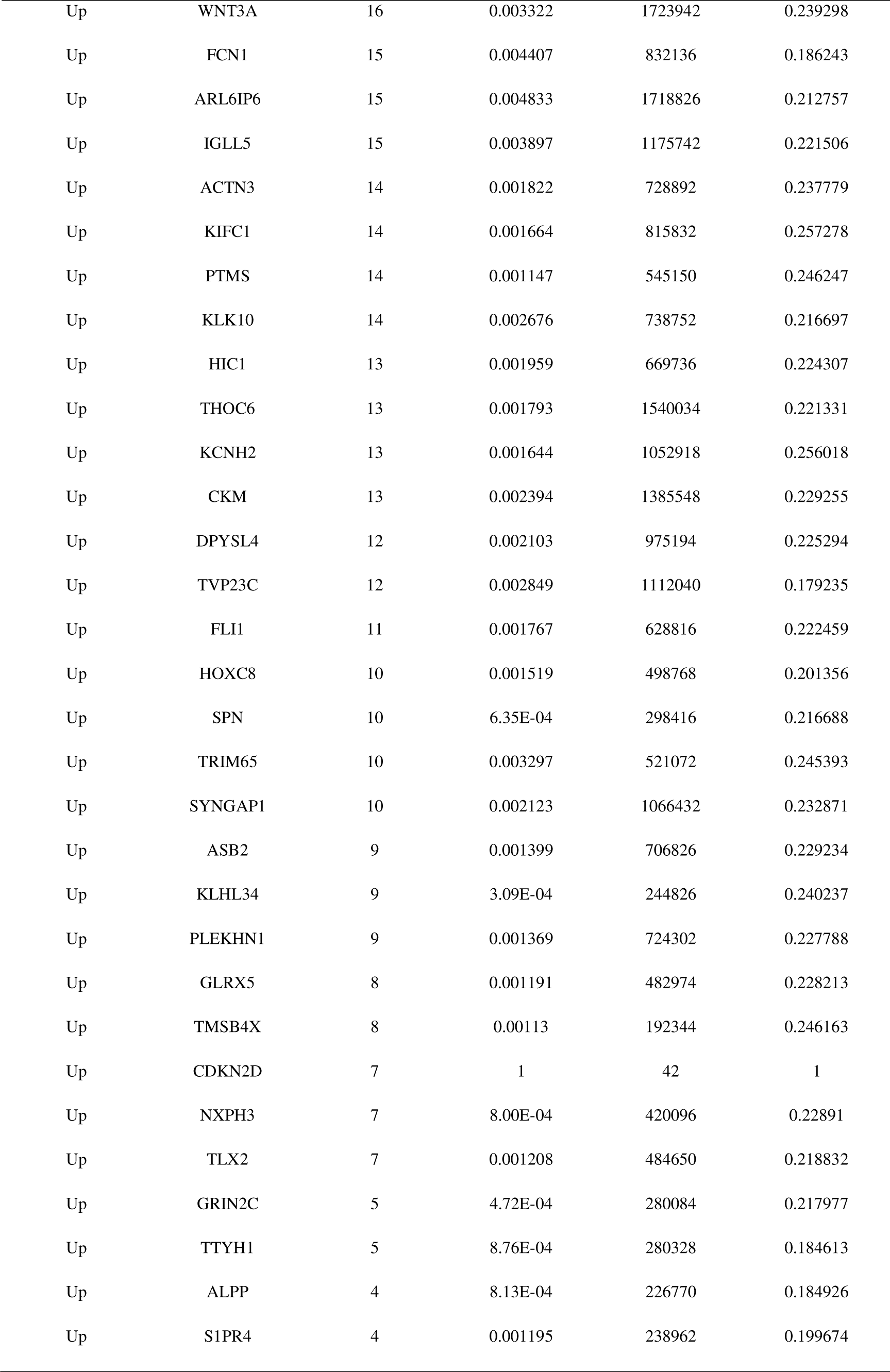
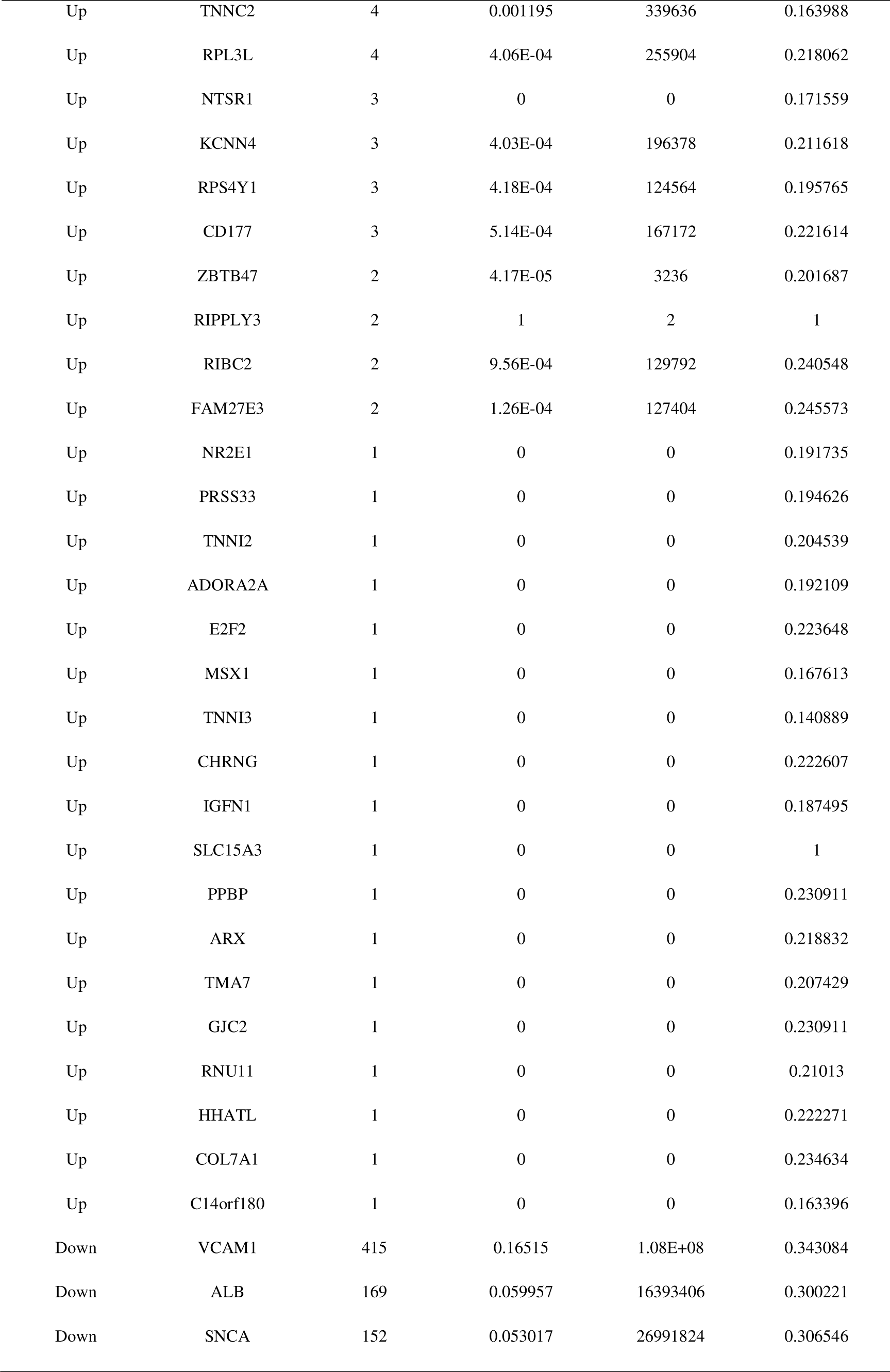
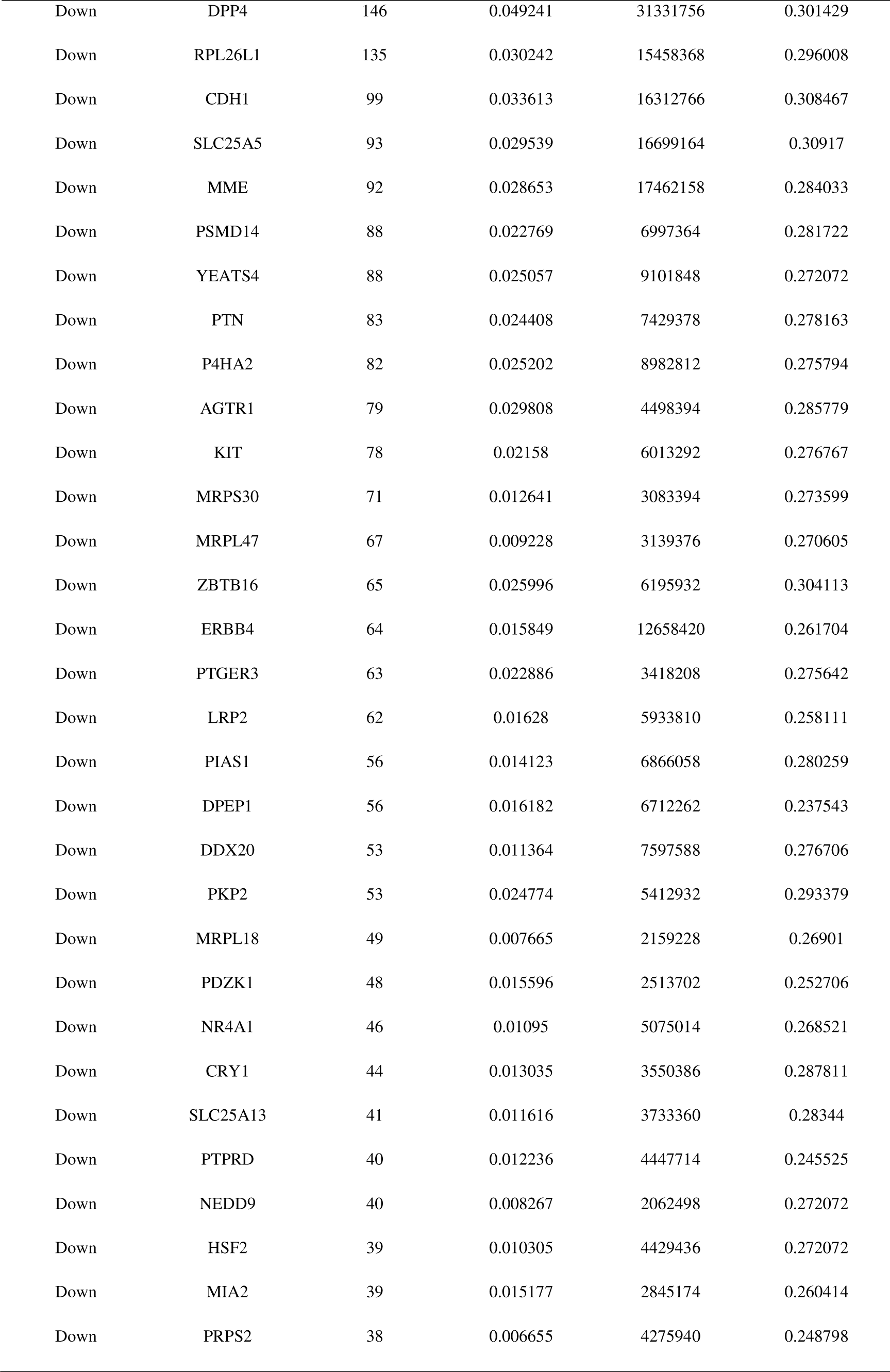
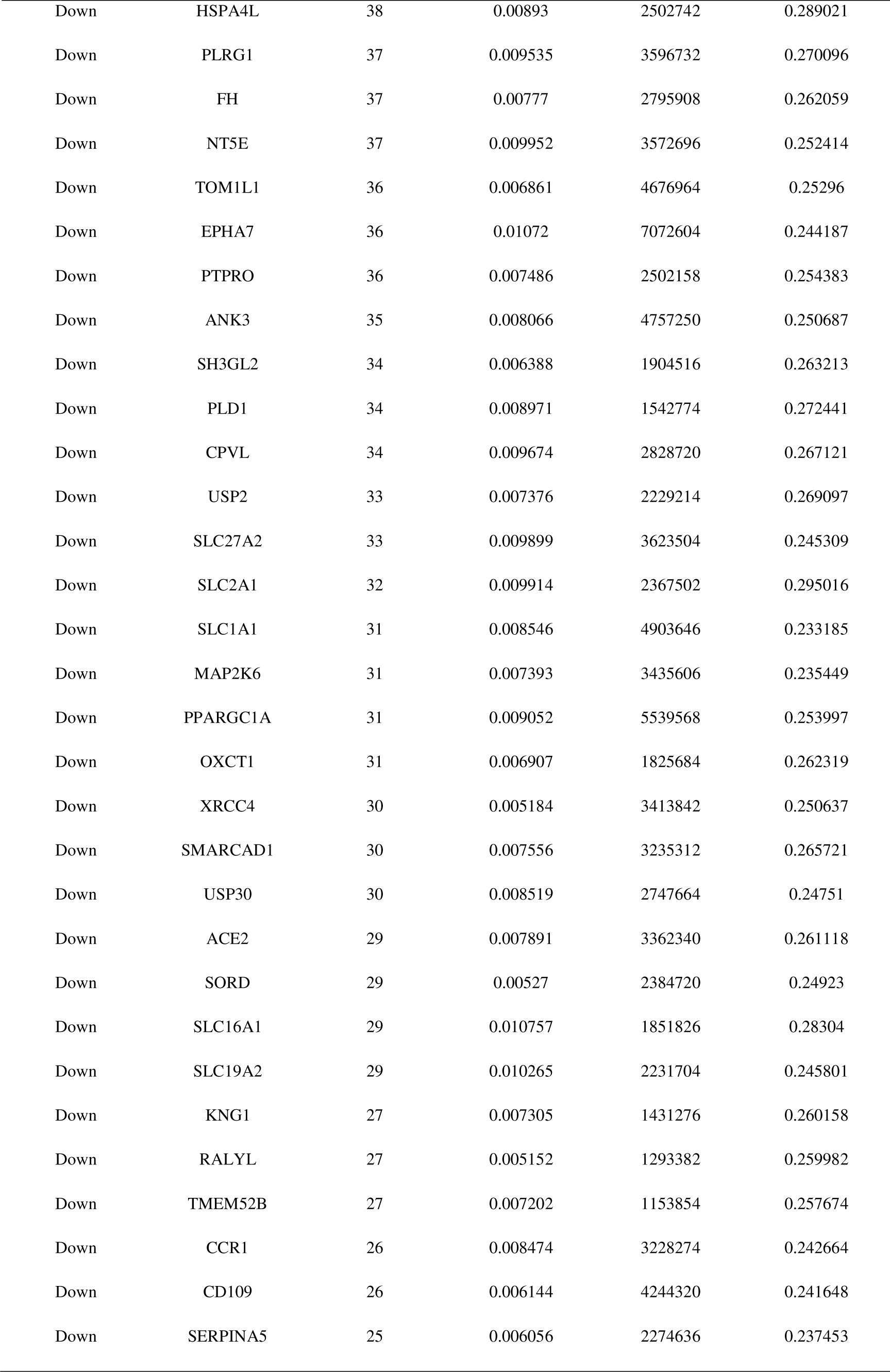
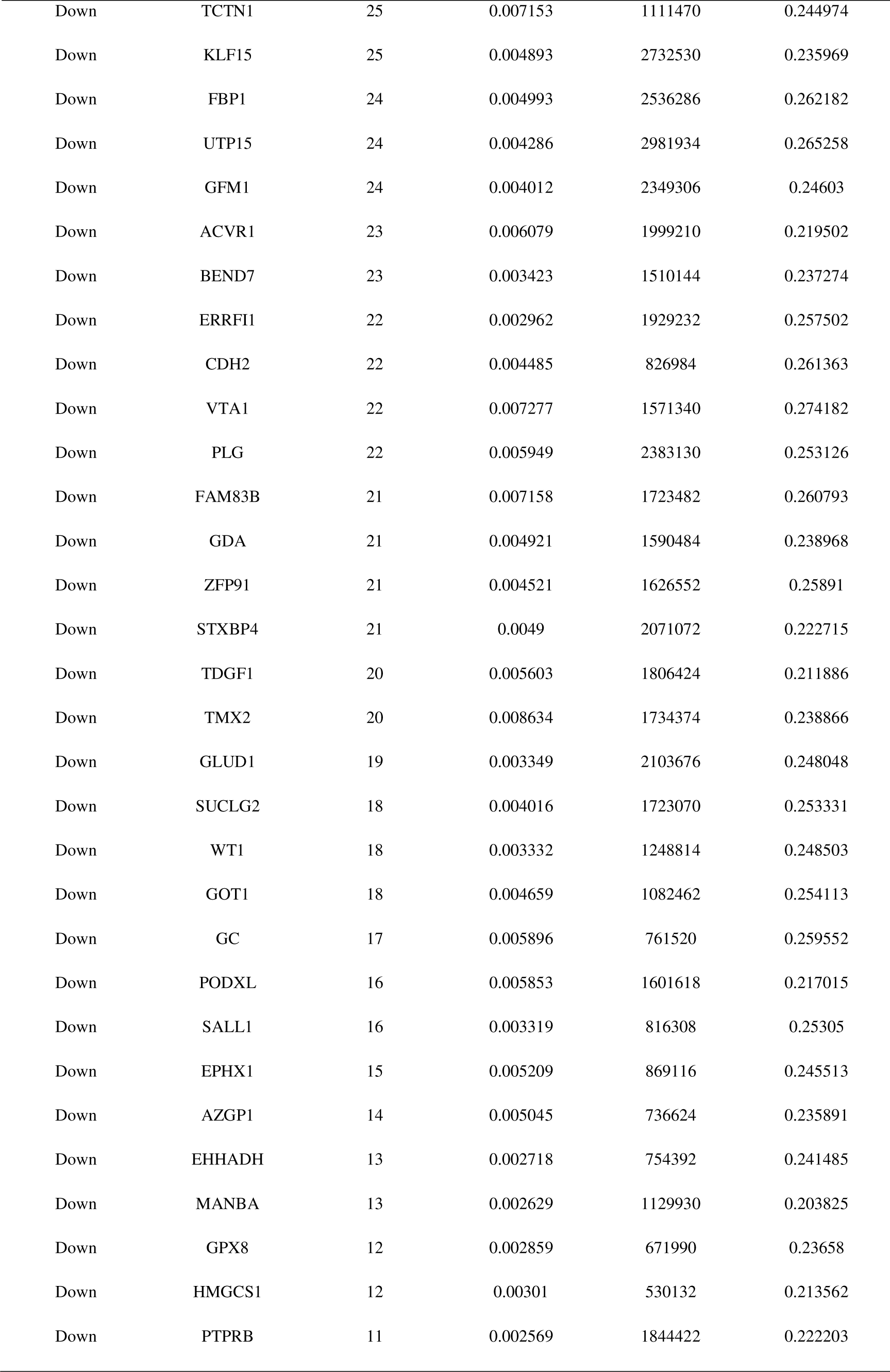
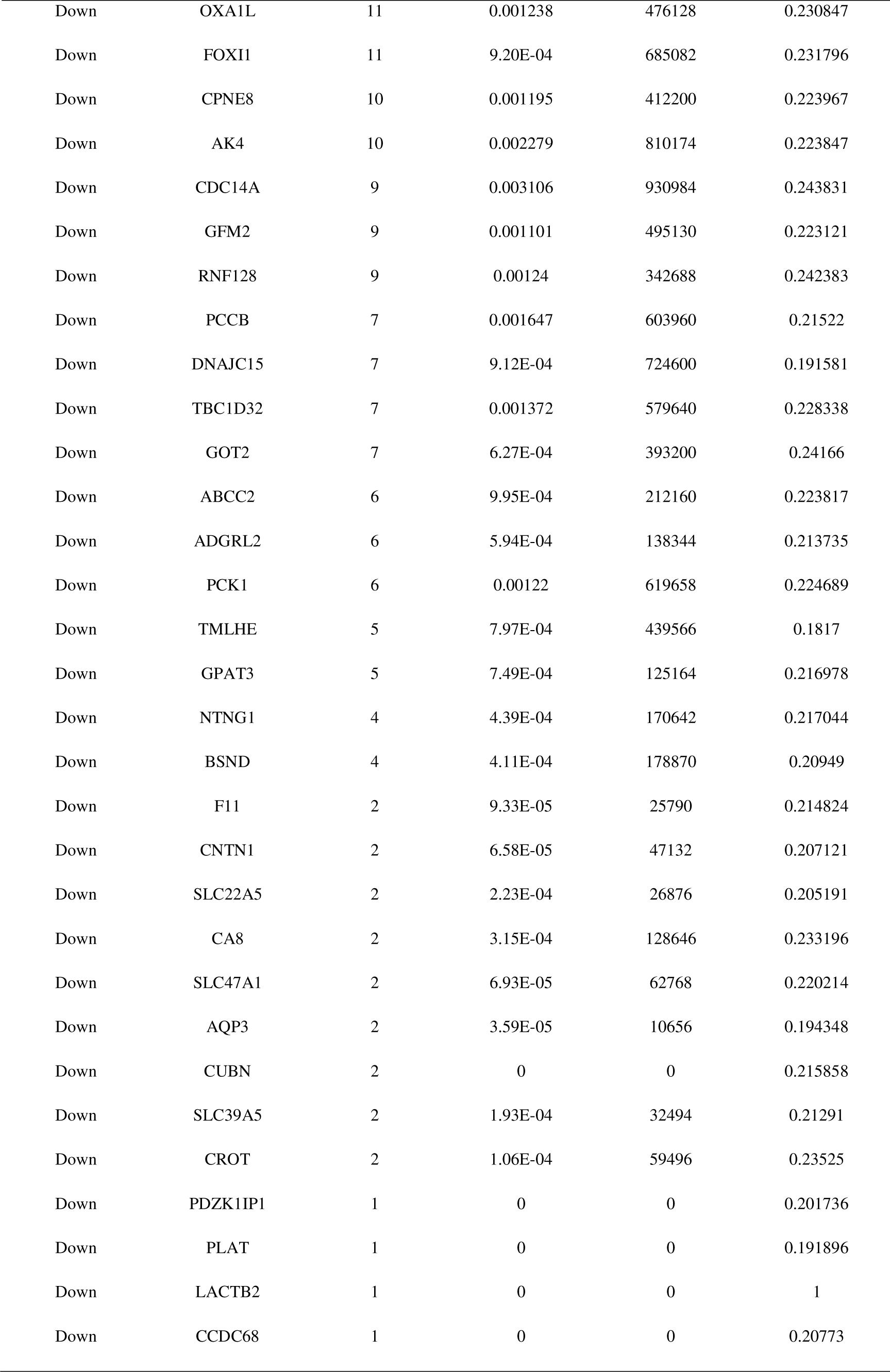
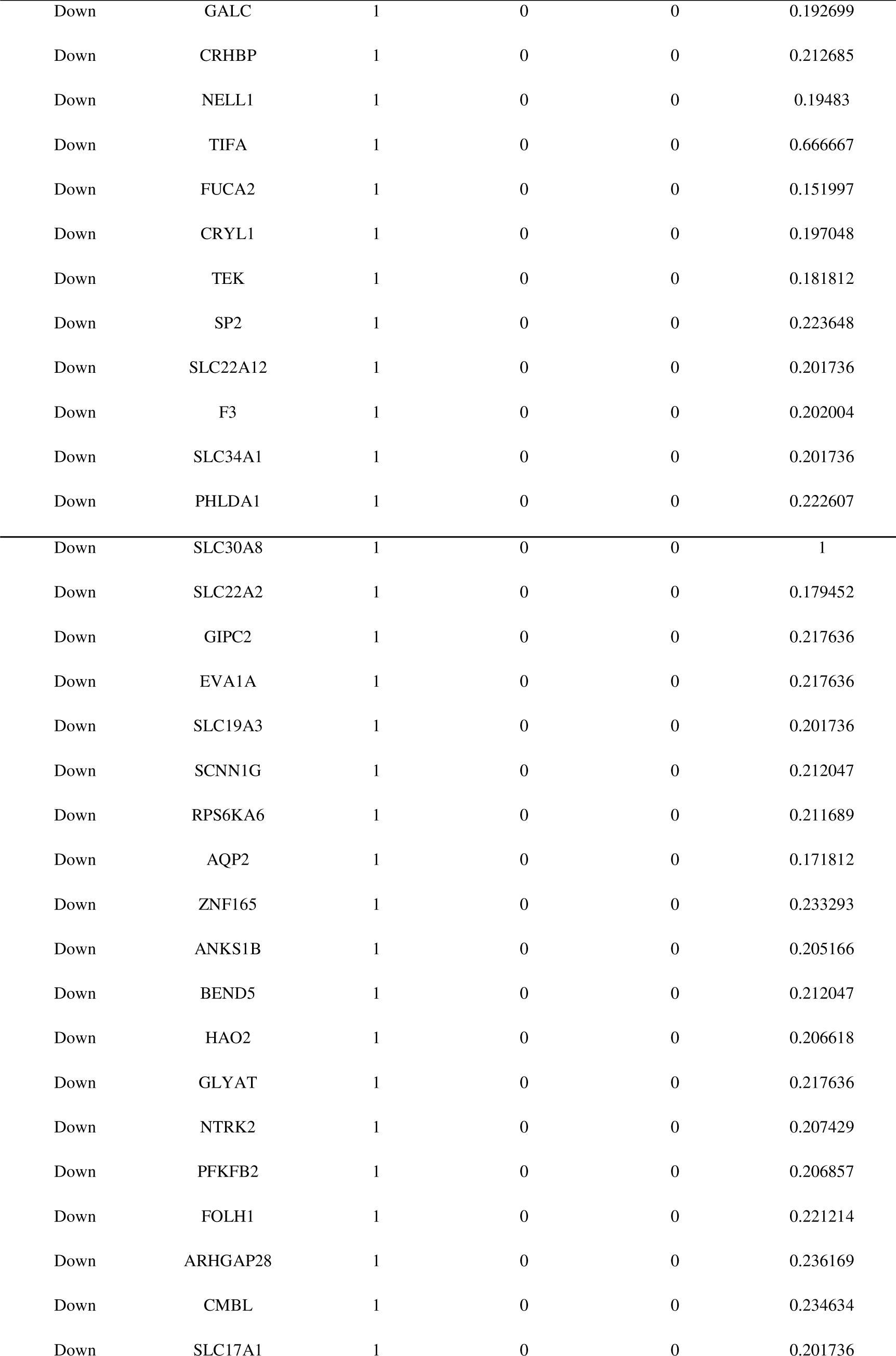
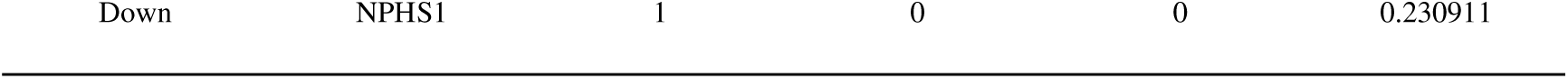
Topology table for up and down regulated genes.

### Construction of the miRNA-hub gene regulatory network

For the hub genes we identified, a miRNA-hub gene regulatory network was constructed including 2298 (miRNA: 2012; Hub Gene: 286) nodes and 10904 edges (Fig. 5). While PPP1CC was found to be regulated by 173 miRNAs (ex; hsa-mir-510-3p), RANGAP1 was found to be regulated by 152 miRNAs (ex; hsa-mir-7847-3p), NAP1L1 was found to be regulated by 145 miRNAs (ex; hsa-mir-202-3p), E2F1 was found to be regulated by 121 miRNAs (ex; hsa-mir-6837-5p), BMI1 was found to be regulated by 115 miRNAs (ex; hsa-mir-548ap-5p), VCAM1 was found to be regulated by 102 miRNAs (ex; hsa-mir-6086), CDH1 was found to be regulated by 75 miRNAs (ex; hsa-mir-638), MRPS30 was found to be regulated by 60 miRNAs (ex; hsa-mir-1246), MME was found to be regulated by 54 miRNAs (ex; hsa-mir-526a) and KIT was found to be regulated by 48 miRNAs (ex; hsa-mir-221) (Table 5).

**Fig. 5.**
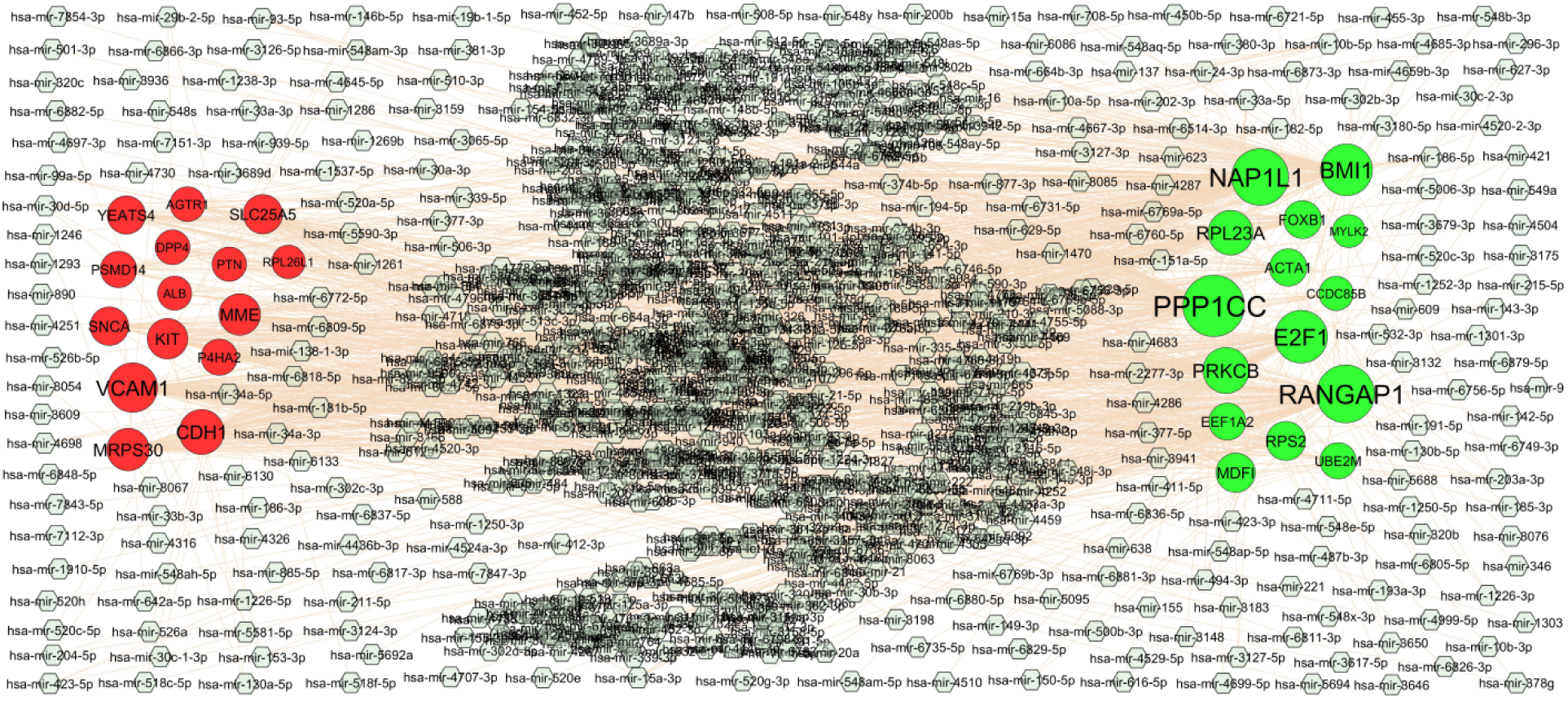
Hub gene - miRNA regulatory network. The ash color diamond nodes represent the key miRNAs; up regulated genes are marked in green; down regulated genes are marked in red.

**Table 5.** MiRNA - hub gene and TF – hub gene topology table.

| Regulation | Hub Genes | Degree | MicroRNA | Regulation | Hub Genes | Degree | TF |
| --- | --- | --- | --- | --- | --- | --- | --- |
| Up | PPP1CC | 173 | hsa-mir-510-3p | Up | E2F1 | 21 | MAX |
| Up | RANGAP1 | 152 | hsa-mir-7847-3p | Up | RANGAP1 | 12 | POU2F2 |
| Up | NAP1L1 | 145 | hsa-mir-202-3p | Up | EEF1A2 | 11 | TFAP2A |
| Up | E2F1 | 121 | hsa-mir-6837-5p | Up | FOXB1 | 9 | MEF2A |
| Up | BMI1 | 115 | hsa-mir-548ap-5p | Up | ACTA1 | 7 | FOXC1 |
| Up | PRKCB | 81 | hsa-mir-4999-5p | Up | UBE2M | 6 | SREBF1 |
| Up | RPL23A | 73 | hsa-mir-33a-3p | Up | BMI1 | 6 | EGR1 |
| Up | MDFI | 41 | hsa-mir-204-5p | Up | MDFI | 6 | EN1 |
| Up | RPS2 | 40 | hsa-mir-423-5p | Up | NAP1L1 | 5 | HOXA5 |
| Up | FOXB1 | 32 | hsa-mir-181b-5p | Up | MYLK2 | 5 | GATA2 |
| Up | ACTA1 | 30 | hsa-mir-5688 | Up | PRKCB | 4 | ZNF354C |
| Up | EEF1A2 | 27 | hsa-mir-412-3p | Up | RPL23A | 4 | JUN |
| Up | UBE2M | 23 | hsa-mir-151a-5p | Up | PPP1CC | 4 | SRF |
| Up | CCDC85B | 11 | hsa-mir-34a-5p | Up | RPS2 | 4 | SOX17 |
| Up | MYLK2 | 3 | hsa-mir-24-3p | Up | CCDC85B | 3 | ELK1 |
| Down | VCAM1 | 102 | hsa-mir-6086 | Down | MRPS30 | 15 | PAX2 |
| Down | CDH1 | 75 | hsa-mir-638 | Down | AGTR1 | 13 | USF2 |
| Down | MRPS30 | 60 | hsa-mir-1246 | Down | SNCA | 11 | YY1 |
| Down | MME | 54 | hsa-mir-526a | Down | MME | 9 | STAT3 |
| Down | KIT | 48 | hsa-mir-221 | Down | VCAM1 | 9 | PDX1 |
| Down | SLC25A5 | 41 | hsa-mir-10a-5p | Down | P4HA2 | 8 | BRCA1 |
| Down | SNCA | 37 | hsa-mir-153-3p | Down | KIT | 8 | ARID3A |
| Down | YEATS4 | 35 | hsa-mir-6811-3p | Down | SLC25A5 | 8 | NFIC |
| Down | PSMD14 | 24 | hsa-mir-186-5p | Down | DPP4 | 8 | RELA |
| Down | P4HA2 | 23 | hsa-mir-494-3p | Down | CDH1 | 7 | CEBPB |
| Down | AGTR1 | 15 | hsa-mir-182-5p | Down | YEATS4 | 7 | FOXL1 |
| Down | DPP4 | 14 | hsa-mir-1303 | Down | PTN | 6 | ELF5 |
| Down | PTN | 12 | hsa-mir-147b | Down | PSMD14 | 4 | NKX3-1 |
| Down | ALB | 10 | hsa-mir-29b-2-5p | Down | ALB | 4 | NFKB1 |
| Down | RPL26L1 | 7 | hsa-mir-421 | Down | RPL26L1 | 3 | FOXF2 |

### Construction of the TF-hub gene regulatory network

For the hub genes we identified, a TF-hub gene regulatory network was constructed including 375 (TF: 86; Hub Gene: 289) nodes and 2294 edges (Fig. 6). While E2F1 was found to be regulated by 21 TFs (ex; MAX), RANGAP1 was found to be regulated by 12 TFs (ex; POU2F2), EEF1A2 was found to be regulated by 11 TFs (ex; TFAP2A), FOXB1 was found to be regulated by 9 TFs (ex; MEF2A), ACTA1 was found to be regulated by 7 TFs (ex; FOXC1), MRPS30 was found to be regulated by 15 TFs (ex; PAX2), AGTR1 was found to be regulated by 13 TFs (ex; USF2), SNCA was found to be regulated by 11 TFs (ex; YY1), MME was found to be regulated by 9 TFs (ex; STAT3) and VCAM1 was found to be regulated by 9 TFs (ex; PDX1) (Table 5).

**Fig. 6.**
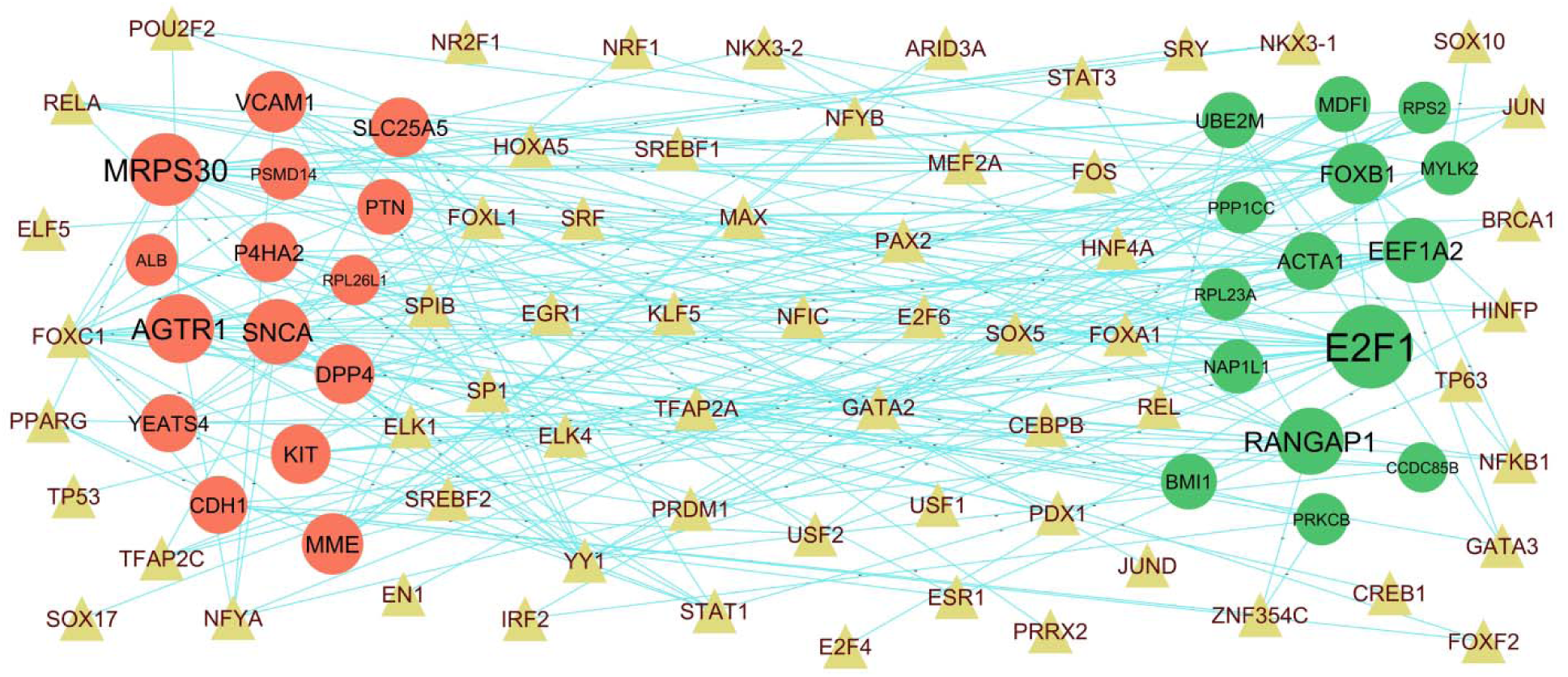
Hub gene - TF regulatory network. The olive color triangle nodes represent the key TFs; up regulated genes are marked in dark green; down regulated genes are marked in dark red.

### Receiver operating characteristic curve (ROC) analysis

To evaluate the discriminative potential of the identified hub genes, we generated receiver operating characteristic (ROC) curves for the ten hub genes we identified, as depicted in Fig. 7. Ten hub genes such as PPP1CC, RPS2, MDFI, BMI1, RPL23A, VCAM1, ALB, SNCA, DPP4 and RPL26L1 with AUC more than 0.80, indicating that they have the capability to diagnose AKI patients with excellent specificity and sensitivity.

**Fig. 7.**
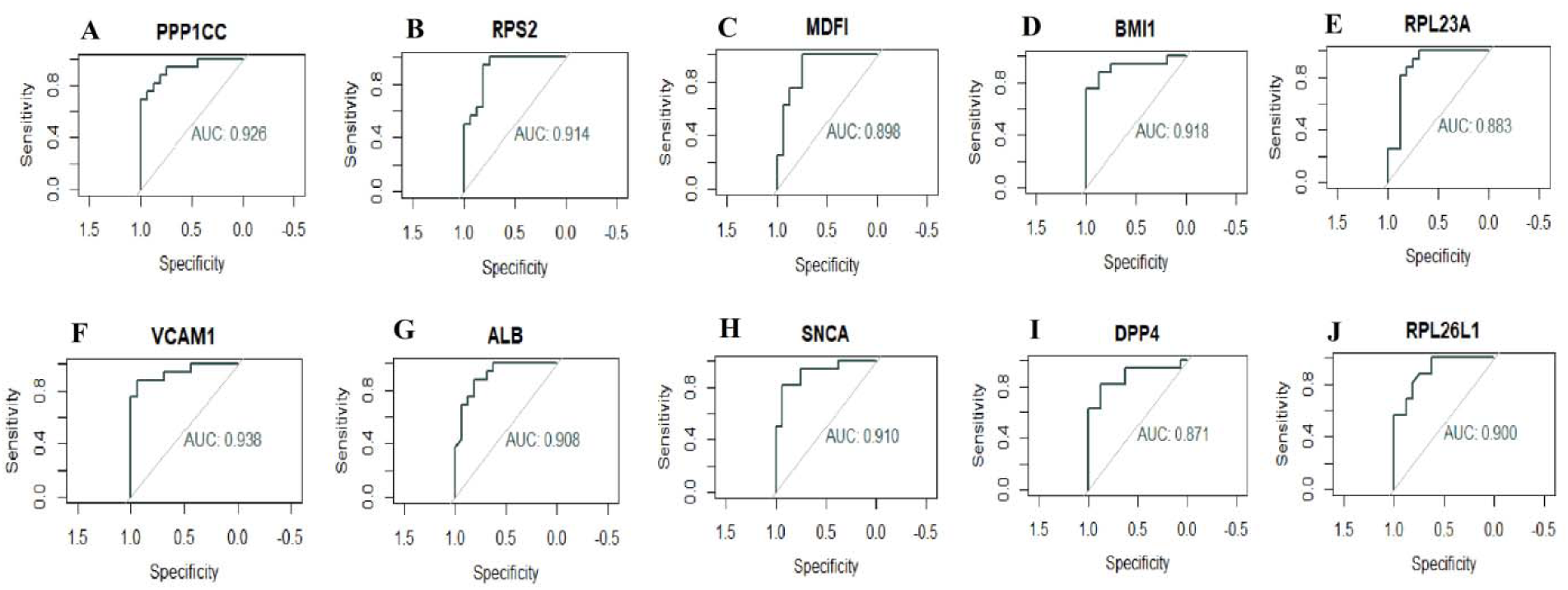
ROC curve analyses of hub genes. A) PPP1CC B) RPS2 C) MDFI D) BMI1 E) RPL23A F) VCAM1 G) ALB H) SNCA I) DPP4 J) RPL26L1

## Discussion

AKI is an acute renal failure with rapid loss of the kidney’s excretory function [51]. Timely diagnoses of AKI are vital for the management of such diseases, whereas such an early diagnostic tool is still lacking in clinical practice [52]. If not treated promptly and effectively, AKI can seriously reduce the quality of life. Therefore, sensitive and specific biomarkers of AKI are urgently needed to be detected. With the advancement of NGS technology, we now realize that genes play a key role in the diagnosis, occurrence, progression and treatment of AKI. Thus, many mRNAs, miRNAs and TFs are associated with AKI have been discovered and supported by a large number of studies.

In this investigation, we analyzed the AKI NGS data GSE139061 from the GEO database. It includes 39 AKI and 9 normal control samples, we found 956 DEGs (including 478 up regulated genes and 478 down regulated genes). Studies suggested that PLG (plasminogen) [53], AFM (afamin) [54] and UMOD (uromodulin) [55] are involved in advancement of AKI. PLG (plasminogen) [56], GSTA1 [57], AGTR1 [58], CYP4A11 [59], UMOD (uromodulin) [60] and IL1RL1 [61] might be a potential prognostic markers or therapeutic targets against the cardiovascular disorders. Previous studies have reported that the PLG (plasminogen) [62], GSTA1 [63], AGTR1 [64], CYP4A11 [65], UMOD (uromodulin) [66] and SLC12A1 [67] are a key regulators of hypertension. Recently, increasing evidence demonstrated that PLG (plasminogen) [68] was altered expressed in hypercholesterolemia. Studies had shown that PLG (plasminogen) [69], GSTA1 [70], AGTR1 [71] and CYP4A11 [72] were associated with liver disease. PLG (plasminogen) [73], AFM (afamin) [54], AGTR1 [74] and UMOD (uromodulin) [75] might be a potential therapeutic target for sepsis. Abnormal regulation of PLG (plasminogen) [76], AFM (afamin) [77], GSTA1 [78], AGTR1 [79] and UMOD (uromodulin) [80] are associated with diabetes mellitus. PLG (plasminogen) [81] plays an indispensable role in glomerulonephritis. AFM (afamin) [82], AGTR1 [83] and IL1RL1 [84] could be an early detection biomarkers for obesity. AGTR1 [85] and UMOD (uromodulin) [86] expression is a potential target for diabetic nephropathy. AGTR1 [87] and UMOD (uromodulin) [88] play an important role in chronic kidney disease. UMOD (uromodulin) [89] was observed to be associated with the risk of kidney fibrosis. These genes might also have implications in the influence of important genes on AKI. Although we have seen less investigation of these genes on AKI, we speculate from our current investigation that it might play an essential role in AKI; more experiments will investigate the molecular mechanism in the future.

For a more in-depth understanding of these DEGs, we analyzed the selected genes for GO and REACTOME pathway enrichment analyses and found that GO terms and signaling pathways were significant. Signaling pathways include neuronal system [90], GPCR ligand binding [91], metabolism [92] and metabolism of lipids [93] were responsible for progression of AKI. MYH7 [94], BMI1 [95], IGF2 [96], ADORA2A [97], KLK10 [98], MEIS2 [99], IRF7 [100], PRKCB (protein kinase C beta) [101], CCL5 [102], ADAM33 [103], EEF1A2 [104], ACTN3 [105], TRIM65 [106], FCN1 [107], CARNS1 [108], AMH (anti-Mullerian hormone) [109], E2F1 [110], PF4 [111], RPL3L [112], TRIM72 [113], HOXB4 [114], TP73 [115], KCNH2 [116], (advanced glycosylation end-product specific receptor) [117], SMPD3 [118], TYMP (thymidine phosphorylase) [119], RIPPLY3 [120], GREM1 [121], CYP11B2 [122], MYLK2 [123], WNT3A [124], MSX1 [125], COMP (cartilage oligomeric matrix protein) [126], FLI1 [127], ACTA1 [128], TCAP (titin-cap) [129], TUBB1 [130], TNNI3 [131], HSPB7 [132], DES (desmin) [133], RAP1B [134], TNNT1 [135], BHMT (betaine--homocysteine S-methyltransferase) [136], ANGPTL3 [137], CYP3A5 [138], KMO (kynurenine 3-monooxygenase) [139], HMGCS2 [140], AGXT2 [141], FABP1 [142], SLC22A12 [143], CUBN (cubilin) [144], MIOX (myo-inositol oxygenase) [145], FBP1 [146], ARG2 [147], FGF1 [148], CRY1 [149], PPARGC1A [150], UGT1A6 [151], ECHDC3 [152], CYP2C8 [153], ACOX2 [154], SLC2A9 [155], MSRA (methionine sulfoxidereductase A) [156], GC (GC vitamin D binding protein) [157], VNN1 [158], NOX4 [159], GOT2 [160], EPHX2 [161], SLC2A1 [162], AKR1C3 [163], BPGM (bisphosphoglyceratemutase) [164], NR4A3 [165], EPHX1 [166], PFKFB2 [167], CDH2 [168], SLC22A6 [169], F11 [170], AQP2 [171], TIMD4 [172], EGF (epidermal growth factor) [173], ANGPT1 [174], SLC26A4 [175], KL (klotho) [176], SLC5A10 [177], PDZK1 [178], EVA1A [179], ACE2 [180], NR4A1 [181], FOLH1 [182], NPNT (nephronectin) [183], PKP2 [184], SLC30A8 [185], KCNJ3 [186], LYVE1 [187], CADM1 [188], AZGP1 [189], C6 [190], ABCC2 [191], DPP4 [192], STC1 [193], REN (renin) [194], TRPM6 [195], ABCB1 [196], MTTP (microsomal triglyceride transfer protein) [197], PTN (pleiotrophin) [198], ERRFI1 [199], MSR1 [200], USP2 [201], KCNE4 [202], CA2 [203], PPM1L [204], SLC8A1 [205], CCR1 [206], PRLR (prolactin receptor) [207], NPHS1 [208], GPC6 [209], SELP (selectin P) [210], TDGF1 [211], MR1 [212], TNFRSF11B [213], TRPM3 [214], FZD5 [215], ERBB4 [216], F8 [217], GYG1 [218], PRCP (prolylcarboxypeptidase) [219], VCAM1 [220], PLD1 [221], SLCO4C1 [222], SEMA5A [223], NAT1 [224], PIAS1 [225], PLAT (plasminogen activator, tissue type) [226] and ALB (albumin) [227] can facilitate cardiovascular disorders. MYH7 [228], WNT3A [229], ANGPTL3 [230], CYP3A5 [231], FGF1 [232], HMGCS1 [233], NOX4 [234], AQP2 [235], EGF (epidermal growth factor) [236], NR4A1 [237], DPP4 [238], STC1 [239], ABCB1 [240], MTTP (microsomal triglyceride transfer protein) [241], F8 [242], VCAM1 [243] and ALB (albumin) [244] are participate in pathogenic processes of hypercholesterolemia. BMI1 [245], CCL5 [246], GREM1 [247], ANGPTL3 [248], ARG2 [249], MSRA (methionine sulfoxidereductase A) [250], SNCA (synuclein alpha) [251], NOX4 [252], PFKFB2 [253], PDZK1 [254], SUCNR1 [255], LYVE1 [256], AZGP1 [257], ERBB4 [258] and PLAT (plasminogen activator, tissue type) [259] might serve as molecular markers for kidney fibrosis. BMI1 [260], IGF2 [261], IRF7 [262], CCL5 [263], ACTN3 [264], E2F1 [265], PF4 [266], TEAD4 [267], TBX4 [268], GREM1 [269], CYP11B2 [270], WNT3A [124], COMP (cartilage oligomeric matrix protein) [271], FLI1 [272], RAP1B [273], ANGPTL3 [274], CYP3A5 [275], HSD11B2 [276], HMGCS2 [277], AGXT2 [278], SLC22A12 [279], FGF1 [280], CRY1 [281], PPARGC1A [282], SLC19A3 [283], CYP2C8 [284], ACOX2 [285], SLC2A9 [286], MSRA (methionine sulfoxidereductase A) [287], VNN1 [288], EPHX2 [289], CROT (carnitine O-octanoyltransferase) [290], SCNN1B [291], NR4A3 [292], HSD17B7 [293], SLC22A2 [294], AQP2 [295], SLC2A2 [296], EGF (epidermal growth factor) [297], ANGPT1 [298], SLC26A4 [299], KL (klotho) [300], SCNN1G [301], PDZK1 [302], PTPRD (protein tyrosine phosphatase receptor type D) [303], ACE2 [304], FOLH1 [305], SUCNR1 [306], GLCE (glucuronic acid epimerase) [307], AQP3 [308], DPP4 [309], REN (renin) [310], TRPM6 [311], ABCB1 [312], MTTP (microsomal triglyceride transfer protein) [313], CALCRL (calcitonin receptor like receptor) [314], ENPEP (glutamylaminopeptidase) [315], USP2 [316], SLC8A1 [317], SELP (selectin P) [318], F8 [319], DIO2 [320], PRCP (prolylcarboxypeptidase) [219], VCAM1 [321], NEDD9 [322], SLCO4C1 [323], NTRK2 [324]. PLAT (plasminogen activator, tissue type) [325] and ALB (albumin) [326] might be involved in the occurrence and development of hypertension. BMI1 [327], AMH (anti-Mullerian hormone) [328], E2F1 [329], PF4 [330], VASH2 [331], GRIN1 [332], CYP11B2 [333], COMP (cartilage oligomeric matrix protein) [334], DES (desmin) [335], ANGPTL3 [336], CYP3A5 [337], DPEP1 [338], LRP2 [339], AGXT2 [340], FABP1 [341], SLC22A12 [342], CUBN (cubilin) [343], MIOX (myo-inositol oxygenase) [344], ARG2 [345], KHK (ketohexokinase) [346], CYP2C8 [347], SLC2A9 [348], GC (GC vitamin D binding protein) [349], VNN1 [350], NOX4 [351], EPHX2 [352], AKR1C3 [353], NR4A3 [354], PFKFB2 [355], SLC22A6 [356], F11 [357], SLC22A2 [358], AQP2 [171], EGF (epidermal growth factor) [359], KNG1 [360], SERPINA5 [361], KL (klotho) [362], ACE2 [363], NPNT (nephronectin) [364], SLC47A1 [358], MGAM (maltase-glucoamylase) [365], AQP3 [366], AZGP1 [367], GALNT3 [368], DPP4 [369], STC1 [370], ABCB1 [371], ERRFI1 [372], TREH (trehalase) [373], MANBA (mannosidase beta) [374], ERBB4 [375], VCAM1 [376] and ALB (albumin) [377] might play an important role in the pathophysiology of AKI. BMI1 [378], IGF2 [379], PRKCB (protein kinase C beta) [380], CCL5 [381], E2F1 [382], PF4 [111], CYP11B2 [383], WNT3A [384], COMP (cartilage oligomeric matrix protein) [385], DES (desmin) [335], ANGPTL3 [386], CYP3A5 [387], LRP2 [388], PAH (phenylalanine hydroxylase) [389], HMGCS2 [390], AGXT2 [391], FABP1 [392], CUBN (cubilin) [393], DIO1 [394], MIOX (myo-inositol oxygenase) [344], FGF1 [395], CRY1 [396], PPARGC1A [397], CYP2C8 [398], GC (GC vitamin D binding protein) [399], VNN1 [350], NOX4 [400], EPHX2 [401], F11 [402], AQP2 [403], NAT8 [404], CLDN2 [405], EGF (epidermal growth factor) [406], ANGPT1 [174], CLCN5 [407], KL (klotho) [176], DEFB1 [408], ACE2 [409], NR4A1 [410], SLC13A3 [411], CADM1 [412], AZGP1 [367], DPP4 [413], REN (renin) [414], ABCB1 [415], CCR1 [416], MANBA (mannosidase beta) [417], SELP (selectin P) [418], CLDN16 [419], F8 [420], VCAM1 [421], PLD1 [422], SLCO4C1 [423], TINAG (tubulointerstitial nephritis antigen) [424], SULT1C2 [425], PLAT (plasminogen activator, tissue type) [426] and ALB (albumin) [427] can participate in the occurrence and development of chronic kidney disease. IGF2 [428], SCARNA10 [429], IRF7 [430], PRKCB (protein kinase C beta) [431], CCL5 [432], E2F1 [433], PF4 [434], HOXC8 [435], SMPD3 [118], GREM1 [436], WNT3A [437], COMP (cartilage oligomeric matrix protein) [438], FLI1 [439], DES (desmin) [440], BHMT (betaine--homocysteine S-methyltransferase) [441], ANGPTL3 [442], PCK1 [443], LRP2 [444], HMGCS2 [445], AGXT2 [446], FABP1 [447], DIO1 [448], CYP8B1 [449], FGF1 [450], PPARGC1A [451], PNPLA8 [452], ACOX2 [453], FH (fumaratehydratase) [454], MSRA (methionine sulfoxidereductase A) [455], GC (GC vitamin D binding protein) [456], ECHDC1 [457], NOX4 [458], ABHD6 [459], GOT2 [460], SLC2A1 [461], HSD17B7 [462], F11 [463], AQP2 [464], HRG (histidine rich glycoprotein) [465], KL (klotho) [466], EVA1A [467], ACE2 [468], NR4A1 [469], SUCNR1 [470], AQP3 [471], AZGP1 [472], ABCC2 [473], STC1 [474], MTTP (microsomal triglyceride transfer protein) [475], CALCRL (calcitonin receptor like receptor) [476], MSR1 [477], KCNE4 [478], CA2 [479], FRK (fyn related Src family tyrosine kinase) [480], CCR1 [481], PRLR (prolactin receptor) [482], F8 [483], DIO2 [484], VCAM1 [485], PLD1 [486], AOX1 [487], GFM1 [488], PIAS1 [489], P4HA2 [490], PLAT (plasminogen activator, tissue type) [491], ALB (albumin) [492] and PNPLA8 [493] are significantly related to the progression of liver diseases. IGF2 [494], IRF7 [495], E2F1 [496], TEAD4 [497], KCNH2 [498], E2F2 [499], SNHG7 [500], FLI1 [501], CYP3A5 [502], HMGCS2 [503], GOT1 [504], PPARGC1A [505], GC (GC vitamin D binding protein) [506], VNN1 [507], NOX4 [508], SLC2A1 [509], BPGM (bisphosphoglyceratemutase) [164], NR4A3 [354], PFKFB2 [510], CDH2 [511], F11 [512], AQP2 [513], CLDN2 [514], EGF (epidermal growth factor) [515], ANGPT1 [516], KNG1 [517], SERPINA5 [518], HRG (histidine rich glycoprotein) [519], KL (klotho) [520], DEFB1 [521], ACE2 [522], AQP3 [523], CADM1 [188], DPP4 [524], STC1 [525], REN (renin) [526], TRPM6 [527], MSR1 [528], CCR1 [529], TNFRSF11B [530], FZD5 [531], ERBB4 [216], F8 [532], VCAM1 [533], PTGER3 [534] and ALB (albumin) [535] have been shown to influence the genetic risk of sepsis. IGF2 [536], IRF7 [100], PRKCB (protein kinase C beta) [101], CCL5 [537], EEF1A2 [538], ACTN3 [264], FCN1 [539], BRSK2 [540], MNX1 [541], AMH (anti-Mullerian hormone) [542], E2F1 [543], HAP1 [544], PF4 [545], AGER (advanced glycosylation end-product specific receptor) [546], E2F2 [547], TYMP (thymidine phosphorylase) [548], PPP1CC [549], NR2E1 [550], GREM1 [436], GRIN1 [551], WNT3A [552], COMP (cartilage oligomeric matrix protein) [553], BHMT (betaine--homocysteine S-methyltransferase) [554], ANGPTL3 [555], PCK1 [556], KMO (kynurenine 3-monooxygenase) [557], HSD11B2 [558], UGT1A9 [559], HMGCS2 [140], AGXT2 [560], FABP1 [142], CUBN (cubilin) [561], SLC17A1 [562], MIOX (myo-inositol oxygenase) [563], SORD (sorbitol dehydrogenase) [564], FGF1 [565], CRY1 [566], PPARGC1A [451], FUT6 [567], ECHDC3 [568], SLC19A3 [569], KHK (ketohexokinase) [346], CYP2C8 [570], FH (fumaratehydratase) [571], SLC2A9 [572], GC (GC vitamin D binding protein) [573], SNCA (synuclein alpha) [574], VNN1 [575], NOX4 [576], ABHD6 [459], EPHX2 [577], SLC2A1 [162], SLC19A2 [578], EPHX1 [579], SRR (serine racemase) [580], F11 [581], SLC22A2 [582], AQP2 [583], CLDN8 [584], SLC2A2 [585], CLDN2 [586], EGF (epidermal growth factor) [587], SLC38A4 [588], PIK3C2G [589], KL (klotho) [590], DEFB1 [591], F5 [592], KCNH6 [593], PTPRD (protein tyrosine phosphatase receptor type D) [594], TSPAN7 [595], ACE2 [363], NR4A1 [596], KCNJ10 [597], SUCNR1 [306], KCNJ1 [598], SLC47A1 [599], SLC30A8 [185], SLC22A5 [600], LYVE1 [601], AQP3 [602], CADM1 [603], AZGP1 [604], C6 [605], GALNT3 [606], DPP4 [607], REN (renin) [608], PODXL (podocalyxin like) [609], TRPM6 [610], ABCB1 [611], MTTP (microsomal triglyceride transfer protein) [612], MSR1 [613], HEPH (hephaestin) [614], KCNJ15 [615], TREH (trehalase) [616], PRLR (prolactin receptor) [617], SELP (selectin P) [618], TNFRSF11B [619], TNFRSF21 [620], FZD5 [215], DNER (delta/notch like EGF repeat containing) [621], CDH1 [622], F8 [623], DIO2 [624], PRCP (prolylcarboxypeptidase) [625], VCAM1 [626], DNAJC15 [627], PLD1 [628], NAT1 [629], PIAS1 [225], PLAT (plasminogen activator, tissue type) [630] and ALB (albumin) [631] are considered to be a biomarkers and therapeutic targets for diabetes mellitus. IGF2 [632], RPPH1 [633], PRKCB (protein kinase C beta) [634], CCL5 [635], VASH2 [636], GREM1 [637], CYP11B2 [638], HIC1 [639], PCK1 [640], HSD11B2 [641], MME (membrane metalloendopeptidase) [642], FABP1 [142], MIOX (myo-inositol oxygenase) [643], ARG2 [644], PPARGC1A [645], VNN1 [646], NOX4 [647], EPHX2 [577], DDIT4 [648], SLC2A1 [649], PFKFB2 [650], CDH2 [651], SLC22A2 [294], AQP2 [652], ANGPT1 [653], KL (klotho) [654], ACE2 [655], STC1 [656], REN (renin) [608], ERRFI1 [657], ERBB4 [658], NTNG1 [659], VCAM1 [660], PTGER3 [661] and BBOX1 [662] expression levels are associated with diabetic nephropathy. IGF2 [663], IRF7 [664], PRKCB (protein kinase C beta) [665], CCL5 [666], ACTN3 [667], AMH (anti-Mullerian hormone) [668], E2F1 [669], UBE2M [670], TP73 [671], AGER (advanced glycosylation end-product specific receptor) [672], SMPD3 [118], NR2E1 [673], ANGPTL3 [674], CYP3A5 [675], PCK1 [676], LRP2 [677], FABP1 [678], SLC22A12 [279], CYP8B1 [679], MIOX (myo-inositol oxygenase) [680], ARG2 [681], FGF1 [450], CRY1 [682], PPARGC1A [451], CRYM (crystallin mu) [683], SLC19A3 [283], FH (fumaratehydratase) [684], SLC2A9 [286], GC (GC vitamin D binding protein) [685], RGN (regucalcin) [686], MMAA (metabolism of cobalamin associated A) [687], NOX4 [688], ABHD6 [689], EPHX2 [690], DDIT4 [691], EGF (epidermal growth factor) [173], DEFB1 [692], ACE2 [693], PTPRO (protein tyrosine phosphatase receptor type O) [694], SUCNR1 [695], NPNT (nephronectin) [696], PKP2 [697], LYVE1 [601], GPAT3 [698], AZGP1 [604], DPP4 [699], REN (renin) [700], ABCB1 [701], MTTP (microsomal triglyceride transfer protein) [702], MSR1 [613], CLIC5 [703], USP2 [704], FRK (fyn related Src family tyrosine kinase) [705], CCR1 [706], PRLR (prolactin receptor) [707], SELP (selectin P) [618], ERBB4 [708], DIO2 [709], PRCP (prolylcarboxypeptidase) [625], VCAM1 [710], DNAJC15 [627], NTRK2 [711], STK33 [712], MAP2K6 [713], PIAS1 [714] and ALB (albumin) [715] have been proposed as novel biomarkers for obesity. ADORA2A [716], DPEP1 [717], KMO (kynurenine 3-monooxygenase) [718], E2F1 [719], VASH2 [720], ARG2 [721], MSRA (methionine sulfoxidereductase A) [722], NOX4 [723], AQP2 [724], EGF (epidermal growth factor) [725], ANGPT1 [726], KL (klotho) [727], ACE2 [728], NR4A1 [729], SLC47A1 [358], LYVE1 [730], AQP3 [731], DPP4 [732], STC1 [733], TREH (trehalase) [734], CCR1 [735], VCAM1 [736], PLAT (plasminogen activator, tissue type) [737] and ALB (albumin) [738] might serve as potential biomarkers for renal ischemia. GLRX5 [739], AMH (anti-Mullerian hormone) [740], HOXB4 [741], COMP (cartilage oligomeric matrix protein) [742], FLI1 [743], PCK1 [744], CUBN (cubilin) [745], CYP2C8 [746], BPGM (bisphosphoglyceratemutase) [747], SLC19A2 [578], HRG (histidine rich glycoprotein) [748], KL (klotho) [749], REN (renin) [414], ABCB1 [750], F8 [751], VCAM1 [752], ACVR1 [753] and PLAT (plasminogen activator, tissue type) [754] might be regarded as a valuable biomarker for diagnosis, treatment and prognosis of anemia. Altered expression of BRSK1 [755], SCNN1G [756], CADM1 [757], PODXL (podocalyxin like) [758], NPHS1 [759], CNTN1 [760], NTNG1 [761], PLAT (plasminogen activator, tissue type) [259] and P3H2 [762] are involved in the development and progression of membranous nephropathy. Previous studies have demonstrated that CYP11B2 [763], FLI1 [764], SLC7A7 [765], NPNT (nephronectin) [766], AZGP1 [767], TREH (trehalase) [768], CCR1 [769], CNTN1 [770], VCAM1 [771] and GBP3 [772] are linked with the development mechanisms of glomerulonephritis. A recent study revealed that FABP1 [773] and NPHS1 [774] expression was altered in IgA nephropathy. CLCN5 [775], ENPEP (glutamylaminopeptidase) [776], CCR1 [777] and NPHS1 [778] have been identified in focal segmental glomerulosclerosis. A study indicates that ALB (albumin) [779] is positively correlated with hypoalbuminemia. We gave a new confirmation for that these enriched genes are expected to become a biomarkers for AKI prognosis.

DEGs genes were used to construct a PPI network to understand the mutual biological function of proteins and predict therapeutic targets. Identification of hub genes that might be key therapeutic targets or biomarkers for AKI were associated with numerious molecular pathogeneses. Based on the PPI network constructed by the online database HIPIE, we identified hub genes. PPP1CC [549], VCAM1 [626], ALB (albumin) [631], SNCA (synuclein alpha) [574] and DPP4 [607] are mainly associated with diabetes mellitus. BMI1 [95], VCAM1 [220], ALB (albumin) [227] and DPP4 [192] plays a role in diagnosis of cardiovascular disorders. BMI1 [245] and SNCA (synuclein alpha) [251] have been known to be involved in kidney fibrosis progression. BMI1 [260], VCAM1 [321], ALB (albumin) [326] and DPP4 [309] have a significant prognostic potential in hypertension. BMI1 [327], VCAM1 [376], ALB (albumin) [377] and DPP4 [369] are molecular markers for the diagnosis and prognosis of AKI. BMI1 [378], VCAM1 [421], ALB (albumin) [427] and DPP4 [413] have emerged as a potential targets for chronic kidney disease therapy. VCAM1 [243], ALB (albumin) [244] and DPP4 [238] genes are a potential markers for the detection and prognosis of hypercholesterolemia. A previous study reported that VCAM1 [485] and ALB (albumin) [492] are altered expressed in liver diseases. Studies have also shown that VCAM1 [533], ALB (albumin) [535] and DPP4 [524] are an important biomarkers of sepsis. VCAM1 [660] has been found to influence diabetic nephropathy progression. Study demonstrated that VCAM1 [710], ALB (albumin) [715] and DPP4 [699] were essential for the induction of obesity. VCAM1 [736], ALB (albumin) [738] and DPP4 [732] are potential therapeutic targets in renal ischemia. VCAM1 [752] expression was associated with the severity of anemia. VCAM1 [771] has been reported involved in the glomerulonephritis. ALB (albumin) [779] was identified as a candidate causal gene of hypoalbuminemia. Novel biomarkers include RPS2, MDFI (MyoD family inhibitor), RPL23A, RPL26L1, PCGF6, MRPS30, MRPL18 and MRPL47 might play a role in the development of AKI. This result suggests that these hub genes are closely related to the development of AKI.

To elucidate the potential molecular mechanism of the hub genes in AKI, we conducted a miRNA-hub gene regulatory network and TF-hub gene regulatory network analysis. The identified hub gens, TFs and miRNAs might be involved in the pathological process of AKI. PPP1CC [549], E2F1 [543], EEF1A2 [538], VCAM1 [626], CDH1 [622], AGTR1 [79], SNCA (synuclein alpha) [574], hsa-mir-221 [780], YY1 [781], STAT3 [782] and PDX1 [783] are closely involved with diabetes mellitus. E2F1 [110], BMI1 [95], EEF1A2 [104], ACTA1 [128], VCAM1 [220], AGTR1 [58], hsa-mir-221 [784], MEF2A [785], FOXC1 [786], YY1 [787], STAT3 [788] and BRCA1 [789] were significantly associated with cardiovascular disorders. E2F1 [265], BMI1 [260], VCAM1 [321], AGTR1 [64], hsa-mir-638 [790], hsa-mir-221 [791], FOXC1 [792], YY1 [793], STAT3 [794] and BRCA1 [795] provided a clear picture of the prognosis of patients with hypertension. E2F1 [329], BMI1 [327], VCAM1 [376], USF2 [796], YY1 [797], STAT3 [798] and BRCA1 [799] have always been found in AKI. Identified E2F1 [382], BMI1 [378], VCAM1 [421], AGTR1 [87], MEF2A [800] and STAT3 [798] as a biomarkers for chronic kidney disease. E2F1 [433], VCAM1 [485], AGTR1 [71], hsa-mir-221 [801], MEF2A [802] and YY1 [803] are associated with developing liver diseases. E2F1 [496], AGTR1 [74], VCAM1 [533], hsa-mir-7847-3p [804], FOXC1 [805], USF2 [796], YY1 [806], STAT3 [807] and BRCA1 [799] have been observed in sepsis patients. Studies have shown that E2F1 [669], VCAM1 [715], AGTR1 [83], hsa-mir-221 [780], YY1 [808], STAT3 [809], PDX1 [810] and BRCA1 [811] are associated with obesity. E2F1 [719] and VCAM1 [736] are altered expressed in patients with renal ischemia. BMI1 [245], SNCA (synuclein alpha) [251] and YY1 [812] were an important target biomarker of kidney fibrosis. VCAM1 [243] was related to the early-onset of hypercholesterolemia. The expression pattern of VCAM1 [660], MME (membrane metalloendopeptidase) [642], AGTR1 [85], hsa-mir-221 [813], YY1 [814] and PDX1 [815] might offer useful information for treating diabetic nephropathy. VCAM1 [752], STAT3 [816] and BRCA1 [817] have been used as an independent biomarker to predict prognosis in patients with anemia. VCAM1 [771], USF2 [818] and YY1 [819] can be used as a diagnostic marker for glomerulonephritis. Novel biomarkers associated with diagnosis of AKI were identified in this investigation: RANGAP1, NAP1L1, FOXB1, MRPS30, KIT (KIT proto-oncogene, receptor tyrosine kinase), hsa-mir-510-3p, hsa-mir-202-3p, hsa-mir-6837-5p, hsa-mir-548ap-5p, hsa-mir-6086, hsa-mir-1246, hsa-mir-526a, MAX (MYC associated factor X), POU2F2 and TFAP2A. We gave a new confirmation for that hub genes, miRNA and TFs are expected to become a biomarkers for AKI prognosis.

## Conclusions

This investigation identified significant DEGs between AKI and normal control samples via analyzing NGS dataset. PPP1CC, RPS2, MDFI, BMI1, RPL23A, VCAM1, ALB, SNCA, DPP4 and RPL26L1 were verified and considered as hub genes were associated with disease prognosis, which could be predictive and therapeutic targets. The above results provide new research directions for the relationship between AKI and its associated complications.

## Acknowledgement

I thanks very much to Michael T Eadon, Indiana University, Indianapolis, Indiana, USA, the author who deposited their NGS dataset GSE139061, into the public GEO database.

## Ethical approval

This article does not contain any studies with human participants or animals performed by any of the authors.

## Informed consent

No informed consent because this study does not contain human or animals participants.

## Availability of data and materials

The datasets supporting the conclusions of this article are available in the GEO (Gene Expression Omnibus) (https://www.ncbi.nlm.nih.gov/geo/) repository. [(GSE139061) https://www.ncbi.nlm.nih.gov/geo/query/acc.cgi?acc=GSE139061]

## Consent for publication

Not applicable.

## Author Contributions

B. V. - Writing original draft, and review and editing

C. V. - Software and investigation

